# Biochemical characterization of bacterial methyltransferases reveals necessary residues for sterol side-chain propylation

**DOI:** 10.64898/2025.12.19.695621

**Authors:** Malory O. Brown, Paula V. Welander

**Affiliations:** Department of Earth System Science, Stanford University, Stanford, CA 94305

**Keywords:** C-24 sterol methyltransferase, site-directed mutagenesis, sterol biosynthesis, 24-isopropylcholesterol

## Abstract

The capacity to methylate sterol side-chains via sterol methyltransferases (SMTs) was thought to be widespread in fungi, algae, and plants but limited in animals and nonexistent in bacteria. We have previously demonstrated that yet-uncultured bacteria have the genomic capacity to produce side-chain methylated sterols de novo. Further, we identified three bacterial SMTs capable of producing 24-isopropyl sterols and showed that each of these SMTs was biochemically sufficient for all three of the side-chain methylation steps necessary for 24-isopropyl sterol synthesis. To date, no eukaryotic SMT has been identified that is capable of sequentially adding three methyl groups to generate this propyl structure. To better understand this unique biochemical feature of bacterial SMTs and to potentially identify key amino acid residues involved in sequential methylations on the sterol side-chain, we performed site-directed mutagenesis of propylating bacterial SMTs. Through these analyses, we identified a glycine residue that is necessary but not sufficient for side-chain propylation. This residue is located outside known SMT substrate-binding domains, but inside the active site of an SMT protein model docked with 24-methylenecholesterol. We also show that phenylalanine residues in sterol-binding Region I increase the production of 24-isopropyl sterols, and that 25 residues are conserved among methylating, ethylating, and propylating SMTs. Together, the presence of these residues may allow us to predict if an organism has the genomic capacity to produce C-24 propylated sterols directly from sequencing data, and to generate hypotheses about the environments in which bacterial sterol side-chain propylation may be occurring through analysis of metagenomes. Given the high preservation potential of side-chain alkylated sterols and their use as molecular fossils indicative of ancient life deep in time, a better understanding of the distribution of these unique lipids in modern life and present-day ecosystems will allow for more robust interpretations of side-chain alkylated steranes in the rock record.

## Introduction

Sterol methyltransferases (SMTs) are the only enzymes known to alkylate the side-chain of sterol lipids. These proteins have been extensively characterized due to their potential as antifungal (Pereira et al., 2010; Zhou et al., 2004), anti-leishmanial (Azam et al., 2014), and antiamoebic (Kidane et al., 2017) drug targets. SMTs are also of broad interest due to their importance in understanding the evolution of sterol biosynthesis (Haubrich et al., 2015; Michellod et al., 2023) and the interpretation of sterane molecular fossils in the geologic record (Brown et al., 2023; Gold et al., 2016). For many years, alkylation of the sterol side-chain was thought to be restricted to eukaryotic organisms, specifically plants, fungi, sponges, and algal species like *Chlamydomonas*. More recent work has demonstrated the production of C-24 alkylated sterols in other animals such as annelids (Michellod et al., 2023, Brunoir et al., 2023) and our work has shown that SMT homologs found in bacterial metagenomes are also capable of alkylating sterol side chains. Specifically, SMT homologs from both symbiotic and free-living bacterial metagenomes and metagenome-assembled genomes (MAGs) can produce methylated (C_28_), ethylated (C_29_), and/or propylated (C_30_) sterol in vitro at the C-24 position (Brown et al., 2023). This was the first demonstration that annotated bacterial SMTs are true C-24 sterol methyltransferases and suggests that the production of C-24 alkylated sterols might extend into yet-uncultured bacterial species. Moreover, we observed that a single bacterial SMT is capable of propylating the sterol side-chain in addition to methylating and ethylating – a characteristic not observed with eukaryotic SMTs. However, it is unclear how these novel bacterial SMTs differ biochemically from eukaryotic SMTs.

Eukaryotic SMTs methylate the sterol side-chain at the C-24 or C-28 position at an isolated double bond through an *S*-adenosylmethionine (SAM) dependent mechanism (Nes, 2003). The SMT active site currently remains unverified as no X-ray crystallography, NMR, or Cryo-EM structure of an SMT enzyme has been reported. However, molecular and biochemical studies have identified conserved sterol- and SAM-binding motifs among SMT proteins (Figure 1). Sterol-binding Region I was confirmed through chemical-affinity labeling of the *Saccharomyces cerevisiae* SMT with the irreversible inhibitor 26,27-dehydrozymosterol (Nes et al., 2002) and subsequent site-directed mutagenesis (Ganapathy et al., 2008; Nes et al., 2004). This aromatic-rich region is thought to stabilize the carbocation generated as part of the SMT mechanism (Nes et al., 1998). SAM-binding Region II, which is conserved among SAM-dependent methyltransferases (Kagan & Clarke, 1994), was confirmed through photoaffinity labeling of the *S. cerevisiae* SMT with radiolabeled SAM coupled to site-directed mutagenesis (Jayasimha and Nes, 2008). Region III was identified as a conserved region unique to SMTs (Bouvier-Nave et al., 1998; Jayasimha et al., 2006), and site-directed mutagenesis of the *S. cerevisiae* SMT suggested its role in sterol binding (Ganapathy et al., 2008). Region IV was identified by site-directed mutagenesis of the *S. cerevisiae* SMT, the results of which indicated a role in active site organization (Ganapathy et al., 2008).

**Figure 1.**
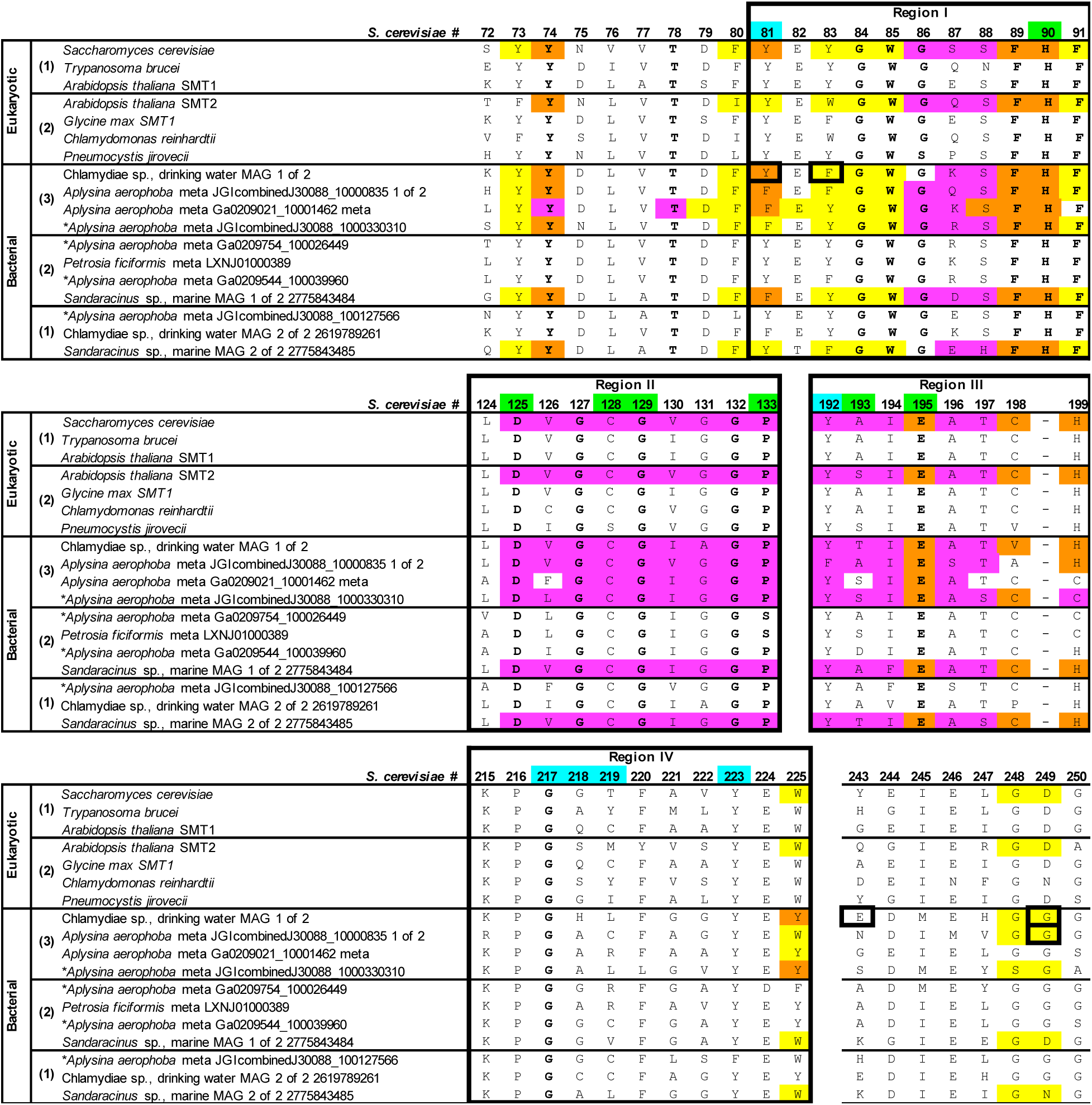
Partial MAFFT-linsi alignment of functional sterol methyltransferase (SMT) protein sequences. Numbers in parenthesis denote the number of methylations performed by the SMT. Bolded residues indicate those conserved among functional SMTs. Boxed residues indicate sites subjected to site-directed mutagenesis in this study. Residues shaded in yellow indicate those within 5 Å of 24-methylenecholesterol docked into the Chl_SMT ColabFold predicted structure or other SMT predicted structures aligned to that SMT. Residues shaded in magenta indicate those within 5 Å of SAM. Residues shaded in orange indicate those within 5 Å of both 24-methylencecholsterol and SAM. Positions shaded in green are essential in *S. cerevisiae* (Ganapathy et al., 2008; Nes et al., 2004). Mutants at the positions shaded in cyan were previously found to permit the *S. cerevisiae* SMT to perform a second methylation (Ganapathy et al., 2008; Nes et al., 1999). The full alignment with additional bacterial sequences is provided in Supplementary Figure 1. *SMT homologs new to this study.

In addition to the identification of regions likely involved in substrate-binding, site-directed mutagenesis studies have identified several amino acid residues essential for C-24 methylation by the *S. cerevisiae* SMT, which methylates zymosterol to produce 24-methyl sterols (Ganapathy et al., 2008; Nes et al., 2004). These studies have also demonstrated specific amino acid changes that permit the *S. cerevisiae* SMT to perform two sequential methylations resulting in 24-ethyl sterols (Figure 1). These include a Y81F mutation in Region I, a Y192F mutation in Region III, and G217L, G218L, T219L and Y223F mutations in Region IV (Ganapathy et al., 2008; Nes et al., 1999). Further, a Y81F mutant of the *Glycine max* SMT1, which normally sequentially methylates cycloartenol to 24(28)-methylene lophenol to produce both 24-methyl and 24-ethyl sterols, resulted in an increase in 24-ethyl sterols. However, this same Y81F mutation of the *Paracoccidiodes brasiliensis* (Pereira et al., 2010) and *Trypanosoma brucei* SMTs (Liu et al., 2011) did not result in the production of 24-ethyl sterols. A phenylalanine at position 81 is therefore insufficient for sterol side-chain ethylation. The other residues where mutations resulted in a gain-of-function of the *S. cerevisiae* SMT are well-conserved among analyzed SMTs (Ganapathy et al., 2008; Nes et al., 1999, 2002) and are not informative when attempting to predict the number of methylations performed by a putative SMT. Site-directed mutagenesis of six Region I aromatic amino acids to leucine each prevented the *G. max* SMT1 from performing a second methylation (Nes et al., 2006) suggesting these residues may help predict if an SMT can perform two sequential methylations. However, these aromatic residues are also highly-conserved among all SMTs.

While informative from a biochemical perspective, the residue changes that affect the ability of an SMT to sequentially methylate the sterol side-chain to produce 24-ethyl sterols are neither necessary nor sufficient for 24-ethyl sterol production. Therefore, the occurrence of these residues alone does not allow us to discern whether an annotated SMT is capable of methylation, ethylation, and/or propylation. The number of SMT gene copies has previously been used to predict the ability of an organism to produce 24-propyl sterols (Gold et al., 2016; Love et al., 2020). However, the three propylating bacterial SMTs we recently identified are each capable of catalyzing the three side-chain methylation steps necessary for 24-isopropyl sterol biosynthesis via a single SMT copy. Thus, identification of key amino acid residues necessary and/or sufficient for side-chain propylation would greatly enhance our ability to determine if an SMT can produce 24-isopropyl sterols directly from sequencing data. This is particularly significant given that geologic 24-isopropylcholestane, historically a biomarker for demosponges, may represent the first evidence for animals on Earth (Love et al., 2009). However, debates over this interpretation continue given the potential for alternative sources such as pelagophyte algae, Rhizaria, bacteria, or abiotic processes (Antcliffe, 2013, 2015; Bobrovskiy et al., 2021; Brown et al., 2023; Hallmann et al., 2020; Love et al., 2020; Love & Summons, 2015; Nettersheim et al., 2019). The ability to predict if an SMT can produce 24-isopropyl sterols directly from sequencing data will allow us to identify SMTs that warrant testing in the laboratory to resolve the interpretation of the 24-isopropylcholestane biomarker. This predictive power would prove especially useful for analyzing the thousands of untested SMT homologs from yet-uncultured bacteria and microbial eukaryotes currently found in metagenomic databases.

Here, we performed site-directed mutagenesis of bacterial SMTs that propylate sterols at the C-24 position to identify potential biochemical requirements for sterol side-chain propylation versus methylation and/or ethlyation. We identified a glycine residue that is necessary but not sufficient for side-chain propylation. This residue is located outside known SMT substrate-binding domains, but inside the active site of an SMT protein model docked with 24-methylenecholesterol. We also show that phenylalanine residues in sterol-binding Region I increase the production of 24-isopropyl sterols, and that 25 residues are conserved among functional SMTs. Together, the presence of these residues may allow us to predict if an organism has the genomic capacity to produce side-chain propylated sterols directly from sequencing data, and to generate hypotheses about the environments in which bacterial sterol side-chain propylation may be occurring through analysis of metagenomes.

## Results

### Mutations of a propylating Chlamydiae sp. SMT alter sterol composition

To identify potential residues required for sterol side-chain propylation, we first performed site-directed mutagenesis of a propylating SMT. We chose the “Chlamydiae sp., drinking water MAG 1 of 2” SMT (Chl_SMT) as it was the only SMT we previously identified that produced both 24*S*-24-isopropylcholest-5,25-dienol and 24-isopropylcholest-5,24-dienol at levels detectable by GC-MS (Brown et al., 2023). All residue positions are referred to using *S. cerevisiae* numbering for consistency and ease of comparison to previously published site-directed mutagenesis studies. Corresponding numbering for the Chl_SMT and “*Aplysina aerophoba* meta JGIcombinedJ30088_10000835 1 of 2” (Ap_sym_SMT), can be found in Supplementary Table 2.

We mutated Chl_SMT at 13 residues (Supplementary Table 1), 11 of which are conserved among at least two of the three propylating SMTs we previously tested, but not among other functional bacterial SMTs (S68, A152, Q175, H180, I239, G249, T312, N318, L343, E350, F362). Rather than mutating each residue to alanine, we chose to mutate these residues to the corresponding amino acid encoded by the *S. cerevisiae* SMT, as this SMT is known to add only one carbon to the sterol side-chain. The one exception was H180, which was mutated to asparagine, the most common amino acid at position 180 among ethylating bacterial SMTs. We also chose to mutate V204 to glutamic acid, as glutamic acid at this position was previously hypothesized to be a methyltransferase site in promiscuous SMTs (Gold et al., 2016). We also mutated E243, as several acidic residues have been previously hypothesized to be involved in sterol or SAM binding (Nes et al., 2004), to alanine, the most common amino acid at this position among ethylating bacterial SMTs.

Each of these mutants was tested in vitro with both desmosterol and 24-methylenecholesterol, and the sterol products were analyzed with GC-MS. The relative abundances of C_28_ (24-methyl), C_29_ (24-ethyl), and C_30_ (24-isopropyl) sterol products for each mutant based on peak areas are shown in Table 1. The relative abundances of C_28_, C_29_, and C_30_ sterols produced by the wild type Chl_SMT, averaged from five reactions with desmosterol, were 22%, 51%, and 27%, respectively. The relative abundances of C_29_ and C_30_ sterols produced by the wild type SMT, averaged from five reactions with 24-methylenecholesterol, were 50% for both structures. Several of the initial 13 mutants showed altered sterol composition, but only two of these mutations resulted in a >20% decrease in the relative abundance of C_30_ sterols in reactions with both desmosterol and 24-methylenecholesterol as the substrate. In the reaction with desmosterol, 2% of the sterols produced by the E243A mutant were C_30_ sterols, down from 27%. In the reaction with 24-methylenechoelsterol, 7% of the sterols produced by this mutant were C_30_ sterols, down from 50%. The G249D mutant still produced the three C_28_ sterols observed as a product of the wild type SMT with desmosterol as the substrate, but C_29_ and C_30_ sterols were no longer detectable (Figure 2a). We chose to focus our subsequent analyses on this glycine residue given that the G249D mutant retained methylation activity but no longer produced detectable 24-ethyl or 24-isopropyl sterols in vitro.

**Figure 2.**
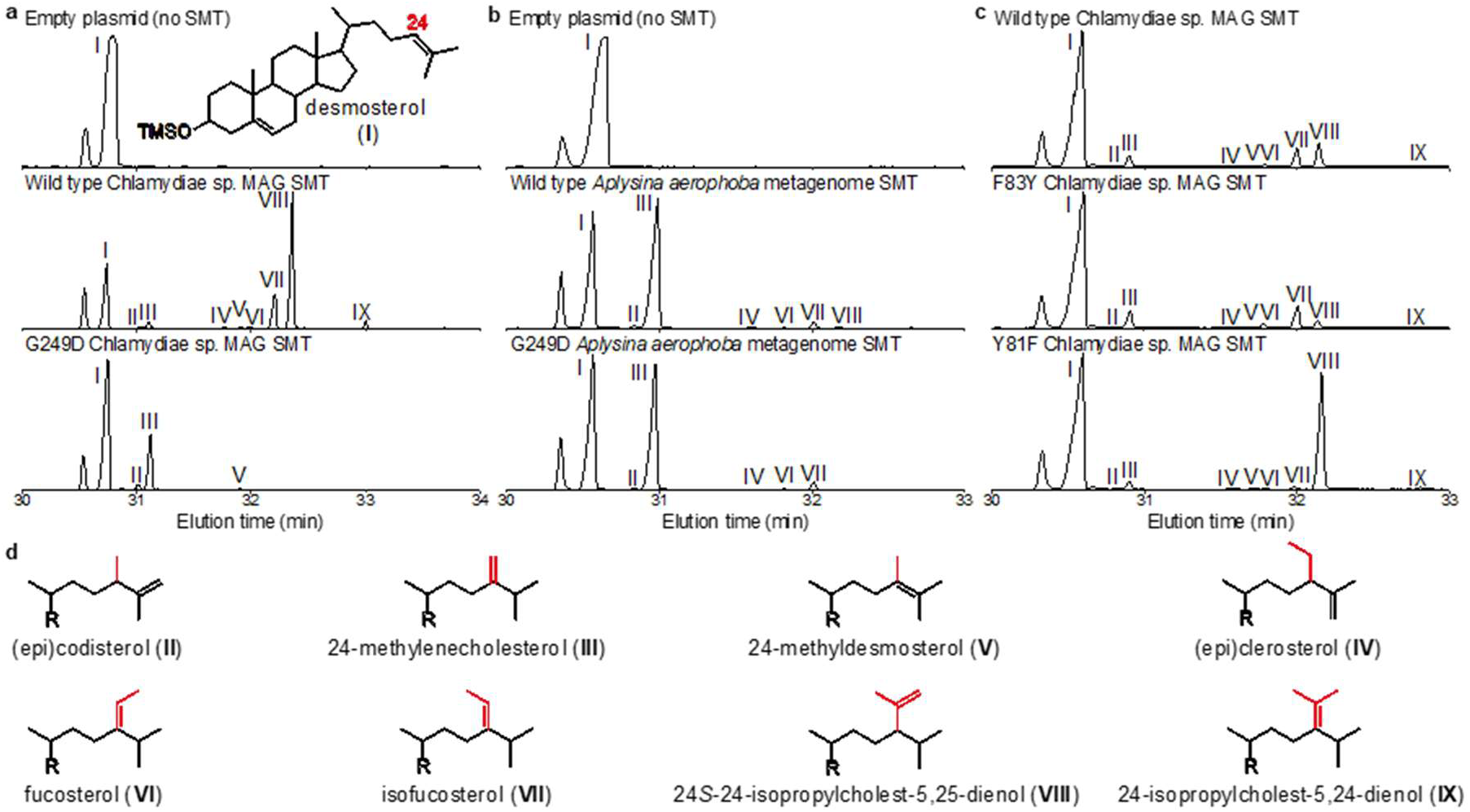
Mutations at Gly249, Phe83, and Tyr81 alter sterol side-chain propylation in vitro. Representative extracted ion chromatograms (*m/z* 456, 470, 484, 498) of total lipid extracts from in vitro reactions performed with desmosterol and *E. coli* lysates. **a.** Reactions performed with an empty plasmid, the wild type Chl_SMT and its G249D (G211D) mutant. **b**. Reactions performed with an empty plasmid, the wild type Ap_sym_SMT, and its G249D (G225D) mutant. **c**. Reactions performed with the wild type Chl_SMT and its F83Y (F45Y) and Y81F (Y43F) mutants. **d**. Side-chain structures of sterols identified in a-c. All lipids were derivatized to trimethylsilyls prior to GC-MS analysis.

**Table 1.** Percentages of C_28_, C_29_, and C_30_ sterols relative to total sterol products of “Chlamydiae sp., drinking water MAG 1 of 2” wild type (WT) and mutant SMT in vitro reactions performed with desmosterol or 24-methylenecholesterol as the substrate.

| Mutation | Desmosterol |  |  |  | 24-Methylenecholesterol |  |  |
| --- | --- | --- | --- | --- | --- | --- | --- |
|  | C<sub>28</sub> | C<sub>29</sub> | C<sub>30</sub> | change in C<sub>30</sub> | C<sub>29</sub> | C<sub>30</sub> | change in C<sub>30</sub> |
| WT | 22% | 51% | 27% | NA | 50% | 50% | NA |
| S68E | 14% | 46% | 40% | 13% | 32% | 68% | 18% |
| Y81F | 18% | 9% | 73% | 46% | 9% | 91% | 41% |
| F83Y | 30% | 66% | 4% | -23% | 75% | 25% | -25% |
| A152D | 11% | 71% | 17% | -10% | 43% | 57% | 7% |
| Q175K | 25% | 46% | 29% | 2% | 49% | 51% | 1% |
| H180N | 18% | 61% | 21% | -6% | 61% | 39% | -11% |
| V204E | 18% | 60% | 22% | -5% | 47% | 53% | 3% |
| I239R | 12% | 63% | 26% | -1% | 57% | 43% | -7% |
| E243A | 45% | 52% | 2% | -25% | 93% | 7% | -43% |
| **G249D** | **100%** | **0%** | **0%** | **-27%** | **0%** | **0%** | **-50%** |
| T312L | 11% | 53% | 36% | 9% | 26% | 74% | 24% |
| N318T | 27% | 63% | 9% | -18% | 70% | 30% | -20% |
| L343N | 22% | 58% | 20% | -7% | 34% | 66% | 16% |
| E350A | 9% | 67% | 25% | -2% | 65% | 35% | -15% |
| F362L | 9% | 68% | 24% | -3% | 65% | 35% | -15% |

### Gly249 is necessary but not sufficient for sterol side-chain propylation

We were surprised that the G249D mutant of the Chl_SMT still methylated the sterol side-chain but lost the ability to produce detectable C_29_ and C_30_ sterols given that this residue lies outside Regions I-V and both conserved protein family (pfam) domains in our MAFFT-linsi alignment (Figure 1, Supplementary Figure 1). To confirm that this mutation eliminates the production of C_30_ sterols in vitro, we generated the same mutation in the *Aplysina aerophoba* SMT, Ap_sym_SMT. Similarly to the G249D mutant of the Chl_SMT, the G249D mutant of the Ap_sym_SMT no longer produced detectable C_30_ sterols in vitro (Table 2). However, the G249D mutant of the Ap_sym_SMT retained its ability to produce all three of the C_29_ sterols produced by the wild type SMT (Figure 2b). This mutant produced 59% of the C_29_ sterols from desmosterol and 85% of the C_29_ sterols from 24-methylenecholesterol relative to the wild type SMT (Table 2.).

**Table 2.** Percentages of C_28_, C_29_, and C_30_ sterol products of total sterol products of “Chlamydiae sp., drinking water MAG 1 of 2” and “*Aplysina aerophoba* meta JGIcombinedJ30088_10000835 1 of 2” SMT mutants relative to wild type (WT).

| SMT | Mutation | Desmosterol |  |  | 24-Methylenecholesterol |  |
| --- | --- | --- | --- | --- | --- | --- |
|  |  | C<sub>28</sub> | C<sub>29</sub> | C<sub>30</sub> | C<sub>29</sub> | C<sub>30</sub> |
| *Aplysina aerophoba* meta | WT | 100% | 100% | 100% | 100% | 100% |
|  | G249D | 48% | 59% | 0% | 85% | 0% |
| Chlamydiae sp. MAG | WT | 100% | 100% | 100% | 100% | 100% |
|  | G249A | 908% | 27% | 0% | 48% | 0% |
|  | G249S | 943% | 10% | 0% | 36% | 0% |
|  | G249N | 815% | 14% | 0% | 38% | 0% |
|  | G249T | 461% | 1% | 0% | 15% | 0% |
|  | G249V | 788% | 0% | 0% | 0% | 0% |
|  | G249C | 556% | 0% | 0% | 0% | 0% |
|  | G249D | 441% | 0% | 0% | 0% | 0% |
|  | G249I | 405% | 0% | 0% | 0% | 0% |
|  | G249M | 362% | 0% | 0% | 0% | 0% |
|  | G249L | 319% | 0% | 0% | 0% | 0% |
|  | G249F | 283% | 0% | 0% | 0% | 0% |

Given that G249D mutations of both propylating SMTs we tested resulted in the loss of detectable C_30_ sterols when given desmosterol and 24-methylenecholesterol as substrates, and that this glycine residue is conserved among all three of the propylating SMTs we previously identified (Figure 1), we decided to test additional bacterial SMTs that encode glycine at position 249. We chose nine bacterial SMTs encoding this glycine, all from sponge metagenomes, for in vitro analysis. We also tested one additional bacterial SMT, also from a sponge metagenome, with a two amino acid gap at this position in our alignment (Supplementary Figure 1). The sterols detected as products of these additional SMTs in reactions with desmosterol or 24-methylenecholesterol are shown in Table 3. Only four of the 10 were functional in our in vitro experiments, and the “*Aplysina aerophoba* meta JGIcombinedJ30088_1000330310” SMT was the only one to produce C_30_ sterols.

**Table 3.** Sterols identified as products of bacterial sterol methyltransferases with glycine or gaps at the 249.

|  | C<sub>28</sub> |  |  | C<sub>29</sub> | C<sub>30</sub> |  |
| --- | --- | --- | --- | --- | --- | --- |
|  | (II) | (III) | (V) | (VI) | (VIII) | (IX) |
| *Aplysina aerophoba* meta JGIcombinedJ30088_1000330310 | + | + | + | + | + | + |
| *Aplysina aerophoba* meta Ga0209754_100026449 |  | + |  | + |  |  |
| *Aplysina aerophoba* meta Ga0209544_100039960 | + | + | + | + |  |  |
| *Aplysina aerophoba* meta JGIcombinedJ30088_100127566 | + | + |  |  |  |  |
| *Aplysina aerophoba* meta JGIcombinedJ30088_100153964 |  |  |  |  |  |  |
| *Aplysina aerophoba* meta JGIcombinedJ30088_100077196 |  |  |  |  |  |  |
| *Aplysina aerophoba* meta JGIcombinedJ30088_100209763 |  |  |  |  |  |  |
| *Aplysina aerophoba* meta JGIcombinedJ30088_100057164 |  |  |  |  |  |  |
| *Sarcotragus foetidus* meta LXNI01006544 |  |  |  |  |  |  |
| *Sarcotragus foetidus* meta LXNI01016685 |  |  |  |  |  |  |

The results of our in vitro experiments with additional metagenomic SMTs demonstrated that glycine at the 249 position alone is not sufficient for C_30_ sterol production from desmosterol and 24-methylenecholesterol. We next asked if this residue is essential for an SMT to produce 24-isopropyl sterols. To address this question, we generated ten additional 249 mutants of the Chl_SMT for in vitro testing. G249 was mutated to alanine, asparagine, cysteine, isoleucine, leucine, methionine, phenylalanine, serine, threonine, and valine, and each mutant was again tested with both desmosterol and 24-methylenecholesterol. Each additional 249 mutant retained its ability to produce C_28_ sterols from desmosterol (Table 2). Four 249 mutants produced detectable C_29_ sterols from desmosterol and 24-methylencecholesterol (G249A, G249S, G249N, and G249T). None of these additional 249 mutants produced detectable C_30_ sterols.

### Structure predictions and docking experiments suggest glycine 249 resides in the SMT active site

The results of our in vitro experiments suggest glycine 249 is necessary for an SMT to produce 24-isopropyl sterols. We therefore hypothesized that this residue exists in the active site of the SMT protein. To test this hypothesis, we first generated a protein structure prediction of the Chl_SMT using AlphaFold2 (Mirdita et al., 2022). The highest-ranked model is shown in Figure 3. While G249 occurs outside Regions I-IV and both pfam domains in our protein sequence alignment (Figure 1), it exists near sterol-binding Region I in the structure prediction (Figure 3a, b). Specifically, G249 occurs within 10 Å of sterol-binding Region I Y81, a residue that was previously shown to affect the ability of a eukaryotic SMT to sequentially methylate the sterol side-chain to produce 24-ethyl sterols (Nes et al., 1999, 2006). E243, which we found decreases the relative abundance of C_30_ sterols when mutated to alanine, does not exist in this region in our model.

**Figure 3.**
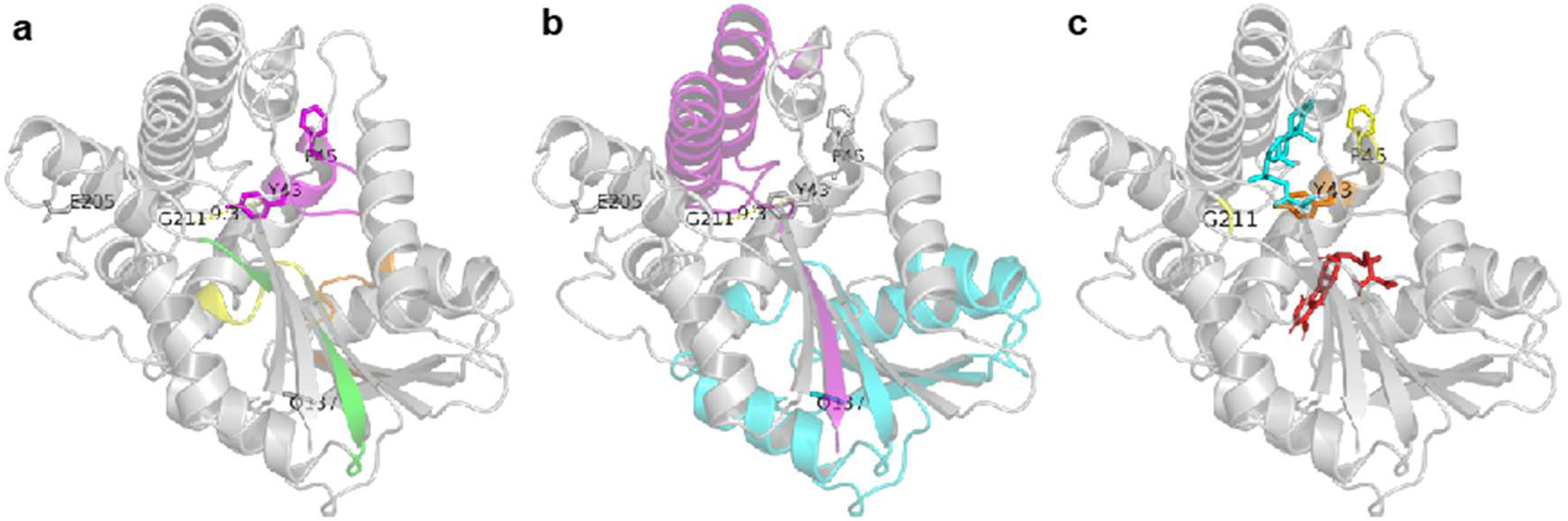
Glycine 249 (G211) occurs outside known substrate-binding regions and pfam domains but near the sterol-binding site in a structure of the Chl_SMT predicted by ColabFold-AlphaFold2. **a**. Structure prediction is colored by substrate binding regions. Region I is magenta, Region II is orange, Region III is yellow, and Region IV is green. The distance between the centroids of G249 (G311) and Y81 (Y43) is 9.3 Å. **b.** Structure prediction is colored by pfam domains. The methyltransferase domain is cyan, and the sterol MT c-term domain is purple. **c**. Chl_SMT model with docked 24-methylenecholesterol (cyan) and SAM (red).

To provide additional evidence that G249 exists in the SMT active site, we performed docking experiments with both 24-methylenecholesterol and SAM using the Vina function in the PyMol plugin DockingPie (Rosignoli & Paiardini, 2022). In the best-scoring docking scenario, the sterol side-chain is positioned between G249 and Y81 (Figure 3c). To determine if this active site structure is conserved among other SMTs, we generated protein structure predictions for three other bacterial propylating SMTs from *A. aerophoba* metagenomes. We also generated models for the plant *A. thaliana* SMT2 and the bacterial *Sandaracinus* sp. MAG SMT 1 of 2, which ethylate sterols, and the yeast *S. cerevisiae* SMT and the *Sandaracinus* sp. MAG SMT 2 of 2, which methylate sterols (Brown et al., 2023). After aligning these models to the Chl_SMT predicted structure docked with 24-methylenecholesterol and SAM, we found that residue 249 exists within 5 Å of 24-methylenecholesterol in all but one SMT model (Figures 4). Residue 81 exists within 5 Å of 24-methylenecholesterol in all eight SMT models. Structure predictions of propylating SMTs aligned to methylating and ethylating SMTs and colored by root-mean-square-deviation (RMSD) of atomic coordinates suggest no major structural deviations in this region of the SMT models (Figure 4). RMSD values also suggest that propylating bacterial SMTs are generally more structurally similar to other bacterial SMTs than to eukaryotic SMTs. However, one set of RMSD calculations suggests that the Chl_SMT model is most similar to the *S. cerevisiae* SMT (Figure 4f).

**Figure 4.**
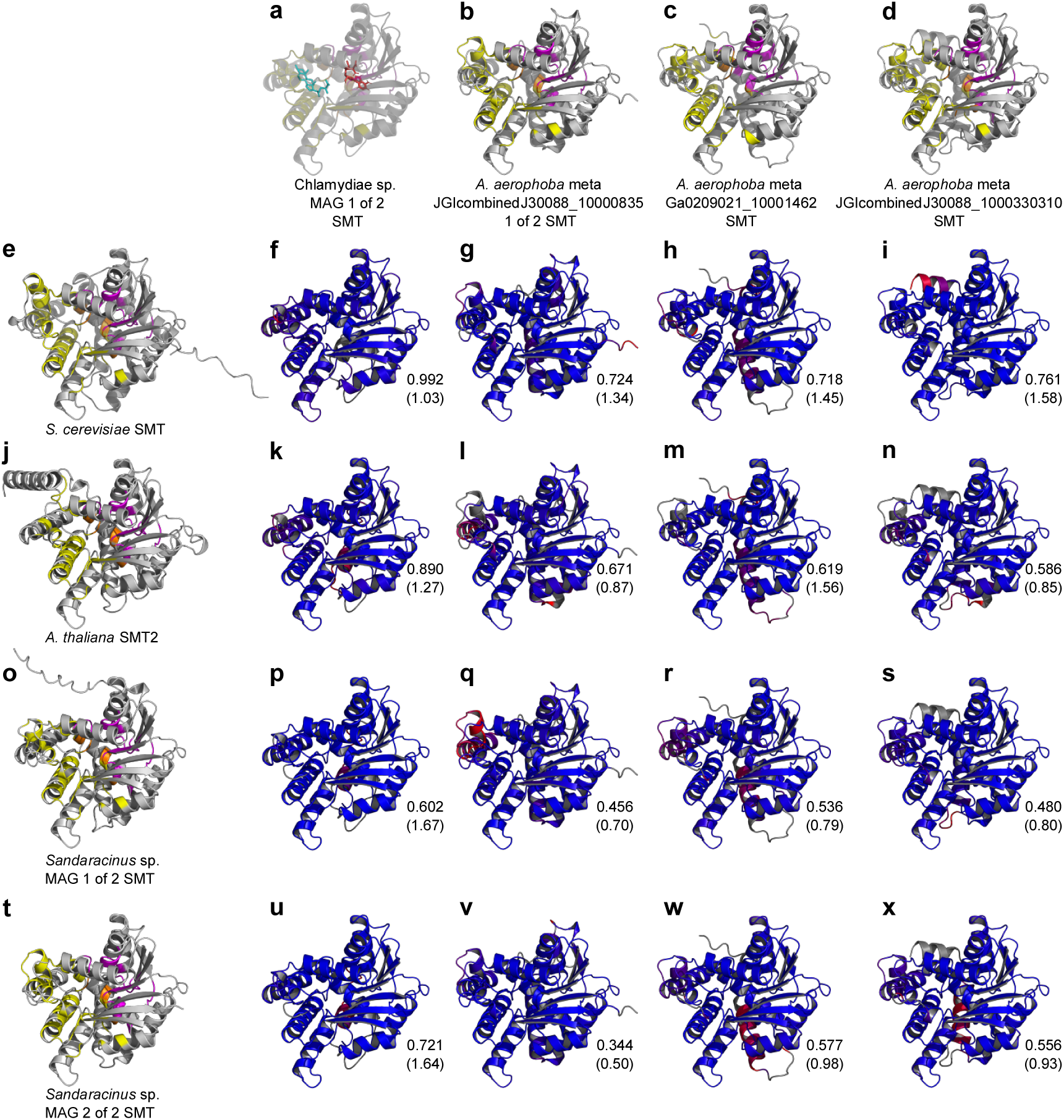
Highest-ranked SMT protein structure predictions. **a.** Chl_SMT model with docked 24-methylenecholesterol (cyan) and SAM (red). **b-d.** Other propylating bacterial SMTs. **e.** *Saccharomyces cerevisiae* SMT, also known as ERG6. **j.** *Arabidopsis thaliana* SMT2. **o.** *Sandaracinus* sp. MAG SMT 1 of 2. **t.** *Sandaracinus* sp. MAG SMT 2 of 2. Yellow residues indicate those within 5 Å of 24-methylenecholesterol, magenta residues indicate those within 5 Å of SAM, and orange indicate those within 5 Å of both 24-methylencecholsterol and SAM. **f-i.** Propylating SMTs aligned against the *S. cerevisiae* SMT. **k-n.** Propylating SMTs aligned against *A. thaliana* SMT2. **p-s.** Propylating SMTs aligned against *Sandaracinus* sp. MAG SMT 1 of 2. **u-x.** Propylating SMTs aligned against *Sandaracinus* sp. MAG SMT 2 of 2. Residues are colored by RMSD with blue indicating the lowest values (most similar) and red indicating the highest values (least similar). Numbers in the lower right corners indicate overall RMSD values assigned by the align function and, in parenthesis, ColorByRMSD, both in PyMOL. Structures were predicted using ColabFold-AlphaFold2.

### Region I phenylalanines increase C_30_ sterol production in vitro

Glycine 249 is conserved among all four of the propylating SMTs we identified (Figure 1) and appears to be necessary but not sufficient for bacterial C_30_ production. Thus, the presence of a glycine at this position in a protein sequence alone does not imply an SMT will produce 24-isopropyl sterols. We therefore sought to identify additional residues involved in C_30_ sterol production to refine predictions. In our MAFFT-linsi alignment, we found that all four of the propylating SMTs we tested encode a phenylalanine in Region I at the 81 and/or 83 position (Figure 1, Supplementary Figure 1). Given that a tyrosine to phenylalanine (Y81F) mutation of the *S. cerevisiae* SMT allowed the production of C_30_ sterols (Nes et al., 1999) but not in *T. brucei* (Liu et al., 2011) or *P. brasiliensis* (Pereira et al., 2010), we first generated this same Y81F mutation in the Chl_SMT to confirm that phenylalanine at this position increases sequential methylation in a bacterial SMT. 73% of the sterols produced by the Y81F mutant from desmosterol were C_30_ sterols compared to 27% for the wild type Chl_SMT(Figure 2c). Further, 91% of the sterols produced by this mutant when provided 24-methylenecholesterol as a substrate were C_30_ sterols, an increase from the 50% observed for the wild type Chl_SMT (Table 1). Given that this extra Region I phenylalanine does appear to increase sequential methylation in a bacterial SMT, we next sought to determine if a decrease in Region I phenylalanines decreases sequential methylation. We generated a F83Y mutant of the Chl_SMT and tested its activity in vitro. This mutant still produced C_30_ sterols from both desmosterol and 24-methylenecholesterol (Table 1). In the reaction with desmosterol, 4% of the sterols produced by the F83Y mutant were C_30_ sterols, a decrease from the 27% observed for the wild type protein (Figure 3c). In the reaction with 24-methylenecholesterol, 25% of the sterols produced by this mutant were C_30_ sterols compared to the 50% observed for the wild type.

### Identification of 25 residues conserved among functional SMTs

Our data suggests that Region I phenylalanine residues at position 81 and 83 in an SMT appear to increase C_30_ sterol production. However, they alone are not sufficient for producing 24-isopropyl sterols as nine of the functional bacterial SMTs we tested encode phenylalanine at one of these positions but do not produce C_30_ sterols in our in vitro experiments (Figure 1, Supplementary Figure 1). We hypothesized that phenylalanine at 81 and/or 83 and glycine at 249 together may be sufficient for an SMT to produce 24-isopropyl sterols. However, three of the bacterial SMTs we tested encode these residues but only perform one or two side-chain methylations (Figure 1). Further, one SMT we tested, *“Sarcotragus foetidus* meta LXNI01016685”, meets both requirements but did not methylate desmosterol or 24-methylenecholesterol in vitro (Table 3, Supplementary Figure 1). We therefore expanded our analyses to residues conserved among functional SMTs that might aid in predicting if an SMT is capable of any type of alkylation.

We identified 25 residues that are conserved among all functional SMTs in our MAFFT-linsi alignment (Figure 1, Supplementary Figure 1). These include the 84GWGXXFHF91 motif of Region I, the 125DXGXGXXGP133 motif of Region II, 195E in Region III, and 217G of Region IV. Additional residues conserved among functional SMTs include 147G, 154Q, and 189D of the methyltransferase pfam domain, 333G, 341L, 352G, 357F, and 359P of the sterol MT C-term pfam domain, and 74Y, 78T, 268G, and 286W outside all known sterol-binding, SAM-binding, and pfam domains. Deviations from at least one of these conserved residues are found in seven out of the 11 SMTs we tested that did not methylate desmosterol or 24-methylenecholesterol in vitro. The four SMTs that retain all 12 of the residues conserved among functional SMTs but did not function in our experiments include “Hot spring, Beatty, Nevada meta_Ga0114945_100210811”, “*Aplysina aerophoba meta* JGIcombinedJ30088_100209763”, “*Sarcotragus foetidus* meta LXNI01006544”, and “*Sarcotragus foetidus* meta LXNI01016685”. The “*Sarcotragus foetidus* meta LXNI01006544” SMT has a two amino acid gap at position 249, while the “*Sarcotragus foetidus* meta LXNI01016685” SMT encodes a proline insertion in Region III. “Hot spring, Beatty, Nevada meta_Ga0114945_100210811” and “*Aplysina aerophoba meta* JGIcombinedJ30088_100209763” SMTs have no obvious deviations. All SMTs we tested retain G84, W85, H90, and F91 in Region I, D125, G127, G129, and P133 in Region II, E195 in Region III, G217 in Region IV, G333, G252, P359 in the sterol MT-C pfam domain, and W286 regardless of their function in vitro.

## Discussion

In this study, we identified a novel residue involved in determining the number of alkalytions added to a sterol side-chain at the C-24 position by a sterol methyltransferase. Our protein modeling and docking experiments suggest residue 249 is positioned in the SMT active site near the sterol side-chain and is found outside the known sterol- and SAM-binding domains identified in all C-24 sterol methyltransferases. Each of the four SMTs demonstrated to produce 24-isopropyl sterols encode glycine at this position, and our in vitro analyses of wild type and mutant SMTs suggest this glycine is necessary but not sufficient for 24-isopropyl sterol biosynthesis. None of the 11 glycine 249 mutants of the Chl_SMT we tested produced C_30_ sterols in vitro. However, four retained the ability to produce C_29_ sterols (G249A, G249S, G249N, G249T). Glycine, alanine, and serine are small amino acids, and asparagine and threonine are slightly larger but still considered small. The larger amino acids we tested at this position did not support sequential methylations. Thus, the size of the 249 residue appears to be the major factor affecting the number of methylations performed by an SMT, likely due to steric hindrance in the binding site. Further, glycine residues are proposed to provide flexibility in enzyme active sites (Yan & Sun, 1997). Thus, as the sterol substrate is sequentially alkylated, the glycine at position 249 may accommodate these larger side-chains by increasing flexibility in the SMT active site. While additional SMTs capable of producing 24-isopropyl sterols with other amino acid residues at the 249 position may yet be discovered, our data suggest that a glycine at 249 indicates that an SMT may be capable of producing 24-isopropyl sterols and warrants testing in the laboratory.

Our results also suggest that, in combination with a glycine at 249, phenylalanine at 81 and/or 83 can provide further evidence that an SMT may produce 24-isopropyl sterols. An F81Y mutation of the Chl_SMT resulted in decreased C_30_ sterol production in our experiments. While we did not perform extensive mutagenesis at these sites, Y81 of the *S. cerevisiae* SMT was previously mutated to phenylalanine, tryptophan, alanine, isoleucine, leucine, and valine (Nes et al., 2008). Only the phenylalanine mutant resulted in 24-ethyl sterols. This gain-of-function of the *S. cerevisiae* SMT Y81F mutant was attributed to decreased steric hindrance in the sterol binding site (Nes et al., 2006), consistent with our structural analyses. Further, this same Y81F mutation of the soybean *G. max* SMT1 resulted in an increase in 24-ethyl sterols (Nes et al., 2006). However, one SMT we tested, “Chlamydiae sp., drinking water MAG 2 of 2”, encodes a phenylalanine position 81 but produces only 24-methyl sterols in vitro. Thus, F81 is neither necessary nor sufficient for sequential side-chain methylations. The Y83F mutant of the Chl_SMT resulted in increased C_30_ sterol production in our experiments, but this same mutation of the *S. cerevisiae* SMT did not results in 24-ethyl sterols (Ganapathy et al., 2008). Further, four SMTs we tested encode a phenylalanine at position 83 but produce only 24-methyl sterols in vitro. Thus, F83 is also neither necessary nor sufficient for sequential side-chain methylations. However, our results currently suggest that SMTs encoding glycine at the 249 position and phenylalanine at the 81 and/or 83 position are good candidates for 24-isopropyl sterol production and warrant further testing.

In addition to glycine at 249 and phenylalanine at 81 and/or 83, deviations from the 25 residues conserved among functional SMTs identified here can also be helpful in predicting the function of an SMT directly from sequencing data. Five of the conserved residues, H90, D125, G129, P133, and E195, were previously shown to be essential in the *S. cerevisiae* SMT (Ganapathy et al., 2008; Nes et al., 2004). These studies suggested that H90 is involved in deprotonation of the sterol substrate, D12, G129, and P133 are involved in binding of the SAM cofactor, and E195 is involved in sterol binding. The essentiality and function of the remaining 20 conserved residues can be confirmed by additional site-directed mutagenesis. Further, additional in vitro testing of SMTs that did not methylate desmosterol or 24-methylencecholesterol may reveal these SMTs function but require different sterol substrates. Until then, alternative amino acids at these 25 conserved positions may indicate that an SMT is not functional and is therefore a poor candidate for 24-isopropyl sterol biosynthesis and laboratory testing.

While an SMT crystal structure would greatly enhance future mutagenesis studies and allow for more robust predictions of SMT function directly from sequencing data, our mutagenesis and modeling results suggest that SMTs encoding the 25 conserved residues, a glycine at 249, and a phenylalanine at 81 and/or 83 currently represent the most promising candidates for 24-isopropyl sterol biosynthesis. Identification of these residues in an SMT in a genome from an organism in which 24-isopropyl sterols have been found, such as a demosponge, may provide preliminary evidence that the organism can synthesize these sterols de novo. Further, the presence of these residues in SMTs from organisms, such as free-living bacteria, or environments, such as freshwater, currently unknown to produce 24-isopropyl sterols may suggest alternative sources of the 24-isopropylcholestane biomarker. The function of these SMTs can then be confirmed in the laboratory, allowing for an improved understanding of side-chain methylated sterol biosynthesis in modern organisms and more robust interpretations of side-chain methylated steranes in the rock record.

## Methods

### SMT identification and artificial synthesis

Twenty-four bacterial SMTs were previously identified in Brown et al., 2023. Additional SMT homologs were identified in the Joint Genome Institute Integrated Microbial Genomes & Microbiomes (JGI IMG; https://img.jgi.doe.gov) or GenBank (https://www.ncbi.nlm.nih.gov/genbank/) databases using the *Saccharomyces cerevisiae* S299C SMT amino acid sequence as the BLASTP search query (locus tag: YML008C, maximum e-value: 1e^−50^, minimum percent identity: 30%). Ten metagenomic SMT homologs that contained the conserved methyltransferase and C-terminal domains as identified using HmmerWeb and glycine or gaps at the 249 position (yeast numbering) in our alignment were chosen for additional in vitro analyses. These SMT DNA sequences were codon-optimized for expression in *E. coli* and artificially synthesized through the Department of Energy Joint Genome Institute (DOE JGI) DNA Synthesis Science Program and obtained in the IPTG-inducible plasmid pSRKGm-*lac*UV5-rbs5 (Banta et al., 2017) in *E. coli* TOP10. Codon-optimized DNA sequences for the two SMTs that underwent site-directed mutagenesis are provided in Supplementary Table 1.

### Site-directed mutagenesis

Plasmid site-directed mutagenesis was performed using a modified single-mutagenic primer method(Y. Huang & Zhang, 2017). Briefly, each 25 uL reaction consisted of 100 ng of plasmid DNA, 0.2 µM of a primer encoding the desired change, 0.2 µM dNTPs, and 2.5 U PfuUltra II Fusion HS DNA Polymerase (Agilent). Cycling conditions were as follows: 95 °C for 3 min, 30 cycles of 95 °C for 30 sec, 55 °C for 1 min, and 65 °C for 7 min, and 65 °C for 5 min. Reactions were then digested with 1 µL *DpnI* for 12 hours at 37 °C. 2 µL of each heat-killed reaction was then transformed into electrocompetent *E. coli* DH10B using a MicroPulser Electroporator (BioRad). Oligonucleotides were purchased from Integrated DNA Technologies (Coralville, IA). Plasmid DNA was isolated using the GeneJET Plasmid Miniprep Kit (ThermoFisher Scientific). Plasmid DNA was sequenced to confirm the desired changes by ELIM Biopharm (Hayward, CA) using the following primers: 5′-AATGCAGCTGGCACGACAGG-3′ (forward) and 5′-CCAGGGTTTTCCCAGTCAC-3′ (reverse).Mutagenic primer sequences are provided in Supplementary Table 2.

### Bacterial culture and heterologous expression

*E. coli* expression strains were cultured in 50 mL terrific broth (TB) supplemented with gentamycin (15 μg/mL) at 37 °C while shaking at 225 rpm. Cultures were induced with 500 μM isopropyl β-D-1-thiogalactopyranoside (IPTG) at an OD_600_ of ∼0.6, then incubated an additional 4 hours at 30 °C while shaking at 225 rpm. Cells were harvested by centrifugation at 4,500 × g for 10 minutes at 4 °C. Cell pellets were stored at -80 °C until sonication.

### Sterol methyltransferase assay

Cells pellets were resuspended in 5 mL buffer containing the following: 50 mM Tris-HCl, 2 mM MgCl_2_, 20% glycerol (v/v), and 0.1% β-mercaptoethanol (v/v), pH 7.5. Cells were then lysed on ice via a Qsonica Sonicator Q500 equipped with a 3.2 mm probe at 30% amplitude pulsing at 5 seconds on, 15 seconds off for 8 minutes of total on time. Lysates were then partially clarified by centrifugation at 4,500 × g for 10 minutes at 4 °C. Total protein concentration in the resulting supernatant was quantified using a Coomassie (Bradford) Protein Assay Kit (Thermo Scientific) according to the manufacturer’s Standard Microplate Protocol and a BioTek Synergy HT Microplate Reader. SMT assays were then immediately prepared using enough supernatant to give 6,500 μg total protein. Other reaction components included 100 μM desmosterol or 24-methylenecholesterol (Avanti Polar Lipids, Inc.), 100 μM S-(5′-sdenosyl)-L-methionine chloride dihydrochloride (Sigma-Aldrich), and 0.1% Tween-80 (v/v) to a total reaction volume of 400 μL. Reactions were held at 30 °C for 20 hours and then stored at -20 °C until lipid extraction.

### Lipid extractions

Lipids were extracted using a modified Bligh-Dyer method (Bligh & Dyer, 1959; Welander et al., 2012) with the completed in vitro reactions as the water phase. Reactions were sonicated in 10:5:4 (vol:vol:vol) methanol:dichloromethane:water for 1 hour. The organic phase was then separated with twice the volume of 1:1 (vol:vol) dichloromethane:water followed by storage at -20 °C for >1 hr. Following centrifugation at 2,800 x g for 10 min at 4 °C, the organic phase was transferred and evaporated under N_2_ to give total lipid extracts (TLEs). TLEs were derivatized to trimethylsilyl ethers in 1:1 (vol:vol) pyridine:Bis(trimethylsilyl)trifluoroacetamide for 1 hr at 70 °C prior to GC-MS analysis.

### Gas Chromatography-Mass Spectrometry (GC-MS) analysis

Lipids were separated with an Agilent 7890B Series GC equipped with two Agilent DB-17HT columns (30 m x 0.25 mm i.d. x 0.15 μm film thickness) in tandem with helium as the carrier gas at a constant flow of 1.1 ml/min and programmed as follows: 100°C for 2 min, then 12°C/min to 250°C and held for 10 min, then 10°C/min to 330°C and held for 17.5 min. 2 uL of each sample was injected in splitless mode at 250°C. The GC was coupled to an Agilent 5977A Series MSD with the ion source at 230°C and operated at 70 eV in EI mode scanning from 50 to 850 Da in 0.5 s. Sterols were identified based on retention time and comparison to previously published spectra and laboratory standards. Peaks were integrated in Agilent MassHunter Qualitative Analysis B.06.00 with the default settings of the general integrator except for the start threshold, which was decreased to 0.100.

### SMT sequence alignment and structural analyses

SMT protein sequences were aligned using MAFFT-linsi v7.490 (Katoh & Standley, 2013). Predicted SMT protein structures were generated using ColabFold-AlphaFold2 using MMseqs2 v1.5.2 (Mirdita et al., 2022). The highest ranked structure predictions were aligned and assigned root-mean-square deviation (RMSD) values of atomic coordinates with PyMOL version 2.5.2. Docking was performed with the DockingPie v1.2.1 PyMOL plugin using the Vina function (Rosignoli & Paiardini, 2022) set to 10 poses, an exhaustiveness of 8, and an energy range of 3 without flex. The “Chlamydiae sp., drinking water MAG 1 of 2” SMT ColabFold model was set as the receptor, and 24-methylenecholesterol or SAM were set as the ligands with all torsions but guanidinium and amide and hydrogens added. The search space for 24-methylenecholesterol was set as a 4×4x4 grid centered at Region I, and the best-scoring pose had an affinity of -8.8 kcal/mol. The search space for SAM was set as a 4×4x4 grid centered at Region II, and the best-scoring pose had an affinity of -5.5 kcal/mol.

## Acknowledgments

We thank members of the Welander Lab for helpful discussions. The work (proposal: 503267) conducted by the U.S. Department of Energy Joint Genome Institute (https://ror.org/04xm1d337), a DOE Office of Science User Facility, is supported by the Office of Science of the U.S. Department of Energy operated under Contract No. DE-AC02-05CH11231. Portions of the experiments were performed in the Stanford Geomicrobiology Shared Laboratories Core Facility (RRID:SCR_025000). Funding for this study was provided by the National Science Foundation Grant EAR-1752564 to P.V.W. and the National Science Foundation Graduate Research Fellowship to M.O.B.

## Author Contributions

M.O.B. wrote the manuscript with input from P.V.W. M.O.B. performed the bioinformatics analyses, molecular biology experiments, lipid extractions, and GC-MS analyses. M.O.B. analyzed and interpreted the bioinformatic and GC-MS data. P.V.W. and M.O.B. conceived the project and designed the study.

## Notes

### Competing Interest Statement

The authors have declared no competing interest.

