## Supplemental Info for "Biochemical characterization of bacterial methyltransferases reveals necessary residues for sterol side-chain propylation"

**This file includes:**

Supplementary Tables 1-2
Supplementary Figure 1

**Supplementary Table 1.** Codon-optimized DNA sequences of the "Chlamydiae sp., drinking water MAG 1 of 2" and "*Aplysina aerophoba* meta JGIcombinedJ30088_10000835 1 of 2" SMTs.

| SMT | Codon-optimized sequence |
|---|---|
| Chlamydiae sp., drinking water MAG 1 of 2 | ATGACCAGAGTGAAGGATATGTATACAAAAGAAAAATGCGAAGAGGTGACGA<br>CTTATGTGTCAGCCCGCTCAAAAGTGAAAGATCCCAGTATAACGGAAAAATA<br>CTATGACTTAGTAACCGATTTCTATGAGTTTGGGTGGGGAAAGTCTTTCCAT<br>TTTGCCCCCCAAACCCGTACAGAAGATACCCGTGAAGCACTGGTTCGGCACG<br>AGTTAAAGCTGGCCGATGAGCTCAAACTCAAGCCTGGGATGAAAGTTTTGGA<br>TGTTGGCTGTGGCATAGCGGGTCCGATGAAAAATATTGCGGAAAAGTCCGGT<br>GCTAATATTGTTGGCCTGAACATAAATGCATACCAGATCAAAAAAGCAAAAC<br>AGTATATTAAACGTTGCGGTCTTGAGAACCAGTGTTCTTTCCATCAGGGTAA<br>TTTTATGCATGTCGATTTGCCTGATGCGTCCTTCGATGCCATTTATACAATT<br>GAGGCCACTGTGCATGCACCGGATAGAGTAGCATGTTATAAGGAGATGTATC<br>GCCTGCTGAAACCAGGACATCTGTTCGGGGGTTATGAATACGCCCTGTTAGA<br>CAATTTTGATCCGAAGAATCCAGAGCACATAGCGATTATAGAAGACATGGAG<br>CACGGTGGCGGTCTGCAGAAAGTTCTACCTCTGAAAGAACTCAAACAAACGT<br>TTATCGACGCTGGTTTTAAAGTCTTAGCCTTTGAGGACCTATGTAAAGAAGG<br>TCTGACATGGACACTGCCATTAGAACGCGGCGTCAGGTCCAGCAAGACCGCC<br>CGTGCGCTGACTAATGGTATGGTAAGACTGTTAGAATTCCTGCGTATTGCTC<br>CGAAAGGCAGCACATCCGTATCTAGTTTCCTGAATCTGGGAGCGGATGGCTT<br>CGTAGAGGCGGGTAAGCGCAAACTCTTCACCCCGAACGTTTTTTTTCTGGTG<br>ATGAGACCCTAA |
| *Aplysina aerophoba* meta JGIcombinedJ30088_10000835 1 of 2 | ATGAATCCTACTCCAGCCAAACAGTCCAATAGTCGTTCCCTTGCCGATCTAG<br>CGCCTGACGAAGTGAAGTCAACCGTAGACGGCTATACCGCCATTTATGACGC<br>TGGTCTGGAGTCCCGGAAGGAGAGGTACCAGAGCTTAGTCAACCATTATTAT<br>GATTTAGTTACTGATTTTTTCGAATTCGGCTGGGGGCAGTCCTTCCATTTTG<br>CCCCGCGGCAACGCGGCGAGAGTTTTAAAGCATCACTTCTGAAACACGAACA<br>GTTTCTAGCGGATCGACTCTCTTTAAAGCCCGGCATGCAGGTACTGGATGTT<br>GGCTGTGGCGTCGGCGGTCCGATGGGAGCGCTGGCCCGTTATTCGGGCGCAT<br>CATTTGTCGGCATAAACAACAATGCTTATCAGATCAAAAGGGCCAAACTGCA<br>TACCCGTGATGTCGCCTCACTCTGCCGTTTTATTTGCGGCGATTATATGCAA<br>ATCCCAGAGGGTGACGACCAGTACGATGCTGCGTTCGCAATCGAATCTACCG<br>CGCACGCCCCCGATAAGACCGCAGCGTTTCGCGAAATTTTCAGAGTGCTGCG<br>TCCCGGGGCCTGTTTTGCAGGATATGAATGGTGTCTGACAGACGAGTACGAT<br>CCAGAAAATGCCAAGCATCTGAGGATTAAAAATGATATAATGGTTGGTGGCG<br>GCCTTCCTGATATCGCAGTGACCTCGAAAATTTGCACAGCCTTGAGGACAGC<br>CGGATTTGAACTGTTGGAAGCGTGTGACCGTGCGCCTGAGAGCGATCCCGAA<br>ATGCCGTGGTACCGTGCCCTCCAGGGGCGAGATCTGTCGATGGGTTCCATTC<br>CTCGCACGCCAGTGGGCCGCGCGCTGACCAACTTGGTTTTACGAGTGGGTGA<br>GAAATTGCGCCTGGTTCCCGAAGGTGCTAGAGAGGTTTCCACATTACTGAAC<br>GCGGGAGCTGACGCCCTCGTTGAGGGCGGTAAGTCAGGCATATTTACCCCAA<br>TGTTTTTCTTCCTGGCACGTAAGCCGCGCCGCTCAGAGGATTAA |

**Supplementary Table 2.** Primers used for site-directed mutagenesis of the "Chlamydiae sp., drinking water MAG 1 of 2" and "*Aplysina aerophoba* meta JGIcombinedJ30088_10000835 1 of 2" SMTs.

| Wild type SMT | Mutation | Yeast No. | Primer sequence |
|---|---|---|---|
| Chlamydiae sp., drinking water MAG 1 of 2 | S30E | 68 | CGCTCAAAAGTGAAAGATCCCGAGATAACGGAAAAATACTATGACTTAGTAACC |
| Chlamydiae sp., drinking water MAG 1 of 2 | Y43F | 81 | TACTATGACTTAGTAACCGATTTCTTTGAGTTTGGGTGGGG |
| Chlamydiae sp., drinking water MAG 1 of 2 | F45Y | 83 | ACTTAGTAACCGATTTCTATGAGTATGGGTGGGGAAAGTC |
| Chlamydiae sp., drinking water MAG 1 of 2 | A114D | 152 | CTAATATTGTTGGCCTGAACATAAATGATTACCAGATCAAAAAAGCAAAACAG |
| Chlamydiae sp., drinking water MAG 1 of 2 | Q137D | 175 | TGAGAACCAGTGTTCTTTCCATGATGGTAATTTTATGCATGTCGATTTGC |
| Chlamydiae sp., drinking water MAG 1 of 2 | Q137E | 175 | TGAGAACCAGTGTTCTTTCCATGAGGGTAATTTTATGCATGTCGATTTG |
| Chlamydiae sp., drinking water MAG 1 of 2 | Q137H | 175 | AGAACCAGTGTTCTTTCCATCACGGTAATTTTATGCATGTCGATTTGC |
| Chlamydiae sp., drinking water MAG 1 of 2 | Q137K | 175 | TGAGAACCAGTGTTCTTTCCATAAGGGTAATTTTATGCATGTCGATTTG |
| Chlamydiae sp., drinking water MAG 1 of 2 | H142N | 180 | TGTTCTTTCCATCAGGGTAATTTTATGAATGTCGATTTGCCTGATGC |
| Chlamydiae sp., drinking water MAG 1 of 2 | V166E | 204 | TGCATGCACCGGATAGAGAAGCATGTTATAAGGAGATGTATCGC |
| Chlamydiae sp., drinking water MAG 1 of 2 | I201R | 239 | CCGAAGAATCCAGAGCACAGAGCGATTATAGAAGACATGGAGC |
| Chlamydiae sp., drinking water MAG 1 of 2 | E205A | 243 | CCAGAGCACATAGCGATTATAGCAGACATGGAGCACGG |
| Chlamydiae sp., drinking water MAG 1 of 2 | G211A | 249 | ACATGGAGCACGGTGCCGGTCTGCAGAAAGTTC |
| Chlamydiae sp., drinking water MAG 1 of 2 | G211C | 249 | GACATGGAGCACGGTTGCGGTCTGCAGAAAGTTCTACC |
| Chlamydiae sp., drinking water MAG 1 of 2 | G211D | 249 | GACATGGAGCACGGTGACGGTCTGCAGAAAGTTCTACC |
| Chlamydiae sp., drinking water MAG 1 of 2 | G211F | 249 | GAAGACATGGAGCACGGTTTCGGTCTGCAGAAAGTTCTACC |

| | | | |
|---|---|---|---|
| Chlamydiae sp., drinking water MAG 1 of 2 | G211I | 249 | GAAGACATGGAGCACGGTATCGGTCTGCAGAAAGTTCTA CC |
| Chlamydiae sp., drinking water MAG 1 of 2 | G211L | 249 | GACATGGAGCACGGTCTCGGTCTGCAGAAAGTTCTACC |
| Chlamydiae sp., drinking water MAG 1 of 2 | G211M | 249 | AGACATGGAGCACGGTATGGGTCTGCAGAAAGTTCTACC |
| Chlamydiae sp., drinking water MAG 1 of 2 | G211N | 249 | TATAGAAGACATGGAGCACGGTAACGGTCTGCAGAAAGT TCTACC |
| Chlamydiae sp., drinking water MAG 1 of 2 | G211S | 249 | ATTATAGAAGACATGGAGCACGGTAGCGGTCTGCAGAAA GTTCTAC |
| Chlamydiae sp., drinking water MAG 1 of 2 | G211T | 249 | GAAGACATGGAGCACGGTACCGGTCTGCAGAAAGTTCTA CC |
| Chlamydiae sp., drinking water MAG 1 of 2 | G211V | 249 | AGACATGGAGCACGGTGTCGGTCTGCAGAAAGTTCTACC |
| Chlamydiae sp., drinking water MAG 1 of 2 | T259L | 312 | GGCGTCAGGTCCAGCAAGCTCGCCCGTGCGCTGAC |
| Chlamydiae sp., drinking water MAG 1 of 2 | N265T | 318 | GCCCGTGCGCTGACTACTGGTATGGTAAGACTGTTAGAA TTCCTGC |
| Chlamydiae sp., drinking water MAG 1 of 2 | L290N | 343 | GCACATCCGTATCTAGTTTCCTGAATAATGGAGCGGATG GCTTCG |
| Chlamydiae sp., drinking water MAG 1 of 2 | E297A | 350 | GGAGCGGATGGCTTCGTATTGGCGGGTAAGCGCAA |
| Chlamydiae sp., drinking water MAG 1 of 2 | F309L | 362 | TCACCCCGAACGTTTTATTTCTGGTGATGAGACCC |
| *Aplysina aerophoba* meta JGIcombinedJ30088_1 0000835 1 of 2 | G225D | 249 | AGGATTAAAAATGATATAATGGTTGGTGACGGCCTTCCT GATATCG |

### Supplementary Figure 1

| | | ***S.cerevisiae* #** | **1** | **2** | | | | | | | | | | | | | | | | | | | | | |
|---|---|---|---|---|---|---|---|---|---|---|---|---|---|---|---|---|---|---|---|---|---|---|---|---|---|
| Eukaryotic | (1) | *Saccharomyces cerevisiae* | M | S | - | - | - | - | - | - | - | - | - | - | - | - | - | - | - | - | - | - | - | - | - |
| | | *Trypanosoma brucei* | M | S | A | - | - | - | - | - | - | - | - | - | - | - | - | - | - | - | - | - | - | - | - |
| | | *Arabidopsis thaliana* SMT1 | - | - | - | - | - | - | - | - | - | - | - | - | - | - | - | - | - | - | - | - | - | - | - | - | - |
| | (2) | *Arabidopsis thaliana* SMT2 | M | D | S | L | - | - | - | - | - | - | - | - | - | - | - | - | - | - | - | - | - | T | L | F | F |
| | | *Glycine max* SMT1 | - | - | - | - | - | - | - | - | - | - | - | - | - | - | - | - | - | - | - | - | - | - | - | - | - |
| | | *Chlamydomonas reinhardtii* | M | A | V | A | L | P | A | A | V | T | S | A | Y | E | R | L | A | G | E | F | D | K | L | - | S |
| | | *Pneumocystis jirovecii* | - | - | - | - | - | - | - | - | - | - | - | - | - | - | - | - | - | - | - | - | - | - | - | - | M |
| Bacterial | (3) | Chlamydiae sp., drinking water MAG 1 of2 | - | - | - | - | - | - | - | - | - | - | - | - | - | - | - | - | - | - | - | - | - | - | - | - | - |
| | | *Aplysina aerophoba* meta JGIcombinedJ30088_10000835 1 o | - | - | - | - | - | - | - | - | - | - | - | - | - | - | - | - | - | - | - | - | - | - | - | - | - |
| | | *Aplysina aerophoba* meta Ga0209021_10001462 meta | M | - | - | - | - | - | - | - | - | - | - | - | - | - | - | - | - | - | - | - | - | - | - | - | - |
| | | **Aplysina aerophoba* meta JGIcombinedJ30088_1000330310 | - | - | - | - | - | - | - | - | - | - | - | - | - | - | - | - | - | - | - | - | - | - | - | - | - |
| | (2) | **Aplysina aerophoba* meta Ga0209754_100026449 | M | - | - | - | - | - | - | - | - | - | - | - | - | - | - | - | - | - | - | - | - | - | - | - | - |
| | | *Petrosia ficiformis* meta LXNJ01000389 | M | - | - | - | - | - | - | - | - | - | - | - | - | - | - | - | - | - | - | - | - | - | - | - | - |
| | | **Aplysina aerophoba* meta Ga0209544_100039960 | M | - | - | - | - | - | - | - | - | - | - | - | - | - | - | - | - | - | - | - | - | - | - | - | - |
| | | *Nitrospira* sp., coral reef MAG 1 of2 2682229274 | - | - | - | - | - | - | - | - | - | - | - | - | - | - | - | - | - | - | - | - | - | - | - | - | - |
| | | *Sandaracinus* sp., marine MAG 1 of2 2775843484 | M | T | T | - | - | - | - | - | - | - | - | - | - | - | - | - | - | - | - | - | - | - | - | - | - |
| | | Freshwater, Lake Fryxell, Antarctica meta | M | V | - | - | - | - | - | - | - | - | - | - | - | - | - | - | - | - | - | - | - | - | - | - | - |
| | | Western Arctic Ocean meta Ga0133547_104202862 | M | - | - | - | - | - | - | - | - | - | - | - | - | - | - | - | - | - | - | - | - | - | - | - | - |
| | (1) | **Aplysina aerophoba* meta JGIcombinedJ30088_100127566 | M | - | - | - | - | - | - | - | - | - | - | - | - | - | - | - | - | - | - | - | - | - | - | - | - |
| | | Chlamydiae sp., drinking water MAG 2 of2 2619789261 | - | - | - | - | - | - | - | - | - | - | - | - | - | - | - | - | - | - | - | - | - | - | - | - | - |
| | | *Sandaracinus* sp., marine MAG 2 of2 2775843485 | - | - | - | - | - | - | - | - | - | - | - | - | - | - | - | - | - | - | - | - | - | - | - | - | - |
| | | Marine sediment, Gulf of Thailand meta | M | Q | P | - | - | - | - | - | - | - | - | - | - | - | - | - | - | - | - | - | - | - | - | - | - |
| | | Spirochaetes sp., soil MAG 2709604172 | - | - | - | - | - | - | - | - | - | - | - | - | - | - | - | - | - | - | - | - | - | - | - | - | - |
| | | Hotspring sediment, Dewer Creek, BC meta | M | - | - | - | - | - | - | - | - | - | - | - | - | - | - | - | - | - | - | - | - | - | - | - | - |
| | | Lake sediment, Walker Lake, Nevada meta | M | Q | L | L | - | - | - | - | - | - | - | - | - | - | - | - | - | - | - | - | - | - | - | - | - |
| | | Marine sediment, Helgoland, North Sea meta | M | P | - | - | - | - | - | - | - | - | - | - | - | - | - | - | - | - | - | - | - | - | - | - | - |
| | | *Sarcotragus foetidus* meta LXNI01004480 | M | G | P | R | R | L | A | H | R | D | - | - | - | - | - | - | - | - | - | - | - | - | - | - | L |
| | | *Theonella swinhoei* associated Chromatiales sp. MAG 1 of2 | - | - | - | - | - | - | - | - | - | - | - | - | - | - | - | - | - | - | - | - | - | - | - | - | - |
| | | *Petrosia ficiformis* meta LXNJ01003202 | M | T | - | - | - | - | - | - | - | - | - | - | - | - | - | - | - | - | - | - | - | - | - | - | - |
| | | *Petrosia ficiformis* meta LXNJ01014139 | - | - | - | - | - | - | - | - | - | - | - | - | - | - | - | - | - | - | - | - | - | - | - | - | - |
| | No activity | *Aplysina aerophoba* meta JGIcombinedJ30088_10000835 2 o | M | - | - | - | - | - | - | - | - | - | - | - | - | - | - | - | - | - | - | - | - | - | - | - | - |
| | | *Petrosia ficiformis* meta LXNJ01004423 | M | L | - | - | - | - | - | - | - | - | - | - | - | - | - | - | - | - | - | - | - | - | - | - | - |
| | | *Theonella swinhoei* associated Chromatiales sp. MAG 2 of2 | - | - | - | - | - | - | - | - | - | - | - | - | - | - | - | - | - | - | - | - | - | - | - | - | - |
| | | Hotspring, Beatty, Nevada meta_Ga0114945_100210811 | M | R | - | - | - | - | - | - | - | - | - | - | - | - | - | - | - | - | - | - | - | - | - | - | - |
| | | *Nitrospira* sp., coral reef MAG 2 of2 2682229270 | L | - | - | - | - | - | - | - | - | - | - | - | - | - | - | - | - | - | - | - | - | - | - | - | - |
| | | **Aplysina aerophoba* meta JGIcombinedJ30088_100153964 | M | - | - | - | - | - | - | - | - | - | - | - | - | - | - | - | - | - | - | - | - | - | - | - | - |
| | | **Aplysina aerophoba meta* JGIcombinedJ30088_100077196 | M | - | - | - | - | - | - | - | - | - | - | - | - | - | - | - | - | - | - | - | - | - | - | - | - |
| | | **Aplysina aerophoba meta* JGIcombinedJ30088_100209763 | M | - | - | - | - | - | - | - | - | - | - | - | - | - | - | - | - | - | - | - | - | - | - | - | - |
| | | **Aplysina aerophoba meta* JGIcombinedJ30088_100057164 | L | - | - | - | - | - | - | - | - | - | - | - | - | - | - | - | - | - | - | - | - | - | - | - | - |
| | | **Sarcotragus foetidus* meta LXNI01006544 | M | D | - | - | - | - | - | - | - | - | - | - | - | - | - | - | - | - | - | - | - | - | - | - | - |
| | | **Sarcotragus foetidus* meta LXNI01016685 | M | - | - | - | - | - | - | - | - | - | - | - | - | - | - | - | - | - | - | - | - | - | - | - | - |
| | | Consensus | M | - | - | - | - | - | - | - | - | - | - | - | - | - | - | - | - | - | - | - | - | - | - | - | - |
| | | **Consensus #** | **1** | **2** | **3** | **4** | **5** | **6** | **7** | **8** | **9** | **10** | **11** | **12** | **13** | **14** | **15** | **16** | **17** | **18** | **19** | **20** | **21** | **22** | **23** | **24** | **25** |

| | | *S.cerevisiae* # | 3 | 4 | 5 | 6 | 7 | 8 | | | | | | | | | | 9 | 10 | 11 | 12 | 13 | 14 | | | | |
|---|---|---|---|---|---|---|---|---|---|---|---|---|---|---|---|---|---|---|---|---|---|---|---|---|---|---|---|
| Eukaryotic | (1) | *Saccharomyces cerevisiae* | E | T | E | L | R | K | - | - | - | - | - | - | - | - | - | R | Q | A | Q | F | T | - | - | - | - |
| | | *Trypanosoma brucei* | - | - | - | - | - | - | - | G | S | R | - | - | - | - | - | - | - | - | - | - | - | - | - | - | - |
| | | *Arabidopsis thaliana* SMT1 | - | - | - | - | - | - | - | - | - | - | - | - | - | - | - | - | - | - | - | - | - | - | - | - | - |
| | (2) | *Arabidopsis thaliana* SMT2 | T | G | A | L | V | A | V | G | I | - | - | - | - | - | - | - | - | Y | W | F | L | C | - | - | - |
| | | *Glycine max* SMT1 | - | - | - | - | - | - | - | - | - | - | - | - | - | - | - | - | - | - | - | - | - | - | - | - | - |
| | | *Chlamydomonas reinhardtii* | T | T | Q | K | Y | A | V | G | I | - | A | G | G | V | T | S | L | Y | L | L | A | K | - | - | - |
| | | *Pneumocystis jirovecii* | T | F | E | L | E | P | - | - | - | - | - | L | D | I | E | K | D | Y | E | F | S | - | - | - | - |
| Bacterial | (3) | Chlamydiae sp., drinking water MAG 1 of2 | - | - | - | - | - | - | - | - | - | - | - | - | - | - | - | - | - | - | - | - | - | - | - | - | - |
| | | *Aplysina aerophoba* meta JGIcombinedJ30088_10000835 1 of | - | - | - | - | - | - | - | - | - | - | - | - | - | - | - | - | - | - | - | - | - | - | - | - | - |
| | | *Aplysina aerophoba* meta Ga0209021_10001462 meta | - | - | - | - | - | - | - | - | - | - | - | - | - | - | - | - | - | - | - | - | - | - | - | - | - |
| | | **Aplysina aerophoba* meta JGIcombinedJ30088_1000330310 | - | - | - | - | - | - | - | - | - | - | - | - | - | - | - | - | - | - | - | - | - | - | - | - | - |
| | (2) | **Aplysina aerophoba* meta Ga0209754_100026449 | - | - | - | - | - | - | - | - | - | - | - | - | - | - | - | - | - | - | - | - | - | - | - | - | - |
| | | *Petrosia ficiformis* meta LXNJ01000389 | - | - | - | - | - | - | - | - | - | - | - | - | - | - | - | - | - | - | - | - | - | - | - | - | - |
| | | **Aplysina aerophoba* meta Ga0209544_100039960 | - | - | - | - | - | - | - | - | - | - | - | - | - | - | - | - | - | - | - | - | - | - | - | - | - |
| | | *Nitrospira* sp., coral reef MAG 1 of2 2682229274 | - | - | - | - | - | - | - | - | - | - | - | - | - | - | - | - | - | - | - | - | - | - | - | - | - |
| | | *Sandaracinus* sp., marine MAG 1 of2 2775843484 | - | - | - | - | - | - | - | - | - | - | - | - | - | - | - | - | - | - | - | - | - | - | - | - | - |
| | | Freshwater, Lake Fryxell, Antarctica meta | - | - | - | - | - | - | - | - | - | - | - | - | - | - | - | Q | L | N | E | M | R | A | - | - | - |
| | | Western Arctic Ocean meta Ga0133547_104202862 | - | - | - | - | - | - | - | - | - | - | - | - | - | - | - | - | - | - | - | - | - | - | - | - | - |
| | (1) | **Aplysina aerophoba* meta JGIcombinedJ30088_100127566 | - | - | - | - | - | - | - | - | - | - | - | - | - | - | - | - | - | - | - | - | - | - | - | - | - |
| | | Chlamydiae sp., drinking water MAG 2 of2 2619789261 | - | - | - | - | - | - | - | - | - | - | - | - | - | - | - | - | - | - | - | - | - | - | - | - | - |
| | | *Sandaracinus* sp., marine MAG 2 of2 2775843485 | - | - | - | - | - | - | - | - | - | - | - | - | - | - | - | - | - | - | - | - | - | - | - | - | - |
| | | Marine sediment, Gulf of Thailand meta | - | - | - | - | - | - | - | - | - | - | - | - | - | - | - | - | - | - | - | - | - | - | - | - | - |
| | | Spirochaetes sp., soil MAG 2709604172 | - | - | - | - | - | - | - | - | - | - | - | - | - | - | - | - | - | - | - | - | - | - | - | - | - |
| | | Hotspring sediment, Dewer Creek, BC meta | - | - | - | - | - | - | - | - | - | - | - | - | - | - | - | - | - | - | - | - | - | - | - | - | - |
| | | Lake sediment, Walker Lake, Nevada meta | - | - | - | - | - | - | - | - | - | - | - | - | - | - | - | - | - | - | - | - | - | - | - | - | - |
| | | Marine sediment, Helgoland, North Sea meta | - | - | - | - | - | - | - | - | - | - | - | - | - | - | - | - | - | - | - | - | - | - | - | - | - |
| | | *Sarcotragus foetidus* meta LXNI01004480 | Q | D | E | R | R | R | - | - | - | - | A | L | G | L | A | Q | E | R | A | F | R | V | S | C | K |
| | | *Theonella swinhoei* associated Chromatiales sp. MAG 1 of2 | - | - | - | - | - | - | - | - | - | - | - | - | - | - | - | - | - | - | - | - | - | - | - | - | - |
| | | *Petrosia ficiformis* meta LXNJ01003202 | - | - | - | - | - | - | - | - | - | - | - | - | - | - | - | - | - | N | T | F | R | G | - | - | - |
| | | *Petrosia ficiformis* meta LXNJ01014139 | - | - | - | - | - | - | - | - | - | - | - | - | - | - | - | - | - | - | - | - | - | - | - | - | - |
| | No activity | *Aplysina aerophoba* meta JGIcombinedJ30088_10000835 2 of | - | - | - | - | - | - | - | - | - | - | - | - | - | - | - | - | - | - | - | - | - | - | - | - | - |
| | | *Petrosia ficiformis* meta LXNJ01004423 | - | - | - | - | - | - | - | - | - | - | - | - | - | - | - | - | - | - | - | - | - | - | - | - | - |
| | | *Theonella swinhoei* associated Chromatiales sp. MAG 2 of2 | - | - | - | - | - | - | - | - | - | - | - | - | - | - | - | - | - | - | - | - | - | - | - | - | - |
| | | Hotspring, Beatty, Nevada meta_Ga0114945_100210811 | - | - | - | - | - | - | - | - | - | - | - | - | - | - | - | - | - | - | - | - | - | - | - | - | - |
| | | *Nitrospira* sp., coral reef MAG 2 of2 2682229270 | - | - | - | - | - | - | - | - | - | - | - | - | - | - | - | - | - | - | - | - | - | - | - | - | - |
| | | **Aplysina aerophoba* meta JGIcombinedJ30088_100153964 | - | - | - | - | - | - | - | - | - | - | - | - | - | - | - | - | - | - | - | - | - | - | - | - | - |
| | | **Aplysina aerophoba meta* JGIcombinedJ30088_100077196 | - | - | - | - | - | - | - | - | - | - | - | - | - | - | - | - | - | - | - | - | - | - | - | - | - |
| | | **Aplysina aerophoba meta* JGIcombinedJ30088_100209763 | - | - | - | - | - | - | - | - | - | - | - | - | - | - | - | - | - | - | - | - | - | - | - | - | - |
| | | **Aplysina aerophoba meta* JGIcombinedJ30088_100057164 | - | - | - | - | - | - | - | - | - | - | - | - | - | - | - | - | - | - | - | - | - | - | - | - | - |
| | | **Sarcotragus foetidus* meta LXNI01006544 | - | - | - | - | - | - | - | - | - | - | - | - | - | - | - | - | - | - | - | - | - | - | - | - | - |
| | | **Sarcotragus foetidus* meta LXNI01016685 | - | - | - | - | - | - | - | - | - | - | - | - | - | - | - | - | - | - | - | - | - | - | - | - | - |
| | | Consensus | - | - | - | - | - | - | - | - | - | - | - | - | - | - | - | - | - | - | - | - | - | - | - | - | - |
| | | **Consensus #** | **26** | **27** | **28** | **29** | **30** | **31** | **32** | **33** | **34** | **35** | **36** | **37** | **38** | **39** | **40** | **41** | **42** | **43** | **44** | **45** | **46** | **47** | **48** | **49** | **50** |

| | | *S.cerevisiae* # | | | | | | | | | | | | | | | | | | | | | | | | |
|---|---|---|---|---|---|---|---|---|---|---|---|---|---|---|---|---|---|---|---|---|---|---|---|---|---|---|
| Eukaryotic | (1) | *Saccharomyces cerevisiae* | - | - | - | - | - | - | - | - | - | - | - | - | - | - | - | - | - | - | - | - | - | - | - | - | - |
| | | *Trypanosoma brucei* | - | - | - | - | - | - | - | - | - | - | - | - | - | - | - | - | - | - | - | - | - | - | - | - | - |
| | | *Arabidopsis thaliana* SMT1 | - | - | - | - | - | - | - | - | - | - | - | - | - | - | - | - | - | - | - | - | - | - | - | - | - |
| | (2) | *Arabidopsis thaliana* SMT2 | - | - | - | - | - | - | - | - | - | - | - | - | - | - | - | - | - | - | - | - | - | - | - | - | - |
| | | *Glycine max* SMT1 | - | - | - | - | - | - | - | - | - | - | - | - | - | - | - | - | - | - | - | - | - | - | - | - | - |
| | | *Chlamydomonas reinhardtii* | - | - | - | - | - | - | - | - | - | - | - | - | - | - | - | - | - | - | - | - | - | - | - | - | - |
| | | *Pneumocystis jirovecii* | - | - | - | - | - | - | - | - | - | - | - | - | - | - | - | - | - | - | - | - | - | - | - | - | - |
| Bacterial | (3) | Chlamydiae sp., drinking water MAG 1 of2 | - | - | - | - | - | - | - | - | - | - | - | - | - | - | - | - | - | - | - | - | - | - | - | - | - |
| | | *Aplysina aerophoba* meta JGIcombinedJ30088_10000835 1 of | - | - | - | - | - | - | - | - | - | - | - | - | - | - | - | - | - | - | - | - | - | - | - | - | - |
| | | *Aplysina aerophoba* meta Ga0209021_10001462 meta | - | - | - | - | - | - | - | - | - | - | - | - | - | - | - | - | - | - | - | - | - | - | - | - | - |
| | | **Aplysina aerophoba* meta JGIcombinedJ30088_1000330310 | - | - | - | - | - | - | - | - | - | - | - | - | - | - | - | - | - | - | - | - | - | - | - | - | - |
| | (2) | **Aplysina aerophoba* meta Ga0209754_100026449 | - | - | - | - | - | - | - | - | - | - | - | - | - | - | - | - | - | - | - | - | - | - | - | - | - |
| | | *Petrosia ficiformis* meta LXNJ01000389 | - | - | - | - | - | - | - | - | - | - | - | - | - | - | - | - | - | - | - | - | - | - | - | - | - |
| | | **Aplysina aerophoba* meta Ga0209544_100039960 | - | - | - | - | - | - | - | - | - | - | - | - | - | - | - | - | - | - | - | - | - | - | - | - | - |
| | | *Nitrospira* sp., coral reef MAG 1 of2 2682229274 | - | - | - | - | - | - | - | - | - | - | - | - | - | - | - | - | - | - | - | - | - | - | - | - | - |
| | | *Sandaracinus* sp., marine MAG 1 of2 2775843484 | - | - | - | - | - | - | - | - | - | - | - | - | - | - | - | - | - | - | - | - | - | - | - | - | - |
| | | Freshwater, Lake Fryxell, Antarctica meta | - | - | - | - | - | - | - | - | - | - | - | - | - | - | - | - | - | - | - | - | - | - | - | - | - |
| | | Western Arctic Ocean meta Ga0133547_104202862 | - | - | - | - | - | - | - | - | - | - | - | - | - | - | - | - | - | - | - | - | - | - | - | - | - |
| | (1) | **Aplysina aerophoba* meta JGIcombinedJ30088_100127566 | - | - | - | - | - | - | - | - | - | - | - | - | - | - | - | - | - | - | - | - | - | - | - | - | - |
| | | Chlamydiae sp., drinking water MAG 2 of2 2619789261 | - | - | - | - | - | - | - | - | - | - | - | - | - | - | - | - | - | - | - | - | - | - | - | - | - |
| | | *Sandaracinus* sp., marine MAG 2 of2 2775843485 | - | - | - | - | - | - | - | - | - | - | - | - | - | - | - | - | - | - | - | - | - | - | - | - | - |
| | | Marine sediment, Gulf of Thailand meta | - | - | - | - | - | - | - | - | - | - | - | - | - | - | - | - | - | - | - | - | - | - | - | - | - |
| | | Spirochaetes sp., soil MAG 2709604172 | - | - | - | - | - | - | - | - | - | - | - | - | - | - | - | - | - | - | - | - | - | - | - | - | - |
| | | Hotspring sediment, Dewer Creek, BC meta | - | - | - | - | - | - | - | - | - | - | - | - | - | - | - | - | - | - | - | - | - | - | - | - | - |
| | | Lake sediment, Walker Lake, Nevada meta | - | - | - | - | - | - | - | - | - | - | - | - | - | - | - | - | - | - | - | - | - | - | - | - | - |
| | | Marine sediment, Helgoland, North Sea meta | - | - | - | - | - | - | - | - | - | - | - | - | - | - | - | - | - | - | - | - | - | - | - | - | - |
| | | *Sarcotragus foetidus* meta LXNI01004480 | F | A | V | P | A | A | G | K | S | D | E | C | H | E | A | G | C | C | G | V | L | D | L | S | D |
| | | *Theonella swinhoei* associated Chromatiales sp. MAG 1 of2 | - | - | - | - | - | - | - | - | - | - | - | - | - | - | - | - | - | - | - | - | - | - | - | - | - |
| | | *Petrosia ficiformis* meta LXNJ01003202 | - | - | - | - | - | - | - | - | - | - | - | - | - | - | - | - | - | - | - | - | - | - | - | - | - |
| | | *Petrosia ficiformis* meta LXNJ01014139 | - | - | - | - | - | - | - | - | - | - | - | - | - | - | - | - | - | - | - | - | - | - | - | - | - |
| | No activity | *Aplysina aerophoba* meta JGIcombinedJ30088_10000835 2 of | - | - | - | - | - | - | - | - | - | - | - | - | - | - | - | - | - | - | - | - | - | - | - | - | - |
| | | *Petrosia ficiformis* meta LXNJ01004423 | - | - | - | - | - | - | - | - | - | - | - | - | - | - | - | - | - | - | - | - | - | - | - | - | - |
| | | *Theonella swinhoei* associated Chromatiales sp. MAG 2 of2 | - | - | - | - | - | - | - | - | - | - | - | - | - | - | - | - | - | - | - | - | - | - | - | - | - |
| | | Hotspring, Beatty, Nevada meta_Ga0114945_100210811 | - | - | - | - | - | - | - | - | - | - | - | - | - | - | - | - | - | - | - | - | - | - | - | - | - |
| | | *Nitrospira* sp., coral reef MAG 2 of2 2682229270 | - | - | - | - | - | - | - | - | - | - | - | - | - | - | - | - | - | - | - | - | - | - | - | - | - |
| | | **Aplysina aerophoba* meta JGIcombinedJ30088_100153964 | - | - | - | - | - | - | - | - | - | - | - | - | - | - | - | - | - | - | - | - | - | - | - | - | - |
| | | **Aplysina aerophoba meta* JGIcombinedJ30088_100077196 | - | - | - | - | - | - | - | - | - | - | - | - | - | - | - | - | - | - | - | - | - | - | - | - | - |
| | | **Aplysina aerophoba meta* JGIcombinedJ30088_100209763 | - | - | - | - | - | - | - | - | - | - | - | - | - | - | - | - | - | - | - | - | - | - | - | - | - |
| | | **Aplysina aerophoba meta* JGIcombinedJ30088_100057164 | - | - | - | - | - | - | - | - | - | - | - | - | - | - | - | - | - | - | - | - | - | - | - | - | - |
| | | **Sarcotragus foetidus* meta LXNI01006544 | - | - | - | - | - | - | - | - | - | - | - | - | - | - | - | - | - | - | - | - | - | - | - | - | - |
| | | **Sarcotragus foetidus* meta LXNI01016685 | - | - | - | - | - | - | - | - | - | - | - | - | - | - | - | - | - | - | - | - | - | - | - | - | - |
| | | Consensus | - | - | - | - | - | - | - | - | - | - | - | - | - | - | - | - | - | - | - | - | - | - | - | - | - |
| | | **Consensus #** | **51** | **52** | **53** | **54** | **55** | **56** | **57** | **58** | **59** | **60** | **61** | **62** | **63** | **64** | **65** | **66** | **67** | **68** | **69** | **70** | **71** | **72** | **73** | **74** | **75** |

| | | S.cerevisiae # | | | | | | | | | | 15 | 16 | 17 | 18 | 19 | 20 | 21 | 22 | 23 | 24 | 25 | 26 | 27 | 28 | 29 | 30 |
|---|---|---|---|---|---|---|---|---|---|---|---|---|---|---|---|---|---|---|---|---|---|---|---|---|---|---|---|
| Eukaryotic | (1) | *Saccharomyces cerevisiae* | – | – | – | – | – | – | – | – | – | R | E | L | H | G | D | D | I | G | K | K | T | G | L | S | A |
| | | *Trypanosoma brucei* | – | – | – | – | – | – | – | – | – | – | – | – | – | – | G | P | L | S | L | L | I | A | R | E | R |
| | | *Arabidopsis thaliana* SMT1 | – | – | – | – | – | – | – | – | – | – | – | – | – | – | – | – | – | – | M | D | L | A | S | N | L |
| | (2) | *Arabidopsis thaliana* SMT2 | – | – | – | – | – | – | – | – | – | V | L | G | P | A | E | R | K | G | K | R | A | V | D | L | S |
| | | *Glycine max* SMT1 | – | – | – | – | – | – | – | – | – | – | – | – | – | – | – | – | – | – | M | D | L | A | S | N | L |
| | | *Chlamydomonas reinhardtii* | – | – | – | – | – | – | – | – | – | V | L | K | G | S | D | R | D | K | P | T | T | L | Q | L | S |
| | | *Pneumocystis jirovecii* | – | – | – | – | – | – | – | – | – | K | A | M | H | G | S | D | A | K | S | K | H | G | I | C | A |
| Bacterial | (3) | Chlamydiae sp., drinking water MAG 1 of2 | – | – | – | – | – | – | – | – | – | – | – | – | – | – | – | – | – | M | T | R | V | K | D | M | Y |
| | | *Aplysina aerophoba* meta JGIcombinedJ30088_10000835 1 of | – | – | – | – | – | – | – | – | – | – | – | – | M | N | P | T | P | A | K | Q | S | N | S | R | S |
| | | *Aplysina aerophoba* meta Ga0209021_10001462 meta | – | – | – | – | – | – | – | – | – | – | – | – | – | – | – | – | – | P | F | R | L | S | R | T | M |
| | | **Aplysina aerophoba* meta JGIcombinedJ30088_1000330310 | – | – | – | – | – | – | – | – | – | – | – | – | – | – | – | – | – | V | T | R | L | T | D | I | L |
| | (2) | **Aplysina aerophoba* meta Ga0209754_100026449 | – | – | – | – | – | – | – | – | – | – | – | – | – | – | – | – | – | A | V | K | L | T | D | I | L |
| | | *Petrosia ficiformis* meta LXNJ01000389 | – | – | – | – | – | – | – | – | – | – | – | – | – | – | – | – | – | A | V | K | I | T | R | A | L |
| | | **Aplysina aerophoba* meta Ga0209544_100039960 | – | – | – | – | – | – | – | – | – | – | – | – | – | – | – | – | – | A | V | K | L | S | D | V | L |
| | | *Nitrospira* sp., coral reef MAG 1 of2 2682229274 | – | – | – | – | – | – | – | – | – | – | – | – | M | S | R | T | P | A | K | R | S | N | S | R | S |
| | | *Sandaracinus* sp., marine MAG 1 of2 2775843484 | – | – | – | – | – | – | – | – | – | Q | P | F | E | P | F | P | F | H | P | A | D | P | A | D | P |
| | | Freshwater, Lake Fryxell, Antarctica meta | – | – | – | – | – | – | – | – | – | V | P | A | H | E | D | S | A | Q | L | R | S | P | T | E | S |
| | | Western Arctic Ocean meta Ga0133547_104202862 | – | – | – | – | – | – | – | – | – | – | – | – | – | – | – | – | A | R | P | N | V | T | G | A | A |
| | (1) | **Aplysina aerophoba* meta JGIcombinedJ30088_100127566 | – | – | – | – | – | – | – | – | – | – | – | – | – | – | – | – | – | P | T | R | L | S | Q | I | L |
| | | Chlamydiae sp., drinking water MAG 2 of2 2619789261 | – | – | – | – | – | – | – | – | – | – | – | – | – | – | – | – | – | M | P | R | V | K | D | V | Y |
| | | *Sandaracinus* sp., marine MAG 2 of2 2775843485 | – | – | – | – | – | – | – | – | – | – | – | – | – | – | – | – | – | – | – | – | – | – | – | – | – |
| | | Marine sediment, Gulf of Thailand meta | – | – | – | – | – | – | – | – | – | – | – | – | – | – | – | – | – | – | – | – | V | K | A | N | L |
| | | Spirochaetes sp., soil MAG 2709604172 | – | – | – | – | – | – | – | – | – | – | – | – | – | – | – | – | – | M | P | S | I | K | N | P | I |
| | | Hotspring sediment, Dewer Creek, BC meta | – | – | – | – | – | – | – | – | – | – | – | – | – | – | – | – | K | R | R | S | V | F | S | F | L |
| | | Lake sediment, Walker Lake, Nevada meta | – | – | – | – | – | – | – | – | – | – | – | – | – | – | – | – | – | – | – | – | – | – | – | R | Q |
| | | Marine sediment, Helgoland, North Sea meta | – | – | – | – | – | – | – | – | – | – | – | – | – | – | Q | A | Q | A | V | R | N | T | T | F | Y |
| | | *Sarcotragus foetidus* meta LXNI01004480 | L | G | E | T | T | R | L | C | H | N | I | L | M | M | R | S | S | A | K | R | S | G | S | R | S |
| | | *Theonella swinhoei* associated Chromatiales sp. MAG 1 of2 | – | – | – | – | – | – | – | – | – | – | – | – | M | S | R | T | S | T | K | Q | S | S | S | R | S |
| | | *Petrosia ficiformis* meta LXNJ01003202 | – | – | – | – | – | – | – | – | – | – | – | – | V | A | L | E | S | I | R | T | K | R | S | R | S |
| | | *Petrosia ficiformis* meta LXNJ01014139 | – | – | – | – | – | – | – | – | – | – | – | – | M | N | R | P | A | A | T | R | S | K | A | R | S |
| | No activity | *Aplysina aerophoba* meta JGIcombinedJ30088_10000835 2 of | – | – | – | – | – | – | – | – | – | – | – | – | – | – | – | – | Q | L | G | K | L | L | H | A | L |
| | | *Petrosia ficiformis* meta LXNJ01004423 | – | – | – | – | – | – | – | – | – | P | A | L | A | E | N | A | M | L | T | T | L | S | R | S | L |
| | | *Theonella swinhoei* associated Chromatiales sp. MAG 2 of2 | – | – | – | – | – | – | – | – | – | – | – | – | – | – | – | – | M | L | T | E | L | P | S | S | V |
| | | Hotspring, Beatty, Nevada meta_Ga0114945_100210811 | – | – | – | – | – | – | – | – | – | – | – | – | – | – | – | – | A | D | P | R | L | T | D | L | V |
| | | *Nitrospira* sp., coral reef MAG 2 of2 2682229270 | – | – | – | – | – | – | – | – | – | – | – | – | – | – | – | – | E | L | G | K | L | Q | H | A | L |
| | | **Aplysina aerophoba* meta JGIcombinedJ30088_100153964 | – | – | – | – | – | – | – | – | – | – | – | – | – | – | – | – | – | A | S | R | L | T | D | I | L |
| | | **Aplysina aerophoba meta* JGIcombinedJ30088_100077196 | – | – | – | – | – | – | – | – | – | – | – | – | – | – | – | – | – | S | F | K | L | S | N | L | L |
| | | **Aplysina aerophoba meta* JGIcombinedJ30088_100209763 | – | – | – | – | – | – | – | – | – | – | – | – | – | – | – | – | – | A | S | K | L | S | G | I | M |
| | | **Aplysina aerophoba meta* JGIcombinedJ30088_100057164 | – | – | – | – | – | – | – | – | – | – | – | – | – | – | – | – | – | A | P | R | L | S | R | I | L |
| | | **Sarcotragus foetidus* meta LXNI01006544 | – | – | – | – | – | – | – | – | – | I | P | S | L | V | R | P | M | G | E | K | L | L | R | L | F |
| | | **Sarcotragus foetidus* meta LXNI01016685 | – | – | – | – | – | – | – | – | – | – | – | – | – | – | – | – | – | S | S | K | L | T | R | V | L |
| | | Consensus | – | – | – | – | – | – | – | – | – | – | – | – | – | – | – | – | – | A | P | R | L | X | S | R | L |
| | | **Consensus #** | **76** | **77** | **78** | **79** | **80** | **81** | **82** | **83** | **84** | **85** | **86** | **87** | **88** | **89** | **90** | **91** | **92** | **93** | **94** | **95** | **96** | **97** | **98** | **99** | **100** |

| | | S.cerevisiae # | 31 | 32 | 33 | 34 | 35 | 36 | 37 | 38 | 39 | 40 | 41 | 42 | 43 | 44 | 45 | 46 | 47 | 48 | 49 | 50 | 51 | | | | |
|---|---|---|---|---|---|---|---|---|---|---|---|---|---|---|---|---|---|---|---|---|---|---|---|---|---|---|---|
| **Eukaryotic** | (1) | *Saccharomyces cerevisiae* | L | M | S | K | N | N | S | A | Q | K | E | A | V | Q | K | Y | L | R | N | W | D | - | - | - | - |
| | | *Trypanosoma brucei* | D | A | N | G | V | N | G | D | V | N | A | T | A | G | R | L | R | D | R | Y | - | - | - | - | - |
| | | *Arabidopsis thaliana* SMT1 | G | G | K | I | D | K | S | D | V | L | T | A | V | E | K | Y | E | Q | Y | H | - | - | - | - | - |
| | (2) | *Arabidopsis thaliana* SMT2 | G | G | S | I | S | A | E | K | V | Q | D | N | Y | K | Q | Y | W | S | F | F | - | - | - | - | - |
| | | *Glycine max* SMT1 | G | G | K | I | D | K | A | E | V | L | S | A | V | Q | K | Y | E | K | Y | H | - | - | - | - | - |
| | | *Chlamydomonas reinhardtii* | G | G | S | I | D | S | S | K | V | K | D | E | F | T | A | Y | A | D | S | Y | - | - | - | - | - |
| | | *Pneumocystis jirovecii* | L | F | N | K | N | R | D | A | Q | K | V | A | F | D | S | Y | F | G | F | W | G | D | K | Y | L |
| **Bacterial** | (3) | Chlamydiae sp., drinking water MAG 1 of2 | T | K | E | K | C | E | E | - | - | - | - | - | V | T | T | Y | V | S | A | R | - | - | - | - | - |
| | | *Aplysina aerophoba* meta JGIcombinedJ30088_10000835 1 of | L | A | D | L | A | P | D | E | V | K | S | T | V | D | G | Y | T | A | I | Y | - | - | - | - | - |
| | | *Aplysina aerophoba* meta Ga0209021_10001462 meta | K | D | N | S | G | T | S | G | V | A | P | E | I | S | E | Y | E | E | R | F | - | - | - | - | - |
| | | **Aplysina aerophoba* meta JGIcombinedJ30088_1000330310 | K | G | A | K | S | V | D | S | V | E | A | S | A | Q | E | Y | V | K | R | Y | - | - | - | - | - |
| | (2) | **Aplysina aerophoba* meta Ga0209754_100026449 | K | D | A | K | S | G | D | G | V | A | S | R | A | E | D | Y | D | R | R | Y | - | - | - | - | - |
| | | *Petrosia ficiformis* meta LXNJ01000389 | R | N | R | L | S | S | G | G | V | A | P | A | M | R | S | Y | E | D | R | H | - | - | - | - | - |
| | | **Aplysina aerophoba* meta Ga0209544_100039960 | K | E | K | Q | Q | T | D | G | V | A | P | A | V | N | E | Y | E | R | G | F | - | - | - | - | - |
| | | *Nitrospira* sp., coral reef MAG 1 of2 2682229274 | L | A | D | L | T | P | D | E | V | K | S | A | V | D | G | Y | T | A | I | Y | - | - | - | - | - |
| | | *Sandaracinus* sp., marine MAG 1 of2 2775843484 | R | P | G | H | A | R | G | D | I | R | E | V | V | H | T | Y | R | G | F | H | - | - | - | - | - |
| | | Freshwater, Lake Fryxell, Antarctica meta | R | A | A | V | T | R | A | D | V | R | G | D | I | E | Q | Y | R | S | F | H | - | - | - | - | - |
| | | Western Arctic Ocean meta Ga0133547_104202862 | R | T | G | V | D | A | G | E | V | Q | G | T | V | D | A | Y | R | D | L | Y | - | - | - | - | - |
| | (1) | **Aplysina aerophoba* meta JGIcombinedJ30088_100127566 | K | R | S | L | D | P | D | D | V | E | S | S | V | G | A | H | E | R | S | F | - | - | - | - | - |
| | | Chlamydiae sp., drinking water MAG 2 of2 2619789261 | T | K | E | K | C | E | E | - | - | - | - | - | V | T | T | Y | V | S | A | K | - | - | - | - | - |
| | | *Sandaracinus* sp., marine MAG 2 of2 2775843485 | M | S | G | V | S | R | E | S | F | G | E | T | V | S | Q | Y | R | G | L | H | - | - | - | - | - |
| | | Marine sediment, Gulf of Thailand meta | R | V | G | A | N | G | S | A | P | A | R | T | L | G | S | Y | L | K | T | H | - | - | - | - | - |
| | | Spirochaetes sp., soil MAG 2709604172 | P | K | G | V | E | K | E | N | L | A | S | M | I | H | E | Y | R | G | F | H | - | - | - | - | - |
| | | Hotspring sediment, Dewer Creek, BC meta | R | S | A | H | Q | G | R | G | V | E | R | A | V | R | R | H | R | A | H | Y | - | - | - | - | - |
| | | Lake sediment, Walker Lake, Nevada meta | S | S | T | V | G | G | D | D | I | R | A | Q | V | S | R | Y | R | S | L | H | - | - | - | - | - |
| | | Marine sediment, Helgoland, North Sea meta | T | D | N | D | K | R | Q | R | V | G | G | T | I | R | E | Y | M | G | F | F | - | - | - | - | - |
| | | *Sarcotragus foetidus* meta LXNI01004480 | L | S | N | L | A | P | D | E | V | K | S | V | V | E | G | Y | T | A | I | Y | - | - | - | - | - |
| | | *Theonella swinhoei* associated Chromatiales sp. MAG 1 of2 | F | S | D | L | T | P | D | E | V | T | Q | A | V | D | Q | Y | T | G | L | H | - | - | - | - | - |
| | | *Petrosia ficiformis* meta LXNJ01003202 | L | G | N | L | N | R | D | E | V | K | S | A | V | D | G | Y | N | V | L | H | - | - | - | - | - |
| | | *Petrosia ficiformis* meta LXNJ01014139 | L | G | D | I | A | A | D | D | V | K | S | T | V | D | G | Y | T | E | I | Y | - | - | - | - | - |
| | No activity | *Aplysina aerophoba* meta JGIcombinedJ30088_10000835 2 of | K | E | A | Y | R | E | G | D | V | T | R | R | V | E | H | F | D | G | W | I | - | - | - | - | - |
| | | *Petrosia ficiformis* meta LXNJ01004423 | R | S | A | Y | K | G | R | N | V | A | S | R | V | V | R | F | G | G | W | L | - | - | - | - | - |
| | | *Theonella swinhoei* associated Chromatiales sp. MAG 2 of2 | K | K | A | L | R | G | W | D | I | A | N | R | V | K | D | Y | N | G | W | L | - | - | - | - | - |
| | | Hotspring, Beatty, Nevada meta_Ga0114945_100210811 | Q | T | S | L | P | R | P | E | V | Q | R | A | A | A | Q | Y | L | A | L | H | - | - | - | - | - |
| | | *Nitrospira* sp., coral reef MAG 2 of2 2682229270 | K | E | A | Y | R | G | E | D | V | T | R | Q | V | Q | H | F | D | G | W | I | - | - | - | - | - |
| | | **Aplysina aerophoba* meta JGIcombinedJ30088_100153964 | K | E | A | K | S | G | D | R | V | D | T | A | V | R | D | Y | D | H | R | Y | - | - | - | - | - |
| | | **Aplysina aerophoba meta* JGIcombinedJ30088_100077196 | S | K | L | D | N | Q | I | G | V | D | K | A | F | E | N | Y | S | N | R | E | - | - | - | - | - |
| | | **Aplysina aerophoba meta* JGIcombinedJ30088_100209763 | K | N | A | L | K | R | D | A | V | A | S | T | A | R | S | Y | E | H | V | Y | - | - | - | - | - |
| | | **Aplysina aerophoba meta* JGIcombinedJ30088_100057164 | R | N | A | V | D | G | G | D | V | A | A | A | I | Q | E | Y | E | Y | R | Y | - | - | - | - | - |
| | | **Sarcotragus foetidus* meta LXNI01006544 | S | D | P | A | V | R | H | H | V | D | R | R | A | S | E | F | K | R | V | Y | - | - | - | - | - |
| | | **Sarcotragus foetidus* meta LXNI01016685 | E | N | A | L | K | R | E | A | V | A | S | T | V | R | S | Y | E | S | G | F | - | - | - | - | - |
| | | Consensus | K | G | A | L | X | R | D | D | V | A | S | A | V | D | E | Y | E | G | R | Y | - | - | - | - | - |
| | | **Consensus #** | **101** | **102** | **103** | **104** | **105** | **106** | **107** | **108** | **109** | **110** | **111** | **112** | **113** | **114** | **115** | **116** | **117** | **118** | **119** | **120** | **121** | **122** | **123** | **124** | **125** |

| | | S.cerevisiae # | 52 | 53 | 54 | 55 | 56 | 57 | 58 | 59 | 60 | 61 | 62 | 63 | 64 | 65 | 66 | 67 | 68 | 69 | 70 | 71 | 72 | 73 | 74 | 75 | 76 |
|---|---|---|---|---|---|---|---|---|---|---|---|---|---|---|---|---|---|---|---|---|---|---|---|---|---|---|---|
| Eukaryotic | (1) | *Saccharomyces cerevisiae* | G | R | T | D | K | D | A | E | E | R | R | L | E | D | Y | N | E | A | T | H | S | Y | **Y** | N | V |
| | | *Trypanosoma brucei* | - | D | G | K | G | A | S | A | S | E | R | R | Q | D | A | T | S | L | T | N | E | Y | **Y** | D | I |
| | | *Arabidopsis thaliana* SMT1 | - | V | F | H | G | G | N | E | E | E | R | K | A | N | Y | T | D | M | V | N | K | Y | **Y** | D | L |
| | (2) | *Arabidopsis thaliana* SMT2 | - | - | - | R | R | P | K | E | I | E | T | A | E | K | V | P | D | F | V | D | T | F | **Y** | N | L |
| | | *Glycine max* SMT1 | - | V | C | Y | G | G | Q | E | E | E | R | K | A | N | Y | T | D | M | V | N | K | Y | **Y** | D | L |
| | | *Chlamydomonas reinhardtii* | G | K | N | - | A | G | E | G | I | T | D | R | S | K | T | V | H | L | V | D | V | F | **Y** | S | L |
| | | *Pneumocystis jirovecii* | D | K | K | D | G | V | H | Q | Q | N | R | F | K | G | Y | A | T | L | T | R | H | Y | **Y** | N | L |
| Bacterial | (3) | Chlamydiae sp., drinking water MAG 1 of2 | - | - | - | - | - | - | - | - | - | - | S | K | V | K | D | P | S | I | T | E | K | Y | **Y** | D | L |
| | | *Aplysina aerophoba* meta JGIcombinedJ30088_10000835 1 o | - | - | - | - | D | A | G | L | E | S | R | K | E | R | Y | Q | S | L | V | N | H | Y | **Y** | D | L |
| | | *Aplysina aerophoba* meta Ga0209021_10001462 meta | R | A | D | Q | S | R | S | P | G | E | T | A | G | D | Y | K | K | I | N | E | L | Y | **Y** | D | L |
| | | **Aplysina aerophoba* meta JGIcombinedJ30088_1000330310 | G | D | A | E | R | G | G | L | D | E | R | K | T | D | Y | L | D | F | E | K | S | Y | **Y** | N | L |
| | (2) | **Aplysina aerophoba* meta Ga0209754_100026449 | G | G | A | A | R | G | G | R | D | E | R | K | S | D | Y | K | D | F | E | N | T | Y | **Y** | D | L |
| | | *Petrosia ficiformis* meta LXNJ01000389 | H | S | A | Q | R | G | E | L | K | N | A | A | E | S | Q | T | D | I | I | S | L | Y | **Y** | D | L |
| | | **Aplysina aerophoba* meta Ga0209544_100039960 | Y | H | A | K | D | G | D | L | E | T | K | Q | N | D | F | R | K | L | A | D | L | Y | **Y** | D | L |
| | | *Nitrospira* sp., coral reef MAG 1 of2 2682229274 | - | - | - | - | D | A | G | L | E | S | R | K | E | R | Y | Q | S | L | V | N | H | Y | **Y** | D | L |
| | | *Sandaracinus* sp., marine MAG 1 of2 2775843484 | D | E | D | H | G | G | D | V | A | A | R | K | D | H | Y | F | E | M | V | K | G | Y | **Y** | D | L |
| | | Freshwater, Lake Fryxell, Antarctica meta | D | - | N | A | G | G | N | L | E | A | R | R | D | R | Y | A | D | M | V | R | G | Y | **Y** | D | L |
| | | Western Arctic Ocean meta Ga0133547_104202862 | A | Q | G | G | G | D | D | P | E | A | R | K | Q | A | Y | A | K | L | V | N | Q | Y | **Y** | D | L |
| | (1) | **Aplysina aerophoba* meta JGIcombinedJ30088_100127566 | E | F | G | R | R | N | G | R | D | R | R | A | T | D | H | R | A | F | V | E | N | Y | **Y** | D | L |
| | | Chlamydiae sp., drinking water MAG 2 of2 2619789261 | - | - | - | - | - | - | - | - | - | - | S | K | V | K | D | P | G | I | T | E | K | Y | **Y** | D | L |
| | | *Sandaracinus* sp., marine MAG 2 of2 2775843485 | - | - | - | E | D | A | D | V | E | K | R | K | A | Q | Y | A | E | L | V | N | Q | Y | **Y** | D | L |
| | | Marine sediment, Gulf of Thailand meta | - | - | - | - | D | A | G | R | E | S | R | E | Q | N | S | L | S | F | V | N | Q | Y | **Y** | D | L |
| | | Spirochaetes sp., soil MAG 2709604172 | N | E | A | E | S | G | T | I | V | S | R | T | K | N | Y | K | T | M | I | N | H | Y | **Y** | D | L |
| | | Hotspring sediment, Dewer Creek, BC meta | R | A | H | R | G | G | E | G | C | G | E | R | A | D | A | G | E | V | A | R | N | Y | **Y** | D | L |
| | | Lake sediment, Walker Lake, Nevada meta | D | E | S | S | G | G | G | L | G | A | R | R | T | Q | Y | R | A | M | V | Q | D | Y | **Y** | D | L |
| | | Marine sediment, Helgoland, North Sea meta | D | E | T | N | G | G | G | V | E | A | R | K | A | Q | Y | A | K | M | I | N | N | Y | **Y** | D | V |
| | | *Sarcotragus foetidus* meta LXNI01004480 | - | - | - | - | D | A | G | E | Q | G | R | R | E | Q | Y | R | S | L | V | N | H | Y | **Y** | D | L |
| | | *Theonella swinhoei* associated Chromatiales sp. MAG 1 of2 | - | - | - | - | D | D | D | L | E | N | R | K | E | D | Y | Q | I | L | V | N | H | Y | **Y** | D | L |
| | | *Petrosia ficiformis* meta LXNJ01003202 | - | - | - | - | G | S | D | L | E | R | R | K | A | S | Y | R | Q | L | V | S | S | Y | **Y** | D | L |
| | | *Petrosia ficiformis* meta LXNJ01014139 | - | - | - | - | D | A | G | L | E | R | R | K | E | H | Y | R | S | F | V | N | R | Y | **Y** | D | L |
| | No activity | *Aplysina aerophoba* meta JGIcombinedJ30088_10000835 2 o | D | R | A | T | P | E | A | A | G | V | N | G | Y | D | H | T | Q | T | V | K | D | Y | Y | D | L |
| | | *Petrosia ficiformis* meta LXNJ01004423 | E | K | S | K | Q | A | G | A | E | R | G | D | Y | D | H | A | E | T | V | N | D | Y | Y | D | L |
| | | *Theonella swinhoei* associated Chromatiales sp. MAG 2 of2 | G | R | I | R | Q | - | - | - | D | S | R | D | Y | D | H | T | E | T | V | N | N | Y | Y | S | L |
| | | Hotspring, Beatty, Nevada meta_Ga0114945_100210811 | D | E | T | R | G | G | G | V | E | A | R | R | E | H | Y | A | G | M | V | N | D | F | Y | D | L |
| | | *Nitrospira* sp., coral reef MAG 2 of2 2682229270 | D | R | A | T | P | E | D | A | G | V | N | R | Y | D | H | T | E | T | V | K | E | Y | Y | D | L |
| | | **Aplysina aerophoba* meta JGIcombinedJ30088_100153964 | G | G | D | V | R | G | G | L | D | E | R | R | T | D | Y | R | N | F | Q | N | T | Y | Y | D | L |
| | | **Aplysina aerophoba meta* JGIcombinedJ30088_100077196 | - | - | - | - | - | T | T | A | G | D | Q | K | D | D | P | A | E | R | S | K | D | F | Y | N | L |
| | | **Aplysina aerophoba meta* JGIcombinedJ30088_100209763 | G | S | D | E | V | A | S | L | E | Q | R | K | T | A | Y | E | D | L | T | N | K | Y | Y | D | L |
| | | **Aplysina aerophoba meta* JGIcombinedJ30088_100057164 | G | G | P | A | R | G | G | L | D | L | R | K | S | E | Y | R | N | F | A | K | L | Y | F | D | L |
| | | **Sarcotragus foetidus* meta LXNI01006544 | I | P | S | Q | R | E | D | L | D | R | R | V | E | E | Y | R | R | F | A | N | T | Y | Y | D | L |
| | | **Sarcotragus foetidus* meta LXNI01016685 | G | H | D | E | A | G | S | P | D | R | R | K | S | E | Y | E | D | L | T | N | K | Y | Y | D | L |
| | | Consensus | - | - | - | - | G | G | G | L | E | E | R | K | E | D | Y | R | D | L | V | N | X | Y | **Y** | D | L |
| | | **Consensus #** | **126** | **127** | **128** | **129** | **130** | **131** | **132** | **133** | **134** | **135** | **136** | **137** | **138** | **139** | **140** | **141** | **142** | **143** | **144** | **145** | **146** | **147** | **148** | **149** | **150** |

Region I (S.cerevisiae # 81–91)

| Group | | Sequence | 77 | 78 | 79 | 80 | 81 | 82 | 83 | 84 | 85 | 86 | 87 | 88 | 89 | 90 | 91 | 92 | 93 | 94 | 95 | 96 | 97 | 98 | 99 | 100 | 101 |
|---|---|---|---|---|---|---|---|---|---|---|---|---|---|---|---|---|---|---|---|---|---|---|---|---|---|---|---|
| Eukaryotic | (1) | Saccharomyces cerevisiae | V | T | D | F | Y | E | Y | G | W | G | S | S | F | H | F | S | R | F | Y | K | G | E | S | F | A |
| | | Trypanosoma brucei | V | T | D | F | Y | E | Y | G | W | G | Q | N | F | H | F | A | P | R | Y | M | N | E | T | F | Y |
| | | Arabidopsis thaliana SMT1 | A | T | S | F | Y | E | Y | G | W | G | E | S | F | H | F | A | Q | R | W | K | G | E | S | L | R |
| | (2) | Arabidopsis thaliana SMT2 | V | T | D | I | Y | E | W | G | W | G | Q | S | F | H | F | S | P | S | I | P | G | K | S | H | K |
| | | Glycine max SMT1 | V | T | S | F | Y | E | F | G | W | G | E | S | F | H | F | A | P | R | W | K | G | E | S | L | R |
| | | Chlamydomonas reinhardtii | V | T | D | I | Y | E | W | G | W | G | Q | S | F | H | F | S | P | K | L | P | N | K | D | L | K |
| | | Pneumocystis jirovecii | V | T | D | L | Y | E | Y | G | W | S | P | S | F | H | F | C | R | F | S | R | G | E | S | F | S |
| Bacterial | (3) | Chlamydiae sp., drinking water MAG 1 of 2 | V | T | D | F | Y | E | F | G | W | G | K | S | F | H | F | A | P | Q | T | R | T | E | D | T | R |
| | | Aplysina aerophoba meta JGIcombinedJ13088_10000835 1 of | V | T | D | F | F | E | F | G | W | G | Q | S | F | H | F | A | P | R | Q | R | G | E | S | F | K |
| | | Aplysina aerophoba meta Ga0209021_10001462 meta | V | T | D | F | F | E | Y | G | W | G | K | S | F | H | F | A | P | R | V | P | G | E | S | F | R |
| | | *Aplysina aerophoba meta JGIcombinedJ13088_100033010 | V | T | D | F | F | E | Y | G | W | G | R | S | F | H | F | A | P | R | Y | E | G | E | S | F | A |
| | (2) | *Aplysina aerophoba meta Ga0209754_10002649 | V | T | D | F | Y | E | Y | G | W | G | R | S | F | H | F | A | P | R | I | E | G | E | S | F | A |
| | | Petrosia ficiformis meta LXNI01000389 | V | T | D | F | Y | E | Y | G | W | G | K | S | F | H | F | A | P | R | T | P | G | E | T | F | K |
| | | *Aplysina aerophoba meta Ga0209544_10003960 | V | T | D | F | Y | E | F | G | W | G | R | S | F | H | F | A | P | R | V | P | G | E | T | F | K |
| | | Nitrospira sp., coral reef MAG 1 of 2 268222974 | V | T | D | F | Y | E | F | G | W | G | Q | S | F | H | F | A | P | R | R | R | G | E | S | F | K |
| | | Sandaracinus sp., marine MAG 1 of 2 2775843484 | A | T | D | F | F | E | Y | G | W | G | D | S | F | H | F | A | P | R | Y | R | G | E | T | F | R |
| | | Freshwater, Lake Fryxell, Antarctica meta | V | T | D | I | Y | E | R | G | W | G | D | S | F | H | F | A | P | R | Y | R | G | E | A | F | E |
| | | Western Arctic Ocean meta Ga0133547_10420862 | V | T | D | F | Y | E | Y | G | W | G | R | S | F | H | F | A | P | R | R | K | G | E | S | F | E |
| | (1) | *Aplysina aerophoba meta JGIcombinedJ13088_100127566 | A | T | D | L | Y | E | Y | G | W | G | E | S | F | H | F | A | P | R | V | P | G | E | S | R | A |
| | | Chlamydiae sp., drinking water MAG 2 of 2 2619789261 | V | T | D | F | F | E | Y | G | W | G | K | S | F | H | F | A | P | Q | T | K | T | E | S | T | R |
| | | Sandaracinus sp., marine MAG 2 of 2 2775843485 | A | T | D | F | Y | T | F | G | W | G | E | H | F | H | F | A | P | R | Y | A | H | E | G | F | L |
| | | Marine sediment, Gulf of Thailand meta | V | T | D | F | Y | E | F | G | W | G | T | S | F | H | F | A | P | R | T | R | R | E | T | F | E |
| | | Spirochaetes sp., soil MAG 2709604172 | V | T | D | F | Y | E | Y | G | W | G | Q | S | F | H | F | A | P | R | K | R | D | E | S | F | K |
| | | Hotspring sediment, Dewer Creek, BC meta | A | T | D | A | Y | E | W | G | W | G | E | S | F | H | F | G | V | R | R | R | G | E | S | L | E |
| | | Lake sediment, Walker Lake, Nevada meta | V | T | E | F | Y | E | R | G | W | G | E | S | F | H | F | A | P | R | H | R | R | E | S | F | E |
| | | Marine sediment, Helgoland, North Sea meta | V | T | D | F | Y | E | H | G | W | G | Q | S | F | H | F | A | A | R | S | K | M | E | S | F | D |
| | | Sarcotragus foetidus meta LXNI01004480 | V | T | D | F | Y | E | F | G | W | G | Q | S | F | H | F | A | P | R | R | R | G | E | S | F | K |
| | | Theonella swinhoei associated Chromatiales sp. MAG 1 of 2 | A | T | D | F | Y | E | F | G | W | G | Q | S | F | H | F | A | P | R | L | R | G | E | S | F | K |
| | | Petrosia ficiformis meta LXNI01003202 | V | T | D | F | Y | E | W | G | W | G | K | N | F | H | F | A | P | R | R | R | G | E | S | F | K |
| | | Petrosia ficiformis meta LXNI01014139 | A | T | D | F | Y | E | F | G | W | G | Q | S | F | H | F | A | P | R | R | R | G | E | S | F | K |
| | No activity | Aplysina aerophoba meta JGIcombinedJ13088_10000835 2 of | C | S | E | F | M | V | Y | G | W | G | E | S | L | H | F | A | P | L | T | P | Q | E | S | L | G |
| | | Petrosia ficiformis meta LXNI01004423 | C | S | E | F | M | Q | F | G | W | N | E | S | L | H | F | A | P | L | T | P | G | E | T | L | E |
| | | Theonella swinhoei associated Chromatiales sp. MAG 2 of 2 | C | S | E | F | M | T | W | G | W | N | D | S | L | H | F | A | P | L | A | P | G | E | S | L | Q |
| | | Hotspring, Beatty, Nevada meta_Ga0114945_10021081 1 | V | T | D | F | Y | E | F | G | W | G | Q | A | F | H | F | A | P | R | H | R | G | E | S | L | A |
| | | Nitrospira sp., coral reef MAG 2 of 2 268222970 | C | S | P | F | M | V | Y | G | W | G | E | S | L | H | F | A | P | L | T | P | Q | E | S | L | E |
| | | *Aplysina aerophoba meta JGIcombinedJ13088_100153964 | V | T | D | F | Y | E | Y | G | W | G | Q | S | F | H | F | A | P | R | V | V | G | E | S | F | A |
| | | *Aplysina aerophoba meta JGIcombinedJ13088_100077196 | I | T | D | F | T | Q | Y | G | W | S | T | S | I | H | F | A | P | R | N | P | G | E | S | V | A |
| | | *Aplysina aerophoba meta JGIcombinedJ13088_100209763 | V | T | D | L | Y | E | Y | G | W | G | K | S | F | H | F | A | P | R | A | T | N | E | S | F | A |
| | | *Aplysina aerophoba meta JGIcombinedJ13088_100057164 | I | T | D | F | Y | E | Y | G | W | S | R | S | F | H | F | A | P | R | T | R | N | E | S | F | R |
| | | *Sarcotragus foetidus meta LXNI01006644 | V | T | D | F | Y | E | F | G | W | G | R | S | F | H | F | A | P | Q | A | P | G | E | S | F | A |
| | | *Sarcotragus foetidus meta LXNI01016685 | V | T | G | F | F | E | Y | G | W | G | K | S | F | H | F | A | P | R | A | P | N | E | S | F | A |
| | | Consensus | V | T | D | F | Y | E | Y | G | W | G | Q | S | F | H | F | A | P | R | T | R | G | E | S | F | K |
| | | Consensus # | 151 | 152 | 153 | 154 | 155 | 156 | 157 | 158 | 159 | 160 | 161 | 162 | 163 | 164 | 165 | 166 | 167 | 168 | 169 | 170 | 171 | 172 | 173 | 174 | 175 |

| | | *S.cerevisiae* # | 102 | 103 | 104 | 105 | 106 | 107 | 108 | 109 | 110 | 111 | 112 | 113 | 114 | 115 | 116 | 117 | 118 | 119 | 120 | 121 | 122 |
|---|---|---|---|---|---|---|---|---|---|---|---|---|---|---|---|---|---|---|---|---|---|---|---|
| Eukaryotic | (1) | *Saccharomyces cerevisiae* | A | S | I | A | R | H | E | H | Y | L | A | Y | K | A | G | I | Q | R | G | D | L |
| | | *Trypanosoma brucei* | E | S | L | A | R | Y | E | Y | F | L | A | Y | H | A | Q | F | K | P | T | D | T |
| | | *Arabidopsis thaliana* SMT1 | E | S | I | K | R | H | E | H | F | L | A | L | Q | L | G | I | Q | P | G | Q | K |
| | (2) | *Arabidopsis thaliana* SMT2 | D | A | T | R | L | H | E | E | M | A | V | D | L | I | Q | V | K | P | G | Q | K |
| | | *Glycine max* SMT1 | E | S | I | K | R | H | E | H | F | L | A | S | Q | L | G | L | K | P | G | Q | K |
| | | *Chlamydomonas reinhardtii* | A | S | E | A | A | H | E | A | R | I | A | A | L | L | R | L | Q | P | G | Q | K |
| | | *Pneumocystis jirovecii* | K | A | I | A | R | H | E | H | Y | L | A | S | H | A | G | I | R | E | G | E | T |
| Bacterial | (3) | Chlamydiae sp., drinking water MAG 1 of2 | E | A | L | V | R | H | E | L | K | L | A | D | E | L | K | L | K | P | G | M | K |
| | | *Aplysina aerophoba* meta JGIcombinedJ30088_10000835 1 of | A | S | L | L | K | H | E | Q | F | L | A | D | R | L | S | L | K | P | G | M | Q |
| | | *Aplysina aerophoba* meta Ga0209021_10001462 meta | A | S | L | A | R | H | E | R | N | L | S | Q | A | L | E | L | R | P | G | M | V |
| | | **Aplysina aerophoba* meta JGIcombinedJ30088_1000330310 | A | S | L | A | R | H | E | H | Y | M | A | H | R | L | N | L | Q | P | G | M | V |
| | (2) | **Aplysina aerophoba* meta Ga0209754_100026449 | A | S | L | A | R | H | E | H | Y | L | A | H | R | L | D | L | Q | P | G | M | T |
| | | *Petrosia ficiformis* meta LXNJ01000389 | A | S | L | V | R | H | E | H | R | L | A | D | A | L | E | L | K | E | G | M | T |
| | | **Aplysina aerophoba* meta Ga0209544_100039960 | A | S | L | A | R | H | E | H | F | L | A | L | K | L | G | L | K | P | G | M | L |
| | | *Nitrospira* sp., coral reef MAG 1 of2 2682229274 | A | S | L | L | R | H | E | H | Y | L | A | D | R | L | S | L | K | P | G | M | Q |
| | | *Sandaracinus* sp., marine MAG 1 of2 2775843484 | E | S | I | R | R | H | Q | H | F | L | A | A | R | L | G | L | K | P | G | M | R |
| | | Freshwater, Lake Fryxell, Antarctica meta | A | S | L | K | R | H | Q | H | Y | I | A | L | R | L | G | L | T | T | G | M | T |
| | | Western Arctic Ocean meta Ga0133547_104202862 | A | S | L | L | R | H | E | H | Y | L | A | L | S | I | G | V | G | R | G | Q | R |
| | (1) | **Aplysina aerophoba* meta JGIcombinedJ30088_100127566 | A | S | I | A | R | Q | E | H | F | V | A | L | K | L | G | L | R | P | G | M | R |
| | | Chlamydiae sp., drinking water MAG 2 of2 2619789261 | E | A | M | L | R | Y | E | L | K | M | A | E | A | L | K | L | K | P | G | M | K |
| | | *Sandaracinus* sp., marine MAG 2 of2 2775843485 | E | S | L | K | R | H | E | R | F | L | A | R | K | L | R | L | Y | P | G | M | E |
| | | Marine sediment, Gulf of Thailand meta | E | S | L | A | R | H | E | R | R | L | A | N | A | L | R | L | N | S | K | S | V |
| | | Spirochaetes sp., soil MAG 2709604172 | D | S | L | I | R | H | E | H | Y | L | A | E | Q | L | E | L | D | P | G | S | E |
| | | Hotspring sediment, Dewer Creek, BC meta | A | S | L | V | R | H | E | L | R | L | A | E | R | M | G | L | R | A | G | M | T |
| | | Lake sediment, Walker Lake, Nevada meta | A | S | I | A | R | H | Q | H | F | L | A | S | R | L | R | L | A | A | G | M | R |
| | | Marine sediment, Helgoland, North Sea meta | A | S | L | A | R | A | E | H | Y | L | A | L | R | L | G | L | Q | A | G | M | R |
| | | *Sarcotragus foetidus* meta LXNI01004480 | A | S | L | L | R | H | Q | H | F | L | A | D | R | L | A | L | K | P | G | T | Q |
| | | *Theonella swinhoei* associated Chromatiales sp. MAG 1 of2 | E | S | L | L | R | H | E | H | F | L | A | D | Q | L | S | L | K | P | G | M | Q |
| | | *Petrosia ficiformis* meta LXNJ01003202 | A | S | Q | L | R | H | Q | Y | F | L | A | D | R | L | S | L | K | P | G | M | R |
| | | *Petrosia ficiformis* meta LXNJ01014139 | A | S | L | L | R | H | Q | H | F | L | A | E | R | L | S | L | K | P | G | M | N |
| | No activity | *Aplysina aerophoba* meta JGIcombinedJ30088_10000835 2 of | D | S | K | V | R | H | Q | R | L | M | I | A | K | L | E | L | H | P | G | M | T |
| | | *Petrosia ficiformis* meta LXNJ01004423 | E | S | I | V | R | H | Q | R | L | M | I | D | K | L | E | L | R | E | G | M | R |
| | | *Theonella swinhoei* associated Chromatiales sp. MAG 2 of2 | D | S | I | V | R | H | Q | R | H | M | I | K | S | L | Q | L | Q | E | G | M | R |
| | | Hotspring, Beatty, Nevada meta_Ga0114945_100210811 | A | S | L | A | R | H | E | F | F | L | A | L | R | L | G | L | R | P | G | M | R |
| | | *Nitrospira* sp., coral reef MAG 2 of2 2682229270 | D | S | K | V | R | H | Q | R | L | M | I | A | K | L | E | L | H | P | G | M | T |
| | | **Aplysina aerophoba* meta JGIcombinedJ30088_100153964 | A | S | L | A | R | H | E | H | Y | I | A | H | V | L | S | L | R | P | G | M | V |
| | | **Aplysina aerophoba meta* JGIcombinedJ30088_100077196 | A | S | L | A | R | H | E | H | F | I | A | M | R | L | A | L | E | P | G | M | H |
| | | **Aplysina aerophoba meta* JGIcombinedJ30088_100209763 | D | S | L | A | R | H | E | Q | Y | L | A | Q | R | L | D | L | R | P | G | M | R |
| | | **Aplysina aerophoba meta* JGIcombinedJ30088_100057164 | A | S | L | T | R | H | E | H | Y | L | A | Q | E | L | Q | L | R | P | G | M | V |
| | | **Sarcotragus foetidus* meta LXNI01006544 | A | S | L | A | R | H | E | H | R | V | A | H | V | L | G | L | R | A | G | M | R |
| | | **Sarcotragus foetidus* meta LXNI01016685 | Q | S | I | A | R | H | E | R | N | M | A | E | R | L | A | L | S | P | G | M | R |
| | | Consensus | A | S | L | A | R | H | E | H | F | L | A | D | R | L | G | L | K | P | G | M | R |
| | | **Consensus #** | 176 | 177 | 178 | 179 | 180 | 181 | 182 | 183 | 184 | 185 | 186 | 187 | 188 | 189 | 190 | 191 | 192 | 193 | 194 | 195 | 196 |

| | | | | Region II | | | | | | | | | | Methyltransferase domain | | | | | | | | | | | | | | | | | |
|---|---|---|---|---|---|---|---|---|---|---|---|---|---|---|---|---|---|---|---|---|---|---|---|---|---|---|---|---|---|---|---|
| | | *S. cerevisiae* # | 123 | 124 | 125 | 126 | 127 | 128 | 129 | 130 | 131 | 132 | 133 | 134 | 135 | 136 | 137 | 138 | 139 | 140 | 141 | 142 | 143 | 144 | 145 | 146 | 147 | 148 | 149 | 150 | 151 |
| Eukaryotic | (1) | *Saccharomyces cerevisiae* | V | L | D | V | G | C | G | V | G | G | P | A | R | E | I | A | R | F | T | G | C | N | V | I | G | L | N | N | N |
| | | *Trypanosoma brucei* | V | L | D | V | G | C | G | I | G | G | P | A | R | N | M | V | R | F | T | S | C | N | V | M | G | V | N | N | N |
| | | *Arabidopsis thaliana* SMT1 | V | L | D | V | G | C | G | I | G | G | P | L | R | E | I | A | R | F | S | N | S | V | V | T | G | L | N | N | N |
| | (2) | *Arabidopsis thaliana* SMT2 | I | L | D | V | G | C | G | V | G | G | P | M | R | A | I | A | S | H | S | R | A | N | V | V | G | I | T | I | N |
| | | *Glycine max* SMT1 | V | L | D | V | G | C | G | I | G | G | P | L | R | E | I | S | R | F | S | L | T | S | I | T | G | L | N | N | N |
| | | *Chlamydomonas reinhardtii* | A | L | D | C | G | C | G | V | G | G | P | M | R | T | V | A | A | V | S | G | A | H | I | T | G | I | T | I | N |
| | | *Pneumocystis jirovecii* | V | L | D | I | G | S | G | V | G | G | P | A | L | E | I | S | V | F | T | G | A | N | I | V | G | I | N | N | N |
| Bacterial | (3) | Chlamydiae sp., drinking water MAG 1 of 2 | V | L | D | V | G | C | G | I | A | G | P | M | K | N | I | A | E | K | S | G | A | N | I | V | G | L | N | I | N |
| | | *Aplysina aerophoba* meta JGIcombinedJ30088_10000835 1 of | V | L | D | V | G | C | G | V | G | G | P | M | G | A | L | A | R | Y | S | G | A | S | F | V | G | I | N | N | N |
| | | *Aplysina aerophoba* meta Ga0209021_10001462 meta | A | A | D | F | G | C | G | I | G | G | P | L | L | E | I | A | R | F | S | G | A | K | I | V | G | I | N | I | N |
| | | **Aplysina aerophoba* meta JGIcombinedJ30088_1000330310 | V | L | D | L | G | C | G | I | G | G | P | L | R | E | I | V | R | F | S | G | A | K | I | V | G | V | N | I | N |
| | (2) | **Aplysina aerophoba* meta Ga0209754_100026449 | A | V | D | L | G | C | G | I | G | G | S | L | R | E | I | A | R | F | S | G | A | R | I | L | G | V | N | N | N |
| | | *Petrosia ficiformis* meta LXNJ01000389 | V | A | D | L | G | C | G | I | G | G | S | M | L | E | I | A | R | Y | S | G | A | K | I | V | G | V | N | N | N |
| | | **Aplysina aerophoba* meta Ga0209544_100039960 | A | A | D | I | G | C | G | I | G | G | P | L | I | E | I | A | R | F | S | G | A | N | I | V | G | V | N | T | N |
| | | *Nitrospira* sp., coral reef MAG 1 of 2 2682229274 | V | L | D | V | G | C | G | V | G | G | P | M | G | A | L | A | R | Y | S | G | A | S | F | V | G | I | N | N | N |
| | | *Sandaracinus* sp., marine MAG 1 of 2 2775843484 | V | L | D | V | G | C | G | I | G | G | P | M | R | S | V | A | R | F | S | G | A | T | V | V | G | I | N | N | N |
| | | Freshwater, Lake Fryxell, Antarctica meta | A | L | D | I | G | C | G | V | G | G | P | M | R | E | I | A | R | I | S | G | T | R | V | Y | G | V | N | N | N |
| | | Western Arctic Ocean meta Ga0133547_104202862 | V | L | D | V | G | C | G | V | G | G | P | M | T | N | I | A | H | F | A | A | C | D | V | V | G | I | N | N | N |
| | (1) | **Aplysina aerophoba* meta JGIcombinedJ30088_100127566 | V | A | D | F | G | C | G | V | G | G | P | L | R | E | I | A | R | F | S | G | S | Q | I | V | G | V | N | I | S |
| | | Chlamydiae sp., drinking water MAG 2 of 2 2619789261 | V | L | D | I | G | C | G | I | A | G | P | M | K | N | I | A | E | K | S | G | A | T | I | V | G | L | N | I | N |
| | | *Sandaracinus* sp., marine MAG 2 of 2 2775843485 | C | L | D | V | G | C | G | I | G | G | P | M | R | H | I | A | K | H | T | G | A | R | V | V | G | V | N | N | N |
| | | Marine sediment, Gulf of Thailand meta | V | L | D | V | G | C | G | V | G | G | P | M | R | E | I | A | S | S | S | G | A | R | V | V | G | V | N | N | N |
| | | Spirochaetes sp., soil MAG 2709604172 | V | L | D | L | G | C | G | V | G | G | P | L | R | E | L | A | R | Y | S | G | A | R | I | T | G | V | N | N | N |
| | | Hotspring sediment, Dewer Creek, BC meta | A | L | D | L | G | C | G | I | G | G | P | M | R | N | I | A | R | A | T | G | A | R | I | V | G | V | N | I | C |
| | | Lake sediment, Walker Lake, Nevada meta | V | L | D | V | G | C | G | V | G | G | P | M | R | S | I | A | R | F | T | G | A | H | V | V | G | V | N | I | S |
| | | Marine sediment, Helgoland, North Sea meta | V | L | D | V | G | C | G | V | G | G | P | M | R | S | I | A | R | F | S | D | A | T | V | E | G | V | N | N | N |
| | | *Sarcotragus foetidus* meta LXNI01004480 | V | L | D | V | G | C | G | V | G | G | P | M | G | N | L | A | R | Y | S | G | A | G | F | V | G | I | N | N | N |
| | | *Theonella swinhoei* associated Chromatiales sp. MAG 1 of 2 | V | I | D | L | G | C | G | I | G | G | P | M | R | T | L | A | R | Y | S | G | A | S | F | V | G | L | N | N | N |
| | | *Petrosia ficiformis* meta LXNJ01003202 | I | L | D | A | G | C | G | V | G | G | P | M | S | N | L | A | R | H | S | G | A | S | F | V | G | V | N | I | S |
| | | *Petrosia ficiformis* meta LXNJ01014139 | V | L | D | V | G | C | G | V | G | G | P | M | G | N | L | A | R | H | S | G | A | S | F | V | G | I | N | N | N |
| | No activity | *Aplysina aerophoba* meta JGIcombinedJ30088_10000835 2 of | V | V | D | V | G | C | G | I | G | G | P | M | R | R | V | V | R | E | A | G | V | R | V | V | G | I | N | I | N |
| | | *Petrosia ficiformis* meta LXNJ01004423 | I | V | D | V | G | C | G | V | G | G | P | M | R | R | V | A | R | E | S | G | S | R | V | L | C | I | N | N | N |
| | | *Theonella swinhoei* associated Chromatiales sp. MAG 2 of 2 | V | I | D | V | G | C | G | I | G | E | P | M | R | R | V | A | R | E | S | G | A | R | V | L | C | L | N | N | N |
| | | Hotspring, Beatty, Nevada meta_Ga0114945_100210811 | V | L | D | V | G | C | G | V | G | G | P | M | R | A | I | A | R | L | S | G | A | A | I | V | G | V | N | N | N |
| | | *Nitrospira* sp., coral reef MAG 2 of 2 2682229270 | I | V | D | V | G | C | G | V | G | G | P | M | R | R | V | V | R | E | A | G | V | R | V | V | G | I | N | I | N |
| | | **Aplysina aerophoba* meta JGIcombinedJ30088_100153964 | A | V | D | L | G | C | G | V | G | G | P | A | R | E | I | A | R | F | S | G | A | T | I | L | G | V | N | N | N |
| | | **Aplysina aerophoba meta* JGIcombinedJ30088_100077196 | V | A | D | L | G | C | G | I | G | G | P | L | R | E | I | V | R | L | S | G | A | M | V | T | G | V | N | N | N |
| | | **Aplysina aerophoba meta* JGIcombinedJ30088_100209763 | V | A | D | L | G | C | G | I | G | G | P | L | R | E | I | A | R | F | S | G | A | T | I | A | G | V | N | N | N |
| | | **Aplysina aerophoba meta* JGIcombinedJ30088_100057164 | A | L | D | L | G | C | G | I | G | G | P | L | R | E | M | I | R | F | S | G | A | R | I | V | G | V | S | I | S |
| | | **Sarcotragus foetidus* meta LXNI01006544 | V | I | D | L | G | C | G | I | G | G | P | A | R | Q | I | A | R | F | S | G | A | S | I | V | G | L | N | N | N |
| | | **Sarcotragus foetidus* meta LXNI01016685 | V | A | D | L | G | C | G | V | G | G | P | L | R | E | I | A | R | F | S | G | A | T | I | V | G | V | N | N | N |
| | | Consensus | V | L | D | V | G | C | G | V | G | G | P | M | R | E | I | A | R | F | S | G | A | R | I | V | G | V | N | N | N |
| | | Consensus # | 197 | 198 | 199 | 200 | 201 | 202 | 203 | 204 | 205 | 206 | 207 | 208 | 209 | 210 | 211 | 212 | 213 | 214 | 215 | 216 | 217 | 218 | 219 | 220 | 221 | 222 | 223 | 224 | 225 |

**Methyltransferase domain**

| | | *S. cerevisiae* # | 152 | 153 | 154 | 155 | 156 | 157 | 158 | 159 | 160 | 161 | 162 | 163 | 164 | 165 | 166 | 167 | 168 | 169 | 170 | 171 | 172 | 173 | 174 | 175 | 176 |
|---|---|---|---|---|---|---|---|---|---|---|---|---|---|---|---|---|---|---|---|---|---|---|---|---|---|---|---|
| Eukaryotic | (1) | *Saccharomyces cerevisiae* | D | Y | Q | I | A | K | A | K | Y | Y | A | K | K | Y | N | L | S | D | Q | M | D | F | V | K | G |
| | | *Trypanosoma brucei* | E | Y | Q | I | N | R | A | R | Q | H | D | S | R | Y | G | M | S | G | K | I | N | Y | T | K | T |
| | | *Arabidopsis thaliana* SMT1 | E | Y | Q | I | T | R | G | K | E | L | N | R | L | A | G | V | D | K | T | C | N | F | V | K | A |
| | (2) | *Arabidopsis thaliana* SMT2 | E | Y | Q | V | N | R | A | R | L | H | N | K | K | A | G | L | D | A | L | C | E | V | V | C | G |
| | | *Glycine max* SMT1 | E | Y | Q | I | T | R | G | K | E | L | N | R | I | A | G | V | D | K | T | C | N | F | V | K | A |
| | | *Chlamydomonas reinhardtii* | Q | Y | Q | V | D | R | A | K | T | H | N | A | R | Q | G | L | A | P | L | T | D | V | V | R | G |
| | | *Pneumocystis jirovecii* | D | Y | Q | I | E | R | A | Q | F | Y | A | Q | K | K | G | L | S | D | K | L | S | F | V | K | G |
| Bacterial | (3) | Chlamydiae sp., drinking water MAG 1 of 2 | A | Y | Q | I | K | K | A | K | Q | Y | I | K | R | C | G | L | E | N | Q | C | S | F | H | Q | G |
| | | *Aplysina aerophoba* meta JGIcombinedJ30088_10000835 1 of | A | Y | Q | I | K | R | A | K | L | H | T | R | D | - | - | V | A | S | L | C | R | F | I | C | G |
| | | *Aplysina aerophoba* meta Ga0209021_10001462 meta | G | Y | Q | L | E | R | A | R | Q | L | A | G | E | A | D | L | T | H | L | A | E | F | L | Q | C |
| | | **Aplysina aerophoba* meta JGIcombinedJ30088_1000330310 | E | Y | Q | L | K | R | A | A | E | L | T | E | A | S | G | L | S | H | L | A | E | Y | R | N | C |
| | (2) | **Aplysina aerophoba* meta Ga0209754_100026449 | E | Y | Q | L | K | R | A | A | K | L | T | E | A | E | Y | L | S | H | R | V | E | Y | L | K | C |
| | | *Petrosia ficiformis* meta LXNJ01000389 | S | Y | Q | L | D | R | A | R | S | Y | A | K | Q | L | D | L | S | H | L | T | E | F | I | D | C |
| | | **Aplysina aerophoba* meta Ga0209544_100039960 | A | Y | Q | I | K | R | A | Q | T | L | T | Q | E | A | G | L | S | D | L | A | E | F | L | H | C |
| | | *Nitrospira* sp., coral reef MAG 1 of 2 2682229274 | A | Y | Q | I | E | R | A | K | V | H | T | R | D | - | - | V | R | S | L | C | R | F | I | H | G |
| | | *Sandaracinus* sp., marine MAG 1 of 2 2775843484 | P | Y | Q | L | E | K | A | A | G | Y | N | R | E | A | G | L | A | A | Q | T | E | L | V | E | A |
| | | Freshwater, Lake Fryxell, Antarctica meta | S | Y | Q | V | S | K | A | R | T | Y | N | Q | E | A | R | L | T | A | L | C | E | L | L | E | A |
| | | Western Arctic Ocean meta Ga0133547_104202862 | A | Y | Q | I | E | H | G | E | T | Y | V | A | E | A | G | L | T | G | R | C | S | F | E | K | A |
| | (1) | **Aplysina aerophoba* meta JGIcombinedJ30088_100127566 | A | Y | Q | I | R | R | A | V | A | Y | T | E | A | A | G | L | S | Q | L | G | T | Y | L | E | S |
| | | Chlamydiae sp., drinking water MAG 2 of 2 2619789261 | S | Y | Q | I | K | K | A | T | Q | Y | I | Q | N | S | G | L | E | K | Q | C | S | F | H | H | G |
| | | *Sandaracinus* sp., marine MAG 2 of 2 2775843485 | A | Y | Q | I | E | R | G | Q | K | I | T | R | R | R | K | L | D | H | L | A | S | F | M | K | A |
| | | Marine sediment, Gulf of Thailand meta | E | Y | Q | I | E | R | G | E | R | H | N | E | E | A | G | L | T | N | L | C | S | F | R | R | A |
| | | Spirochaetes sp., soil MAG 2709604172 | A | Y | Q | I | A | K | C | G | K | Y | I | E | E | E | G | L | T | N | Q | C | S | T | L | K | A |
| | | Hotspring sediment, Dewer Creek, BC meta | G | Y | Q | V | R | R | A | V | E | L | N | R | A | A | G | L | D | G | Q | L | G | V | I | E | C |
| | | Lake sediment, Walker Lake, Nevada meta | A | M | Q | V | Q | K | A | V | A | Y | A | A | R | D | G | L | A | D | R | C | E | L | L | E | A |
| | | Marine sediment, Helgoland, North Sea meta | D | Y | Q | L | E | K | V | R | S | H | N | E | E | A | G | L | G | H | L | C | S | A | F | K | G |
| | | *Sarcotragus foetidus* meta LXNI01004480 | A | Y | Q | I | E | R | A | K | H | H | T | R | D | - | - | V | G | S | L | C | R | F | I | H | G |
| | | *Theonella swinhoei* associated Chromatiales sp. MAG 1 of 2 | A | Y | Q | I | K | R | A | E | A | Q | S | Q | D | - | - | I | Q | S | L | C | R | F | I | H | G |
| | | *Petrosia ficiformis* meta LXNJ01003202 | D | Y | Q | I | E | R | A | K | F | H | T | R | D | - | - | V | K | A | L | C | R | F | I | H | G |
| | | *Petrosia ficiformis* meta LXNJ01014139 | A | Y | Q | I | E | R | A | K | L | H | T | R | D | - | - | V | Q | S | L | C | R | F | I | H | G |
| | No activity | *Aplysina aerophoba* meta JGIcombinedJ30088_10000835 2 of | H | I | Q | L | A | N | A | K | R | L | N | A | E | A | G | L | D | H | M | V | D | Y | E | T | C |
| | | *Petrosia ficiformis* meta LXNJ01004423 | E | Q | Q | L | E | K | A | R | R | R | N | V | E | A | G | L | D | H | M | A | E | Y | M | K | C |
| | | *Theonella swinhoei* associated Chromatiales sp. MAG 2 of 2 | E | Q | Q | L | E | K | A | K | E | K | S | L | E | A | G | L | D | H | L | A | E | Y | I | K | C |
| | | Hotspring, Beatty, Nevada meta_Ga0114945_100210811 | D | Y | Q | I | R | R | G | A | E | Q | T | R | A | A | G | L | E | R | L | C | A | L | V | K | R |
| | | *Nitrospira* sp., coral reef MAG 2 of 2 2682229270 | K | I | Q | L | E | R | A | K | R | L | N | S | E | A | G | L | D | H | M | V | D | Y | E | A | C |
| | | **Aplysina aerophoba* meta JGIcombinedJ30088_100153964 | E | Y | Q | L | K | R | A | A | R | Q | T | D | A | E | N | L | S | H | L | V | D | Y | L | K | C |
| | | **Aplysina aerophoba meta* JGIcombinedJ30088_100077196 | S | Y | Q | V | D | R | A | R | E | L | T | Q | E | A | G | L | T | D | Y | A | E | F | M | E | C |
| | | **Aplysina aerophoba meta* JGIcombinedJ30088_100209763 | E | Y | Q | L | D | R | A | R | T | L | T | D | E | A | G | L | G | H | L | A | E | F | L | K | C |
| | | **Aplysina aerophoba meta* JGIcombinedJ30088_100057164 | G | Y | G | I | D | R | A | R | K | Q | T | D | E | A | G | L | G | H | M | A | E | F | V | E | C |
| | | **Sarcotragus foetidus* meta LXNI01006544 | P | S | Q | V | E | R | A | R | R | I | T | E | K | E | G | L | D | H | L | V | S | Y | R | H | G |
| | | **Sarcotragus foetidus* meta LXNI01016685 | E | Y | Q | V | E | R | A | R | K | L | T | N | E | A | G | L | G | H | L | A | E | F | L | K | C |
| | | Consensus | X | Y | Q | I | E | R | A | K | E | L | T | R | E | A | G | L | D | H | L | C | E | F | L | K | C |
| | | **Consensus #** | **226** | **227** | **228** | **229** | **230** | **231** | **232** | **233** | **234** | **235** | **236** | **237** | **238** | **239** | **240** | **241** | **242** | **243** | **244** | **245** | **246** | **247** | **248** | **249** | **250** |

| | | | Methyltransferase domain | | | | | | | | | | | | | | | | Region III | | | | | | | | |
|---|---|---|---|---|---|---|---|---|---|---|---|---|---|---|---|---|---|---|---|---|---|---|---|---|---|---|---|
| | | *S.cerevisiae* # | 177 | 178 | 179 | 180 | 181 | 182 | | 183 | 184 | 185 | 186 | 187 | 188 | 189 | 190 | 191 | 192 | 193 | 194 | 195 | 196 | 197 | 198 | | 199 |
| Eukaryotic | (1) | *Saccharomyces cerevisiae* | D | F | M | K | M | D | - | F | E | E | N | T | F | D | K | V | Y | A | I | E | A | T | C | - | H |
| | | *Trypanosoma brucei* | D | F | C | N | M | C | - | F | G | D | N | E | F | D | G | A | Y | A | I | E | A | T | C | - | H |
| | | *Arabidopsis thaliana* SMT1 | D | F | M | K | M | P | - | F | P | E | N | S | F | D | A | V | Y | A | I | E | A | T | C | - | H |
| | (2) | *Arabidopsis thaliana* SMT2 | N | F | L | Q | M | P | - | F | D | D | N | S | F | D | G | A | Y | S | I | E | A | T | C | - | H |
| | | *Glycine max* SMT1 | D | F | M | K | M | P | - | L | P | D | N | S | F | D | A | V | Y | A | I | E | A | T | C | - | H |
| | | *Chlamydomonas reinhardtii* | D | F | T | N | M | P | - | F | K | E | N | T | F | D | G | A | Y | A | I | E | A | T | C | - | H |
| | | *Pneumocystis jirovecii* | D | F | M | Q | M | P | - | F | S | D | N | Y | F | D | K | V | Y | S | I | E | A | T | V | - | H |
| Bacterial | (3) | Chlamydiae sp., drinking water MAG 1 of 2 | N | F | M | H | V | D | - | L | P | D | A | S | F | D | A | I | Y | T | I | E | A | T | V | - | H |
| | | *Aplysina aerophoba* meta JGIcombinedJ30088_10000835 1 of | D | Y | M | Q | I | P | - | E | G | D | D | Q | Y | D | A | A | F | A | I | E | S | T | A | - | H |
| | | *Aplysina aerophoba* meta Ga0209021_10001462 meta | D | F | L | H | V | E | - | T | P | D | E | S | F | D | A | V | Y | S | I | E | A | T | C | - | C |
| | | **Aplysina aerophoba* meta JGIcombinedJ30088_1000330310 | D | F | M | L | L | D | - | A | P | D | E | S | F | D | A | V | Y | S | I | E | A | S | C | - | C |
| | (2) | **Aplysina aerophoba* meta Ga0209754_100026449 | D | F | M | N | V | D | - | A | P | D | N | T | F | D | A | V | Y | A | I | E | A | T | C | - | C |
| | | *Petrosia ficiformis* meta LXNJ01000389 | D | F | L | N | V | D | - | A | P | D | E | S | Y | D | A | I | Y | S | I | E | A | T | C | - | C |
| | | **Aplysina aerophoba* meta Ga0209544_100039960 | D | F | M | N | I | E | - | T | P | D | N | T | F | D | A | V | Y | D | I | E | A | T | C | - | H |
| | | *Nitrospira* sp., coral reef MAG 1 of 2 2682229274 | D | Y | M | Q | I | P | - | E | D | N | D | H | Y | D | A | A | F | A | L | E | S | M | P | - | H |
| | | *Sandaracinus* sp., marine MAG 1 of 2 2775843484 | D | F | M | N | M | P | - | F | E | E | G | E | F | D | A | V | Y | A | F | E | A | T | C | - | H |
| | | Freshwater, Lake Fryxell, Antarctica meta | D | F | M | N | I | P | - | R | A | D | G | S | V | D | A | A | Y | A | F | E | A | T | C | - | H |
| | | Western Arctic Ocean meta Ga0133547_104202862 | D | F | M | A | M | P | - | F | E | D | A | S | F | D | A | A | Y | A | I | E | A | T | C | - | H |
| | (1) | **Aplysina aerophoba* meta JGIcombinedJ30088_100127566 | D | F | T | A | V | D | - | E | P | D | E | S | F | D | A | I | Y | A | F | E | S | T | C | - | H |
| | | Chlamydiae sp., drinking water MAG 2 of 2 2619789261 | N | F | M | Q | V | D | - | L | P | N | G | S | F | D | A | I | Y | A | V | E | A | T | P | - | H |
| | | *Sandaracinus* sp., marine MAG 2 of 2 2775843485 | D | F | M | N | L | P | - | V | E | D | G | R | F | D | A | V | Y | T | I | E | A | S | C | - | H |
| | | Marine sediment, Gulf of Thailand meta | D | F | G | D | L | P | - | F | D | N | D | S | F | D | A | A | Y | A | I | E | S | T | C | - | H |
| | | Spirochaetes sp., soil MAG 2709604172 | D | F | M | H | L | P | - | L | D | D | G | S | F | D | A | A | Y | T | I | E | A | S | C | - | H |
| | | Hotspring sediment, Dewer Creek, BC meta | D | W | M | Q | L | P | - | L | G | A | A | T | L | D | A | A | Y | T | I | E | A | S | C | - | H |
| | | Lake sediment, Walker Lake, Nevada meta | D | F | T | R | M | P | - | V | A | A | A | S | F | D | A | A | Y | A | F | E | A | T | C | - | H |
| | | Marine sediment, Helgoland, North Sea meta | D | F | M | N | L | Q | - | V | S | D | A | F | Y | D | A | A | Y | Q | I | E | A | T | C | - | H |
| | | *Sarcotragus foetidus* meta LXNI01004480 | D | Y | M | R | I | P | - | E | G | G | D | C | Y | D | A | A | F | A | I | E | S | M | P | - | H |
| | | *Theonella swinhoei* associated Chromatiales sp. MAG 1 of 2 | D | F | M | Q | I | P | - | E | D | D | N | S | Y | D | A | A | I | A | I | E | S | M | P | - | H |
| | | *Petrosia ficiformis* meta LXNJ01003202 | D | Y | M | Q | I | P | - | D | N | D | E | H | Y | D | G | A | I | N | I | E | A | M | P | - | H |
| | | *Petrosia ficiformis* meta LXNJ01014139 | D | Y | M | Q | I | P | - | E | G | D | D | C | Y | D | A | A | F | E | I | E | A | T | A | - | H |
| | No activity | *Aplysina aerophoba* meta JGIcombinedJ30088_10000835 2 of | S | F | M | D | M | S | A | I | K | D | E | T | F | D | R | G | Y | A | I | E | S | T | C | - | H |
| | | *Petrosia ficiformis* meta LXNJ01004423 | N | F | M | D | M | S | A | V | E | A | N | S | L | D | A | A | Y | A | I | E | S | T | C | - | H |
| | | *Theonella swinhoei* associated Chromatiales sp. MAG 2 of 2 | N | F | M | D | M | S | P | I | E | A | D | S | F | D | A | G | Y | A | I | E | S | T | C | - | H |
| | | Hotspring, Beatty, Nevada meta_Ga0114945_100210811 | D | F | M | E | I | P | - | P | D | D | G | P | Y | D | A | V | Y | A | I | E | A | T | C | - | H |
| | | *Nitrospira* sp., coral reef MAG 2 of 2 2682229270 | S | F | M | D | M | S | A | I | K | D | D | T | F | N | R | G | Y | A | I | E | S | T | C | - | H |
| | | **Aplysina aerophoba* meta JGIcombinedJ30088_100153964 | D | F | M | N | I | E | - | A | P | D | N | H | F | D | A | V | Y | S | I | E | A | S | C | - | C |
| | | **Aplysina aerophoba meta* JGIcombinedJ30088_100077196 | S | Y | L | N | V | D | - | T | P | D | E | S | F | D | G | A | Y | A | I | E | S | M | C | - | Y |
| | | **Aplysina aerophoba meta* JGIcombinedJ30088_100209763 | D | F | M | H | M | D | - | A | P | D | N | S | F | D | A | A | Y | A | I | E | A | T | I | - | H |
| | | **Aplysina aerophoba meta* JGIcombinedJ30088_100057164 | D | F | M | K | M | D | - | F | P | D | E | S | F | D | A | V | F | S | I | E | T | T | C | - | C |
| | | **Sarcotragus foetidus* meta LXNI01006544 | D | F | M | Q | V | D | - | A | P | D | A | S | F | D | A | A | Y | M | V | E | A | S | G | - | H |
| | | **Sarcotragus foetidus* meta LXNI01016685 | D | F | M | H | M | D | - | A | E | D | D | S | F | D | A | A | Y | S | I | E | A | T | A | P | H |
| | | Consensus | D | F | M | N | M | P | - | F | P | D | N | S | F | D | A | A | Y | A | I | E | A | T | C | - | H |
| | | Consensus # | 251 | 252 | 253 | 254 | 255 | 256 | 257 | 258 | 259 | 260 | 261 | 262 | 263 | 264 | 265 | 266 | 267 | 268 | 269 | 270 | 271 | 272 | 273 | 274 | 275 |

| | | | Methyltransferase domain | | | | | | | | | | | | | | | Region IV | | | | | | | | | | |
|---|---|---|---|---|---|---|---|---|---|---|---|---|---|---|---|---|---|---|---|---|---|---|---|---|---|---|---|---|
| | | ***S. cerevisiae* #** | **200** | **201** | **202** | **203** | **204** | **205** | **206** | **207** | **208** | **209** | **210** | **211** | **212** | **213** | **214** | **215** | **216** | **217** | **218** | **219** | **220** | **221** | **222** | **223** | **224** | **225** |
| Eukaryotic | (1) | *Saccharomyces cerevisiae* | A | P | K | L | E | G | V | Y | S | E | I | Y | K | V | L | K | P | **G** | G | T | F | A | V | Y | E | W |
| | | *Trypanosoma brucei* | S | E | S | K | V | K | C | Y | S | E | V | F | R | A | I | K | P | **G** | A | Y | F | M | L | Y | E | W |
| | | *Arabidopsis thaliana* SMT1 | A | P | D | A | Y | G | C | Y | K | E | I | Y | R | V | L | K | P | **G** | Q | C | F | A | A | Y | E | W |
| | (2) | *Arabidopsis thaliana* SMT2 | A | P | K | L | E | E | V | Y | A | E | I | Y | R | V | L | K | P | **G** | S | M | Y | V | S | Y | E | W |
| | | *Glycine max* SMT1 | A | P | D | A | Y | G | C | Y | K | E | I | F | R | V | L | K | P | **G** | Q | C | F | A | A | Y | E | W |
| | | *Chlamydomonas reinhardtii* | A | P | K | L | E | Q | V | Y | G | E | I | Y | R | V | L | K | P | **G** | S | Y | F | V | S | Y | E | W |
| | | *Pneumocystis jirovecii* | A | P | S | L | E | G | V | Y | G | E | A | F | R | V | L | K | P | **G** | G | I | F | A | L | Y | E | W |
| Bacterial | (3) | Chlamydiae sp., drinking water MAG 1 of2 | A | P | D | R | V | A | C | Y | K | E | M | Y | R | L | L | K | P | **G** | H | L | F | G | G | Y | E | Y |
| | | *Aplysina aerophoba* meta JGIcombinedJ30088_10000835 1 o | A | P | D | K | T | A | A | F | R | E | I | F | R | V | L | R | P | **G** | A | C | F | A | G | Y | E | W |
| | | *Aplysina aerophoba* meta Ga0209021_10001462 meta | A | P | D | K | L | S | I | Y | G | E | I | F | R | V | L | K | P | **G** | A | R | F | A | A | Y | E | Y |
| | | **Aplysina aerophoba* meta JGIcombinedJ30088_1000330310 | A | P | D | K | A | G | V | Y | S | E | A | F | R | V | L | K | P | **G** | A | L | L | G | V | Y | E | Y |
| | (2) | **Aplysina aerophoba* meta Ga0209754_100026449 | A | P | D | K | A | G | V | F | G | E | A | F | R | L | L | K | P | **G** | G | R | F | G | A | Y | D | F |
| | | *Petrosia ficiformis* meta LXNJ01000389 | A | P | N | K | T | S | T | Y | G | E | I | L | R | L | L | K | P | **G** | A | R | F | A | V | Y | E | Y |
| | | **Aplysina aerophoba* meta Ga0209544_100039960 | A | P | N | K | L | G | V | Y | S | E | V | F | R | V | L | K | P | **G** | G | C | F | G | A | Y | E | Y |
| | | *Nitrospira* sp., coral reef MAG 1 of2 2682229274 | A | P | D | K | T | A | V | F | R | E | I | F | R | V | L | R | Q | **G** | A | C | F | T | G | C | E | W |
| | | *Sandaracinus* sp., marine MAG 1 of2 2775843484 | A | P | D | K | A | A | L | F | R | Q | I | H | R | V | L | K | P | **G** | G | V | F | G | A | Y | E | W |
| | | Freshwater, Lake Fryxell, Antarctica meta | A | P | D | K | T | A | L | F | R | E | L | Y | R | V | L | K | P | **G** | G | K | L | A | I | Y | E | W |
| | | Western Arctic Ocean meta Ga0133547_104202862 | A | P | D | R | R | G | V | Y | G | E | I | F | R | T | L | K | P | **G** | G | V | F | A | S | Y | E | W |
| | (1) | **Aplysina aerophoba* meta JGIcombinedJ30088_100127566 | A | P | D | R | V | D | V | F | A | E | A | F | R | L | L | K | P | **G** | G | C | F | L | S | F | E | W |
| | | Chlamydiae sp., drinking water MAG 2 of2 2619789261 | A | P | D | R | T | A | C | F | K | E | M | Y | R | L | L | K | P | **G** | C | C | F | A | G | Y | E | Y |
| | | *Sandaracinus* sp., marine MAG 2 of2 2775843485 | A | P | D | R | V | K | L | F | R | E | L | Y | R | V | M | K | P | **G** | A | L | F | G | G | Y | E | W |
| | | Marine sediment, Gulf of Thailand meta | A | A | S | R | V | Q | V | F | K | E | I | A | R | V | L | K | P | **G** | G | E | F | A | G | Y | E | W |
| | | Spirochaetes sp., soil MAG 2709604172 | A | P | D | R | L | A | L | F | K | E | V | Y | R | I | L | R | P | **G** | G | Q | F | S | G | Y | E | W |
| | | Hotspring sediment, Dewer Creek, BC meta | A | S | D | R | G | A | L | F | G | E | V | R | R | V | L | K | A | **G** | A | P | F | V | G | Y | E | W |
| | | Lake sediment, Walker Lake, Nevada meta | A | P | D | K | A | A | L | F | A | E | V | R | R | V | L | R | P | **G** | A | E | F | A | L | Y | E | W |
| | | Marine sediment, Helgoland, North Sea meta | A | P | N | K | T | G | A | F | S | E | I | H | R | V | L | K | P | **G** | A | L | F | A | G | Y | E | W |
| | | *Sarcotragus foetidus* meta LXNI01004480 | A | P | D | K | V | G | A | F | R | E | I | L | R | V | L | R | P | **G** | A | C | F | A | G | Y | E | W |
| | | *Theonella swinhoei* associated Chromatiales sp. MAG 1 of2 | A | P | D | K | I | A | A | F | R | E | V | F | R | I | L | R | P | **G** | A | C | F | A | G | Y | E | W |
| | | *Petrosia ficiformis* meta LXNJ01003202 | A | P | D | K | I | A | A | Y | S | E | I | F | R | I | L | R | P | **G** | A | A | F | A | G | Y | E | W |
| | | *Petrosia ficiformis* meta LXNJ01014139 | A | P | D | K | A | G | A | F | G | E | V | L | R | V | L | R | P | **G** | A | C | F | A | G | Y | E | W |
| | No activity | *Aplysina aerophoba* meta JGIcombinedJ30088_10000835 2 o | A | P | D | K | P | R | A | F | A | E | I | F | R | V | L | K | P | G | A | L | F | W | G | Q | D | M |
| | | *Petrosia ficiformis* meta LXNJ01004423 | A | P | D | K | E | G | A | F | A | E | I | F | R | V | L | K | P | G | A | L | F | W | G | Q | E | M |
| | | *Theonella swinhoei* associated Chromatiales sp. MAG 2 of2 | A | P | D | K | K | H | A | F | A | E | I | F | R | V | L | K | P | G | A | L | F | W | G | Q | E | M |
| | | Hotspring, Beatty, Nevada meta_Ga0114945_100210811 | A | P | D | K | A | R | L | F | A | S | L | F | R | V | M | R | P | G | A | C | F | G | G | Y | E | W |
| | | *Nitrospira* sp., coral reef MAG 2 of2 2682229270 | A | P | D | K | P | R | A | F | A | E | I | F | R | I | L | K | P | G | A | L | F | W | G | Q | E | M |
| | | **Aplysina aerophoba* meta JGIcombinedJ30088_100153964 | A | P | D | R | P | G | I | Y | G | E | V | F | R | I | L | K | P | G | A | C | L | A | A | Y | E | Y |
| | | **Aplysina aerophoba meta* JGIcombinedJ30088_100077196 | S | P | S | R | E | E | A | Y | R | E | V | F | R | I | L | K | P | G | A | L | F | A | T | Y | E | V |
| | | **Aplysina aerophoba meta* JGIcombinedJ30088_100209763 | A | P | S | K | V | G | C | Y | G | E | V | F | R | V | L | K | P | G | G | C | F | A | A | Y | E | Y |
| | | **Aplysina aerophoba meta* JGIcombinedJ30088_100057164 | A | P | D | K | V | G | V | F | G | E | A | F | R | L | L | K | P | G | A | R | F | G | V | Y | E | Y |
| | | **Sarcotragus foetidus* meta LXNI01006544 | A | P | D | R | A | A | L | F | A | E | V | F | R | L | L | K | P | G | C | G | L | A | S | Y | E | W |
| | | **Sarcotragus foetidus* meta LXNI01016685 | A | P | S | K | V | G | C | Y | A | E | V | F | R | V | L | R | P | G | G | C | F | A | A | Y | E | Y |
| | | Consensus | A | P | D | K | V | G | V | F | G | E | I | F | R | V | L | K | P | **G** | A | C | F | A | G | Y | E | W |
| | | **Consensus #** | **276** | **277** | **278** | **279** | **280** | **281** | **282** | **283** | **284** | **285** | **286** | **287** | **288** | **289** | **290** | **291** | **292** | **293** | **294** | **295** | **296** | **297** | **298** | **299** | **300** | **301** |

| | | S. cerevisiae # | 226 | 227 | 228 | 229 | 230 | 231 | 232 | 233 | 234 | 235 | 236 | 237 | 238 | 239 | 240 | 241 | 242 | 243 | 244 | 245 | 246 | 247 | 248 | 249 |
|---|---|---|---|---|---|---|---|---|---|---|---|---|---|---|---|---|---|---|---|---|---|---|---|---|---|---|
| Eukaryotic | (1) | *Saccharomyces cerevisiae* | V | M | T | D | K | Y | D | E | N | N | P | E | H | R | K | I | A | Y | E | I | E | L | G | D |
| | | *Trypanosoma brucei* | C | L | T | D | L | Y | D | P | A | N | E | E | H | Q | R | V | R | H | G | I | E | L | G | D |
| | | *Arabidopsis thaliana* SMT1 | C | M | T | D | A | F | D | P | D | N | A | E | H | Q | K | I | K | G | E | I | E | I | G | D |
| | (2) | *Arabidopsis thaliana* SMT2 | V | T | T | E | K | F | K | A | E | D | D | E | H | V | E | V | I | Q | G | I | E | R | G | D |
| | | *Glycine max* SMT1 | C | M | T | D | S | F | D | P | Q | N | P | E | H | Q | K | I | K | A | E | I | E | I | G | D |
| | | *Chlamydomonas reinhardtii* | V | S | T | Q | K | F | D | V | N | N | A | E | H | V | K | I | M | D | E | I | N | F | G | N |
| | | *Pneumocystis jirovecii* | V | L | L | D | K | Y | D | E | T | N | P | E | H | R | K | I | V | Y | G | I | E | I | G | D |
| Bacterial | (3) | Chlamydiae sp., drinking water MAG 1 of2 | A | L | L | D | N | F | D | P | K | N | P | E | H | I | A | I | I | E | D | M | E | H | G | G |
| | | *Aplysina aerophoba* meta JGIcombinedJ30088_10000835 1 of | C | L | T | D | E | Y | D | P | E | N | A | K | H | L | R | I | K | N | D | I | M | V | G | G |
| | | *Aplysina aerophoba* meta Ga0209021_10001462 meta | C | M | T | D | R | F | D | P | Q | S | P | V | Q | L | K | L | K | G | E | I | E | L | G | G |
| | | **Aplysina aerophoba* meta JGIcombinedJ30088_1000330310 | C | M | T | D | L | F | D | P | E | N | P | E | Q | L | K | L | K | S | D | M | E | Y | S | G |
| | (2) | **Aplysina aerophoba* meta Ga0209754_100026449 | C | L | T | D | L | F | D | A | Q | D | P | L | H | L | K | L | K | A | D | M | E | Y | G | G |
| | | *Petrosia ficiformis* meta LXNJ01000389 | C | M | T | D | R | F | D | P | N | N | P | R | H | L | K | L | K | A | D | I | E | L | G | G |
| | | **Aplysina aerophoba* meta Ga0209544_100039960 | C | L | T | D | R | F | D | P | N | D | A | Y | H | L | K | L | K | A | D | I | E | L | G | G |
| | | *Nitrospira* sp., coral reef MAG 1 of2 2682229274 | C | L | T | E | N | F | D | P | R | N | A | E | H | L | R | I | K | N | D | I | V | T | G | N |
| | | *Sandaracinus* sp., marine MAG 1 of2 2775843484 | C | L | T | D | R | Y | D | P | E | S | V | R | H | R | R | I | K | K | G | I | E | E | G | D |
| | | Freshwater, Lake Fryxell, Antarctica meta | C | L | T | S | S | Y | D | A | R | V | A | T | H | A | Q | V | K | R | D | I | E | E | G | N |
| | | Western Arctic Ocean meta Ga0133547_104202862 | C | V | T | D | A | Y | D | R | E | D | P | A | Q | R | E | A | V | A | A | I | E | E | G | D |
| | (1) | **Aplysina aerophoba* meta JGIcombinedJ30088_100127566 | C | L | T | D | R | F | D | V | G | D | P | H | H | C | Q | L | K | H | D | I | E | L | G | G |
| | | Chlamydiae sp., drinking water MAG 2 of2 2619789261 | A | L | L | P | N | Y | D | A | N | N | P | E | H | I | A | I | M | E | D | I | E | H | G | G |
| | | *Sandaracinus* sp., marine MAG 2 of2 2775843485 | C | L | T | D | R | Y | D | E | S | N | A | E | H | R | T | I | K | K | D | I | E | I | G | N |
| | | Marine sediment, Gulf of Thailand meta | C | L | T | G | Q | F | R | L | D | L | R | F | H | K | A | L | K | V | L | I | E | K | G | D |
| | | Spirochaetes sp., soil MAG 2709604172 | C | L | T | D | L | Y | D | D | K | N | P | D | H | R | Q | I | K | K | G | I | E | E | G | D |
| | | Hotspring sediment, Dewer Creek, BC meta | V | L | T | P | A | Y | D | P | S | D | P | R | H | R | R | Y | R | A | G | I | E | E | G | S |
| | | Lake sediment, Walker Lake, Nevada meta | C | L | T | H | R | Y | D | P | T | D | A | D | H | Q | R | I | K | Q | A | V | E | E | G | D |
| | | Marine sediment, Helgoland, North Sea meta | C | M | T | D | A | F | D | P | A | D | P | R | H | Q | W | I | K | N | E | I | E | T | G | D |
| | | *Sarcotragus foetidus* meta LXNI01004480 | C | L | T | D | D | F | D | P | E | N | A | E | H | Q | W | I | K | D | E | I | M | V | G | D |
| | | *Theonella swinhoei* associated Chromatiales sp. MAG 1 of2 | C | L | T | E | D | F | D | P | E | N | A | R | H | L | R | I | K | K | D | I | E | K | G | N |
| | | *Petrosia ficiformis* meta LXNJ01003202 | C | L | T | D | V | F | D | P | S | N | A | E | H | L | R | I | K | E | G | I | V | T | G | S |
| | | *Petrosia ficiformis* meta LXNJ01014139 | C | L | T | E | K | F | E | P | A | N | A | D | H | L | R | I | K | N | D | I | M | V | G | N |
| | No activity | *Aplysina aerophoba* meta JGIcombinedJ30088_10000835 2 of | C | L | T | D | K | Y | D | P | N | D | S | R | H | R | A | I | K | R | E | L | M | R | G | I |
| | | *Petrosia ficiformis* meta LXNJ01004423 | C | M | T | E | K | Y | D | P | D | S | R | R | H | R | N | I | K | D | E | L | M | L | G | I |
| | | *Theonella swinhoei* associated Chromatiales sp. MAG 2 of2 | C | L | T | D | E | F | D | P | S | N | S | R | H | L | S | I | K | D | E | L | M | Q | G | I |
| | | Hotspring, Beatty, Nevada meta_Ga0114945_100210811 | C | L | T | P | R | Y | D | P | A | D | A | G | H | R | A | I | K | R | G | I | E | E | G | D |
| | | *Nitrospira* sp., coral reef MAG 2 of2 2682229270 | C | L | T | D | K | Y | D | P | N | D | S | R | H | R | A | I | K | R | E | L | M | R | G | I |
| | | **Aplysina aerophoba* meta JGIcombinedJ30088_100153964 | C | T | T | D | R | Y | N | A | E | D | P | H | H | R | R | L | K | A | D | M | E | Y | S | G |
| | | **Aplysina aerophoba meta* JGIcombinedJ30088_100077196 | C | L | T | D | K | Y | D | P | E | N | P | R | H | V | A | L | Y | S | Q | I | E | F | Y | G |
| | | **Aplysina aerophoba meta* JGIcombinedJ30088_100209763 | C | L | T | G | R | F | D | Q | N | D | P | R | H | R | Q | I | K | T | D | L | E | V | G | G |
| | | **Aplysina aerophoba meta* JGIcombinedJ30088_100057164 | C | L | T | D | L | F | D | P | E | D | A | N | H | Q | R | L | K | A | D | L | E | L | G | G |
| | | **Sarcotragus foetidus* meta LXNI01006544 | C | M | T | D | S | F | D | P | D | D | P | H | H | A | E | V | R | A | Y | A | Q | T | G | – |
| | | **Sarcotragus foetidus* meta LXNI01016685 | C | M | T | S | R | F | D | P | N | N | P | R | H | R | Q | I | K | N | D | I | E | V | G | G |
| | | Consensus | C | L | T | D | R | F | D | P | E | N | P | E | H | R | R | I | K | A | D | I | E | L | G | X |
| | | **Consensus #** | **302** | **303** | **304** | **305** | **306** | **307** | **308** | **309** | **310** | **311** | **312** | **313** | **314** | **315** | **316** | **317** | **318** | **319** | **320** | **321** | **322** | **323** | **324** | **325** |

| | | *S.cerevisiae* # | 250 | 251 | 252 | 253 | 254 | 255 | 256 | 257 | 258 | 259 | 260 | 261 | 262 | 263 | 264 | 265 | 266 | 267 | 268 | 269 | 270 | 271 | 272 | 273 | 274 |
|---|---|---|---|---|---|---|---|---|---|---|---|---|---|---|---|---|---|---|---|---|---|---|---|---|---|---|---|
| Eukaryotic | (1) | *Saccharomyces cerevisiae* | G | I | P | K | M | F | H | V | D | V | A | R | K | A | L | K | N | C | G | F | E | V | L | V | S |
| | | *Trypanosoma brucei* | G | L | P | E | L | D | T | M | R | Q | V | V | A | A | V | K | A | A | G | F | V | V | E | E | S |
| | | *Arabidopsis thaliana* SMT1 | G | L | P | D | I | R | L | T | T | K | C | L | E | A | L | K | Q | A | G | F | E | V | I | W | E |
| | (2) | *Arabidopsis thaliana* SMT2 | A | L | P | G | L | R | A | Y | V | D | I | A | E | T | A | K | K | V | G | F | E | I | V | K | E |
| | | *Glycine max* SMT1 | G | L | P | D | I | R | L | T | A | K | C | L | E | A | L | K | Q | A | G | F | E | V | I | W | E |
| | | *Chlamydomonas reinhardtii* | G | L | P | E | M | R | T | W | K | E | A | E | D | A | G | K | N | V | G | F | E | L | V | M | S |
| | | *Pneumocystis jirovecii* | S | I | P | K | M | F | K | V | S | E | S | E | A | A | L | R | K | V | G | F | E | I | I | H | S |
| Bacterial | (3) | Chlamydiae sp., drinking water MAG 1 of 2 | G | L | Q | K | V | L | P | L | K | E | L | K | Q | T | F | I | D | A | G | F | K | V | L | A | F |
| | | *Aplysina aerophoba* meta JGIcombinedJ30088_10000835 1 of | G | L | P | D | I | A | V | T | S | K | I | C | T | A | L | R | T | A | G | F | E | L | L | E | A |
| | | *Aplysina aerophoba* meta Ga0209021_10001462 meta | S | L | L | N | I | D | D | P | Q | T | V | V | D | A | L | R | S | V | G | F | E | V | L | E | T |
| | | **Aplysina aerophoba* meta JGIcombinedJ30088_1000330310 | A | L | P | N | I | A | R | P | H | E | V | D | D | A | M | R | A | V | G | F | E | L | L | E | A |
| | (2) | **Aplysina aerophoba* meta Ga0209754_100026449 | G | L | P | D | I | A | R | P | H | E | M | D | D | A | M | R | R | V | G | F | D | L | L | D | A |
| | | *Petrosia ficiformis* meta LXNJ01000389 | G | L | I | N | I | D | E | R | Q | T | V | D | D | V | L | H | A | L | G | F | E | V | L | E | T |
| | | **Aplysina aerophoba* meta Ga0209544_100039960 | S | L | L | D | I | D | D | C | P | T | V | D | Q | A | L | H | D | A | G | F | E | V | L | E | T |
| | | *Nitrospira* sp., coral reef MAG 1 of 2 2682229274 | A | L | P | D | I | A | V | T | S | E | I | G | T | A | L | R | M | A | G | F | E | L | L | E | S |
| | | *Sandaracinus* sp., marine MAG 1 of 2 2775843484 | G | L | P | D | I | W | T | I | P | A | T | V | E | C | L | E | G | A | G | F | T | V | R | E | Q |
| | | Freshwater, Lake Fryxell, Antarctica meta | G | L | P | D | I | W | T | I | D | A | T | R | A | C | F | E | Q | A | G | F | Q | V | L | G | E |
| | | Western Arctic Ocean meta Ga0133547_104202862 | A | L | P | E | L | V | H | T | S | V | V | D | S | A | L | A | E | V | G | F | E | L | L | H | T |
| | (1) | **Aplysina aerophoba* meta JGIcombinedJ30088_100127566 | G | L | Q | D | I | V | S | L | H | E | V | D | T | A | L | D | K | V | G | F | E | V | L | E | A |
| | | Chlamydiae sp., drinking water MAG 2 of 2 2619789261 | G | L | Q | K | I | P | P | I | A | E | L | K | Q | A | F | I | D | A | G | F | K | I | L | A | L |
| | | *Sandaracinus* sp., marine MAG 2 of 2 2775843485 | G | L | P | D | L | S | H | T | S | V | V | D | A | A | L | E | E | A | G | F | E | L | V | E | T |
| | | Marine sediment, Gulf of Thailand meta | A | L | P | E | L | A | F | T | S | D | V | D | D | A | L | R | E | A | G | L | E | I | V | E | A |
| | | Spirochaetes sp., soil MAG 2709604172 | A | L | P | D | I | L | H | T | S | E | I | V | R | V | L | K | E | A | G | F | T | V | C | D | S |
| | | Hotspring sediment, Dewer Creek, BC meta | G | V | P | P | L | A | T | Q | A | D | I | E | G | A | I | T | D | S | G | L | E | L | L | E | C |
| | | Lake sediment, Walker Lake, Nevada meta | G | L | P | D | I | W | T | I | T | A | T | L | D | A | A | R | D | A | G | F | E | L | V | E | H |
| | | Marine sediment, Helgoland, North Sea meta | S | L | P | D | I | A | S | I | P | E | T | I | R | A | L | E | A | A | G | F | E | V | L | D | S |
| | | *Sarcotragus foetidus* meta LXNI01004480 | A | L | P | E | I | A | S | M | P | E | I | C | D | A | L | R | T | A | G | F | E | L | L | E | F |
| | | *Theonella swinhoei* associated Chromatiales sp. MAG 1 of 2 | G | L | P | D | I | T | I | T | S | E | I | G | P | A | L | S | A | A | G | F | E | F | L | E | A |
| | | *Petrosia ficiformis* meta LXNJ01003202 | G | L | P | D | I | A | S | T | E | E | V | C | D | S | L | R | A | V | G | F | E | V | L | E | A |
| | | *Petrosia ficiformis* meta LXNJ01014139 | G | L | P | D | I | A | S | T | S | E | V | S | A | A | L | Q | A | V | G | F | E | L | L | D | A |
| | No activity | *Aplysina aerophoba* meta JGIcombinedJ30088_10000835 2 of | A | L | K | D | I | A | T | F | G | E | V | N | R | A | L | E | A | A | G | F | Q | V | I | E | G |
| | | *Petrosia ficiformis* meta LXNJ01004423 | A | L | D | G | I | A | T | F | A | E | V | N | R | A | L | E | A | V | G | F | R | I | V | E | A |
| | | *Theonella swinhoei* associated Chromatiales sp. MAG 2 of 2 | A | L | S | E | I | A | T | F | A | E | V | N | Q | A | L | E | A | A | G | F | Q | V | I | E | A |
| | | Hotspring, Beatty, Nevada meta_Ga0114945_100210811 | A | L | P | E | L | A | T | F | D | A | V | L | D | A | L | G | R | A | G | F | D | V | L | E | S |
| | | *Nitrospira* sp., coral reef MAG 2 of 2 2682229270 | A | L | K | D | I | A | T | F | R | E | V | N | H | A | L | E | A | A | G | F | Q | V | I | E | G |
| | | **Aplysina aerophoba* meta JGIcombinedJ30088_100153964 | C | I | P | D | I | A | R | P | R | E | I | V | D | T | L | R | Q | V | R | F | E | L | L | D | A |
| | | **Aplysina aerophoba meta* JGIcombinedJ30088_100077196 | G | M | T | D | L | D | L | P | R | D | V | D | S | A | L | K | S | V | G | F | D | L | V | E | N |
| | | **Aplysina aerophoba meta* JGIcombinedJ30088_100209763 | A | L | P | E | I | A | T | P | H | Q | V | D | D | A | L | R | E | V | G | F | E | L | L | E | S |
| | | **Aplysina aerophoba meta* JGIcombinedJ30088_100057164 | G | L | P | D | I | A | Y | P | P | E | V | D | G | A | L | R | T | V | G | F | E | L | L | A | A |
| | | **Sarcotragus foetidus* meta LXNI01006544 | – | V | H | D | C | P | G | T | E | A | V | D | Q | A | L | S | E | A | G | F | E | I | L | E | T |
| | | **Sarcotragus foetidus* meta LXNI01016685 | G | L | Q | D | I | V | F | P | H | Q | L | D | E | A | L | R | Q | A | G | F | E | L | L | E | G |
| | | Consensus | G | L | P | D | I | A | T | T | S | E | V | D | D | A | L | R | A | A | G | F | E | V | L | E | A |
| | | **Consensus #** | 326 | 327 | 328 | 329 | 330 | 331 | 332 | 333 | 334 | 335 | 336 | 337 | 338 | 339 | 340 | 341 | 342 | 343 | 344 | 345 | 346 | 347 | 348 | 349 | 350 |

| | | S.cerevisiae # | 275 | 276 | 277 | 278 | 279 | 280 | | | | | | 281 | 282 | 283 | 284 | 285 | 286 | 287 | 288 | 289 | 290 | 291 | 292 | 293 | 294 |
|---|---|---|---|---|---|---|---|---|---|---|---|---|---|---|---|---|---|---|---|---|---|---|---|---|---|---|---|
| Eukaryotic | (1) | *Saccharomyces cerevisiae* | E | D | L | A | D | N | - | - | - | - | - | D | D | E | I | P | W | Y | Y | P | L | T | G | E | W |
| | | *Trypanosoma brucei* | F | D | M | A | E | R | F | E | S | G | E | P | K | S | V | P | W | Y | E | P | L | Q | G | S | Y |
| | | *Arabidopsis thaliana* SMT1 | K | D | L | A | K | - | - | - | - | - | - | D | S | P | V | P | W | Y | L | P | L | D | K | N | H |
| | (2) | *Arabidopsis thaliana* SMT2 | K | D | L | A | S | - | - | - | - | - | - | P | P | A | E | P | W | W | T | R | L | - | - | - | - |
| | | *Glycine max* SMT1 | K | D | L | A | V | - | - | - | - | - | - | D | S | P | L | P | W | Y | L | P | L | D | K | S | H |
| | | *Chlamydomonas reinhardtii* | L | D | L | A | T | A | S | - | - | - | - | V | V | A | G | P | W | E | G | A | S | S | T | G | S |
| | | *Pneumocystis jirovecii* | E | E | L | S | T | K | - | - | - | - | - | N | N | E | I | P | W | Y | Y | L | L | D | T | D | F |
| Bacterial | (3) | Chlamydiae sp., drinking water MAG 1 of 2 | E | D | L | C | K | E | - | - | - | - | - | - | - | G | L | T | W | T | L | P | L | E | - | - | - |
| | | *Aplysina aerophoba* meta JGIcombinedJ30088_10000835 1 of | C | D | R | A | P | E | S | - | - | - | - | D | P | E | M | P | W | Y | R | A | L | Q | G | R | D |
| | | *Aplysina aerophoba* meta Ga0209021_10001462 meta | R | D | L | A | L | Q | T | - | - | - | - | G | P | S | I | P | W | Y | E | P | L | V | G | S | G |
| | | **Aplysina aerophoba* meta JGIcombinedJ30088_1000330310 | R | D | L | A | A | D | A | - | - | - | - | P | P | G | I | P | W | Y | E | P | L | V | G | S | G |
| | (2) | **Aplysina aerophoba* meta Ga0209754_100026449 | R | D | L | A | A | D | A | - | - | - | - | P | P | G | I | P | W | Y | Q | P | L | V | G | S | G |
| | | *Petrosia ficiformis* meta LXNJ01000389 | R | D | L | A | E | Q | S | - | - | - | - | G | P | S | I | P | W | Y | Q | P | L | A | G | S | G |
| | | **Aplysina aerophoba* meta Ga0209544_100039960 | R | D | L | I | N | Q | P | - | - | - | - | G | P | S | I | P | W | Y | E | P | L | V | G | S | G |
| | | *Nitrospira* sp., coral reef MAG 1 of 2 2682229274 | H | D | L | A | P | E | S | - | - | - | - | H | P | E | M | P | W | Y | R | S | L | Q | G | R | D |
| | | *Sandaracinus* sp., marine MAG 1 of 2 2775843484 | E | D | R | A | V | A | S | - | - | - | - | D | P | E | T | P | W | E | L | P | I | T | G | R | E |
| | | Freshwater, Lake Fryxell, Antarctica meta | E | D | R | A | G | Q | G | - | - | - | - | D | A | E | L | P | W | Y | H | S | L | T | G | R | E |
| | | Western Arctic Ocean meta Ga0133547_104202862 | H | D | V | A | L | D | S | - | - | - | - | D | P | E | T | P | W | F | L | P | L | T | G | K | E |
| | (1) | **Aplysina aerophoba* meta JGIcombinedJ30088_100127566 | L | D | M | A | E | A | P | - | - | - | - | E | P | S | I | P | W | Y | Q | P | L | A | G | P | R |
| | | Chlamydiae sp., drinking water MAG 2 of 2 2619789261 | Q | D | D | C | T | E | - | - | - | - | - | - | - | G | L | T | W | T | L | P | L | E | - | - | - |
| | | *Sandaracinus* sp., marine MAG 2 of 2 2775843485 | E | D | V | A | E | T | A | - | - | - | - | D | E | E | T | P | W | Y | L | P | L | S | G | E | E |
| | | Marine sediment, Gulf of Thailand meta | F | D | C | A | S | E | G | - | - | - | - | D | P | K | R | R | W | Y | T | P | L | D | S | D | A |
| | | Spirochaetes sp., soil MAG 2709604172 | E | D | R | A | L | T | S | - | - | - | - | D | P | R | K | P | W | Y | L | P | I | E | G | K | E |
| | | Hotspring sediment, Dewer Creek, BC meta | R | D | L | A | P | E | C | - | - | - | - | D | P | A | T | P | W | Y | L | P | L | E | G | E | G |
| | | Lake sediment, Walker Lake, Nevada meta | E | D | R | A | A | T | C | - | - | - | - | D | H | E | T | P | W | Y | L | P | L | T | G | R | E |
| | | Marine sediment, Helgoland, North Sea meta | F | D | A | V | Q | R | S | - | - | - | - | D | N | E | T | P | W | Y | L | P | L | R | G | E | T |
| | | *Sarcotragus foetidus* meta LXNI01004480 | Q | D | R | A | S | E | S | - | - | - | - | D | P | E | T | P | W | Y | R | A | L | Q | G | R | D |
| | | *Theonella swinhoei* associated Chromatiales sp. MAG 1 of 2 | R | D | L | A | P | E | A | - | - | - | - | D | P | D | M | P | W | Y | R | A | L | Q | G | R | D |
| | | *Petrosia ficiformis* meta LXNJ01003202 | R | D | L | A | P | E | G | - | - | - | - | D | P | E | T | P | W | Y | R | A | L | Q | G | R | D |
| | | *Petrosia ficiformis* meta LXNJ01014139 | R | D | R | A | P | E | A | - | - | - | - | D | Q | E | M | P | W | Y | R | A | L | Q | G | R | D |
| | No activity | *Aplysina aerophoba* meta JGIcombinedJ30088_10000835 2 of | K | D | R | D | V | K | E | - | - | - | - | G | P | S | T | P | W | Y | Q | P | M | E | T | R | Y |
| | | *Petrosia ficiformis* meta LXNJ01004423 | L | D | R | D | V | Q | E | - | - | - | - | G | P | S | T | P | W | Y | R | P | M | E | G | R | S |
| | | *Theonella swinhoei* associated Chromatiales sp. MAG 2 of 2 | G | D | K | S | V | R | E | - | - | - | T | G | P | S | R | P | W | Y | E | P | M | E | I | V | S |
| | | Hotspring, Beatty, Nevada meta_Ga0114945_100210811 | R | D | L | A | P | E | A | - | - | - | - | D | A | G | T | P | W | Y | Q | P | L | A | A | R | - |
| | | *Nitrospira* sp., coral reef MAG 2 of 2 2682229270 | M | D | R | D | V | Q | E | - | - | - | - | G | P | S | T | P | W | Y | Q | P | M | E | T | R | H |
| | | **Aplysina aerophoba* meta JGIcombinedJ30088_100153964 | R | D | L | A | A | E | A | - | - | - | - | P | P | G | I | P | W | Y | Q | P | L | T | G | S | G |
| | | **Aplysina aerophoba meta* JGIcombinedJ30088_100077196 | R | D | I | A | E | D | D | - | - | - | - | R | F | E | I | P | W | Y | E | P | I | E | G | T | G |
| | | **Aplysina aerophoba meta* JGIcombinedJ30088_100209763 | R | D | L | A | E | K | P | - | - | - | - | G | P | G | N | P | W | Y | Q | P | L | A | G | T | R |
| | | **Aplysina aerophoba meta* JGIcombinedJ30088_100057164 | S | D | K | A | V | Q | D | - | - | - | - | A | R | G | I | P | W | Y | H | P | L | T | G | R | G |
| | | **Sarcotragus foetidus* meta LXNI01006544 | R | D | A | A | H | E | D | - | - | - | - | G | V | V | I | P | W | Y | E | A | L | N | G | S | G |
| | | **Sarcotragus foetidus* meta LXNI01016685 | A | D | L | N | E | L | A | - | - | - | - | E | P | G | I | P | W | Y | Q | P | L | A | G | S | R |
| | | Consensus | R | D | L | A | X | E | S | - | - | - | - | D | P | E | I | P | W | Y | L | P | L | E | G | R | G |
| | | **Consensus #** | 351 | 352 | 353 | 354 | 355 | 356 | 357 | 358 | 359 | 360 | 361 | 362 | 363 | 364 | 365 | 366 | 367 | 368 | 369 | 370 | 371 | 372 | 373 | 374 | 375 |

Sterol MT C-term domain

| | | S.cerevisiae # | 295 | | | | | | 296 | 297 | 298 | 299 | 300 | 301 | 302 | 303 | 304 | 305 | 306 | 307 | 308 | 309 | 310 | 311 | 312 | 313 | 314 |
|---|---|---|---|---|---|---|---|---|---|---|---|---|---|---|---|---|---|---|---|---|---|---|---|---|---|---|---|
| Eukaryotic | (1) | *Saccharomyces cerevisiae* | K | - | - | - | - | - | Y | V | Q | N | L | A | N | L | A | T | F | F | R | T | S | Y | L | G | R |
| | | *Trypanosoma brucei* | T | - | - | - | - | - | S | L | S | G | L | - | - | - | - | - | - | - | R | A | T | P | A | G | R |
| | | *Arabidopsis thaliana* SMT1 | F | - | - | - | - | - | S | L | S | S | F | - | - | - | - | - | - | - | R | L | T | A | V | G | R |
| | (2) | *Arabidopsis thaliana* SMT2 | - | - | - | - | - | - | - | - | - | - | - | - | - | - | - | - | - | - | K | M | G | R | L | A | Y |
| | | *Glycine max* SMT1 | F | - | - | - | - | - | S | L | S | S | F | - | - | - | - | - | - | - | R | L | T | A | V | G | R |
| | | *Chlamydomonas reinhardtii* | G | A | G | L | S | P | S | T | A | W | Y | - | - | - | - | E | R | L | R | M | G | K | Y | T | H |
| | | *Pneumocystis jirovecii* | R | - | - | - | - | - | K | V | R | S | L | W | D | L | F | S | I | A | R | M | T | W | L | G | R |
| Bacterial | (3) | Chlamydiae sp., drinking water MAG 1 of 2 | - | - | - | - | - | - | - | - | R | G | V | - | - | - | - | - | - | - | R | S | S | K | T | A | R |
| | | *Aplysina aerophoba* meta JGIcombinedJ30088_10000835 1 of | L | - | - | - | - | - | S | M | G | S | I | - | - | - | - | - | - | - | P | R | T | P | V | G | R |
| | | *Aplysina aerophoba* meta Ga0209021_10001462 meta | I | - | - | - | - | - | S | F | A | S | F | - | - | - | - | - | - | - | R | S | S | N | V | G | R |
| | | **Aplysina aerophoba* meta JGIcombinedJ30088_1000330310 | L | - | - | - | - | - | S | L | A | S | F | - | - | - | - | - | - | - | R | S | S | A | L | G | R |
| | (2) | **Aplysina aerophoba* meta Ga0209754_100026449 | L | - | - | - | - | - | S | F | A | S | A | - | - | - | - | - | - | - | R | S | S | P | V | G | R |
| | | *Petrosia ficiformis* meta LXNJ01000389 | L | - | - | - | - | - | S | F | V | G | F | - | - | - | - | - | - | - | R | S | S | R | I | G | R |
| | | **Aplysina aerophoba* meta Ga0209544_100039960 | M | - | - | - | - | - | S | V | A | S | F | - | - | - | - | - | - | - | R | S | T | R | I | G | R |
| | | *Nitrospira* sp., coral reef MAG 1 of 2 2682229274 | F | - | - | - | - | - | S | L | T | S | I | - | - | - | - | - | - | - | P | R | T | P | V | G | R |
| | | *Sandaracinus* sp., marine MAG 1 of 2 2775843484 | L | - | - | - | - | - | S | W | T | G | V | - | - | - | - | - | - | - | R | R | T | K | L | G | R |
| | | Freshwater, Lake Fryxell, Antarctica meta | L | - | - | - | - | - | S | N | T | G | I | - | - | - | - | - | - | - | R | R | S | P | V | G | K |
| | | Western Arctic Ocean meta Ga0133547_104202862 | L | - | - | - | - | - | S | I | R | G | L | - | - | - | - | - | - | - | P | H | T | R | P | G | R |
| | (1) | **Aplysina aerophoba* meta JGIcombinedJ30088_100127566 | L | - | - | - | - | - | S | L | T | G | L | - | - | - | - | - | - | - | R | S | S | R | V | G | R |
| | | Chlamydiae sp., drinking water MAG 2 of 2 2619789261 | - | - | - | - | - | - | - | - | K | G | M | - | - | - | - | - | - | - | R | S | S | K | T | G | R |
| | | *Sandaracinus* sp., marine MAG 2 of 2 2775843485 | K | - | - | - | - | - | G | L | R | A | F | - | - | - | - | - | - | - | P | R | T | P | L | G | R |
| | | Marine sediment, Gulf of Thailand meta | R | - | - | - | - | - | S | I | R | S | L | - | - | - | - | - | - | - | P | R | T | P | L | G | R |
| | | Spirochaetes sp., soil MAG 2709604172 | L | S | - | - | - | - | S | L | H | G | L | - | - | - | - | - | - | - | L | C | T | R | P | G | R |
| | | Hotspring sediment, Dewer Creek, BC meta | L | - | - | - | - | - | S | L | A | A | F | - | - | - | - | - | - | - | R | R | S | H | L | G | G |
| | | Lake sediment, Walker Lake, Nevada meta | R | - | - | - | - | - | S | V | I | G | L | - | - | - | - | - | - | - | R | R | S | I | I | G | Q |
| | | Marine sediment, Helgoland, North Sea meta | S | - | - | - | - | - | T | L | L | G | I | - | - | - | - | - | - | - | R | R | G | R | L | G | R |
| | | *Sarcotragus foetidus* meta LXNI01004480 | L | - | - | - | - | - | T | L | S | S | I | - | - | - | - | - | - | - | P | R | T | P | F | G | R |
| | | *Theonella swinhoei* associated Chromatiales sp. MAG 1 of 2 | L | - | - | - | - | - | S | L | A | S | I | - | - | - | - | - | - | - | P | R | T | P | I | G | R |
| | | *Petrosia ficiformis* meta LXNJ01003202 | F | - | - | - | - | - | S | L | A | S | I | - | - | - | - | - | - | - | P | R | T | P | F | G | R |
| | | *Petrosia ficiformis* meta LXNJ01014139 | L | - | - | - | - | - | S | L | S | S | I | - | - | - | - | - | - | - | P | R | T | P | V | G | R |
| | No activity | *Aplysina aerophoba* meta JGIcombinedJ30088_10000835 2 of | G | - | - | - | - | - | T | L | G | N | L | - | - | - | - | - | - | V | C | R | L | P | W | G | R |
| | | *Petrosia ficiformis* meta LXNJ01004423 | G | - | - | - | - | - | T | L | R | G | A | - | - | - | - | - | - | F | R | R | T | P | L | G | R |
| | | *Theonella swinhoei* associated Chromatiales sp. MAG 2 of 2 | G | - | - | - | - | - | P | L | G | S | F | - | - | - | - | - | - | S | R | K | T | L | L | G | R |
| | | Hotspring, Beatty, Nevada meta_Ga0114945_100210811 | V | - | - | - | - | - | A | L | R | G | F | - | - | - | - | - | - | - | H | Q | S | R | A | G | R |
| | | *Nitrospira* sp., coral reef MAG 2 of 2 2682229270 | G | - | - | - | - | - | T | L | G | N | A | - | - | - | - | - | - | L | C | R | L | P | W | G | R |
| | | **Aplysina aerophoba* meta JGIcombinedJ30088_100153964 | L | - | - | - | - | - | S | L | A | N | F | - | - | - | - | - | - | - | R | S | S | T | L | G | R |
| | | **Aplysina aerophoba meta* JGIcombinedJ30088_100077196 | F | - | - | - | - | - | S | L | E | S | F | - | - | - | - | - | - | - | R | S | S | L | I | G | R |
| | | **Aplysina aerophoba meta* JGIcombinedJ30088_100209763 | F | - | - | - | - | - | S | F | A | S | F | - | - | - | - | - | - | - | R | S | T | R | A | G | R |
| | | **Aplysina aerophoba meta* JGIcombinedJ30088_100057164 | M | - | - | - | - | - | S | L | A | N | F | - | - | - | - | - | - | - | R | S | S | K | V | G | R |
| | | **Sarcotragus foetidus* meta LXNI01006544 | W | - | - | - | - | - | S | R | A | A | F | - | - | - | - | - | - | - | R | R | S | R | V | G | R |
| | | **Sarcotragus foetidus* meta LXNI01016685 | F | - | - | - | - | - | S | F | A | S | F | - | - | - | - | - | - | - | R | S | T | S | G | G | R |
| | | Consensus | L | - | - | - | - | - | S | L | A | S | F | - | - | - | - | - | - | - | R | R | T | P | L | G | R |
| | | Consensus # | 376 | 377 | 378 | 379 | 380 | 381 | 382 | 383 | 384 | 385 | 386 | 387 | 388 | 389 | 390 | 391 | 392 | 393 | 394 | 395 | 396 | 397 | 398 | 399 | 400 |

#### Sterol MT C-term domain

| | | *S. cerevisiae* # | 315 | 316 | 317 | 318 | 319 | 320 | 321 | 322 | 323 | 324 | 325 | 326 | 327 | 328 | 329 | 330 | 331 | 332 | 333 | 334 | 335 | 336 | 337 | 338 | 339 |
|---|---|---|---|---|---|---|---|---|---|---|---|---|---|---|---|---|---|---|---|---|---|---|---|---|---|---|---|
| Eukaryotic | (1) | *Saccharomyces cerevisiae* | Q | F | T | T | A | M | V | T | V | M | E | K | L | G | L | A | P | E | G | S | K | E | V | T | A |
| | | *Trypanosoma brucei* | W | L | T | S | V | T | C | R | L | L | E | A | V | R | L | A | P | A | G | T | C | K | A | T | E |
| | | *Arabidopsis thaliana* SMT1 | F | I | T | K | N | M | V | K | I | L | E | Y | I | R | L | A | P | Q | G | S | Q | R | V | S | N |
| | (2) | *Arabidopsis thaliana* SMT2 | W | R | N | H | I | V | V | Q | I | L | S | A | V | G | V | A | P | K | G | T | V | D | V | H | E |
| | | *Glycine max* SMT1 | L | F | T | K | N | M | V | K | V | L | E | Y | V | G | L | A | P | K | G | S | L | R | V | Q | D |
| | | *Chlamydomonas reinhardtii* | A | I | N | H | G | I | V | S | T | V | D | A | L | G | L | A | P | K | G | L | K | E | V | H | H |
| | | *Pneumocystis jirovecii* | W | F | I | G | F | F | L | N | L | M | E | F | I | G | V | A | P | K | G | C | K | K | V | N | D |
| Bacterial | (3) | Chlamydiae sp., drinking water MAG 1 of 2 | A | L | T | N | G | M | V | R | L | L | E | F | L | R | I | A | P | K | G | S | T | S | V | S | S |
| | | *Aplysina aerophoba* meta JGIcombinedJ30088_10000835 1 of | A | L | T | N | L | V | L | R | V | G | E | K | L | R | L | V | P | E | G | A | R | E | V | S | T |
| | | *Aplysina aerophoba* meta Ga0209021_10001462 meta | W | V | T | L | N | S | V | R | V | L | E | T | L | R | I | A | P | K | G | S | V | R | V | L | E |
| | | **Aplysina aerophoba* meta JGIcombinedJ30088_1000330310 | R | L | T | H | G | S | L | W | L | L | E | R | L | R | I | V | P | R | G | S | F | R | V | S | A |
| | (2) | **Aplysina aerophoba* meta Ga0209754_100026449 | R | L | T | R | G | S | L | W | L | L | E | R | L | R | I | V | P | R | G | T | A | Q | V | A | A |
| | | *Petrosia ficiformis* meta LXNJ01000389 | A | I | T | H | G | T | L | K | A | L | E | A | L | R | I | A | P | K | G | S | A | R | V | A | H |
| | | **Aplysina aerophoba* meta Ga0209544_100039960 | E | L | T | T | N | I | L | R | V | L | E | T | L | R | V | V | P | Q | G | S | V | N | V | T | Q |
| | | *Nitrospira* sp., coral reef MAG 1 of 2 2682229274 | A | L | T | N | L | A | L | R | A | A | E | K | L | R | V | V | P | E | G | T | T | A | V | S | T |
| | | *Sandaracinus* sp., marine MAG 1 of 2 2775843484 | R | L | T | W | A | A | I | K | V | M | E | A | A | R | I | A | P | K | G | T | M | E | V | S | Q |
| | | Freshwater, Lake Fryxell, Antarctica meta | A | M | V | K | V | L | T | R | T | L | E | A | A | R | V | A | P | R | G | A | S | R | I | A | D |
| | | Western Arctic Ocean meta Ga0133547_104202862 | M | L | S | H | A | F | V | R | F | M | E | A | V | R | L | A | P | K | G | A | T | S | V | S | E |
| | (1) | **Aplysina aerophoba* meta JGIcombinedJ30088_100127566 | S | A | T | Y | Q | F | L | R | V | L | E | A | L | R | V | V | P | R | G | T | P | G | A | A | R |
| | | Chlamydiae sp., drinking water MAG 2 of 2 2619789261 | A | L | T | N | T | M | V | R | V | L | E | F | L | G | I | A | P | K | G | S | S | E | V | S | R |
| | | *Sandaracinus* sp., marine MAG 2 of 2 2775843485 | E | V | T | N | V | F | T | R | V | L | E | K | L | R | L | A | P | Q | G | T | S | Q | V | S | D |
| | | Marine sediment, Gulf of Thailand meta | W | L | M | H | H | L | T | W | I | L | E | R | T | G | V | A | A | P | G | T | N | E | V | Q | A |
| | | Spirochaetes sp., soil MAG 2709604172 | M | I | T | H | F | L | L | T | I | F | E | K | M | G | I | A | P | K | G | S | I | A | L | S | A |
| | | Hotspring sediment, Dewer Creek, BC meta | L | F | L | Q | G | A | V | G | A | L | E | L | T | R | V | L | P | R | G | A | G | A | V | I | R |
| | | Lake sediment, Walker Lake, Nevada meta | R | V | L | A | V | T | I | R | A | M | Q | A | L | G | A | A | P | R | G | T | L | E | T | L | R |
| | | Marine sediment, Helgoland, North Sea meta | R | I | A | R | R | T | I | G | A | L | E | A | L | G | I | A | P | K | G | S | K | E | V | S | Q |
| | | *Sarcotragus foetidus* meta LXNI01004480 | A | M | T | N | L | T | L | R | I | G | E | R | L | R | L | F | P | K | G | S | R | A | V | S | T |
| | | *Theonella swinhoei* associated Chromatiales sp. MAG 1 of 2 | A | L | T | N | L | T | L | R | V | G | E | T | L | R | V | V | P | K | G | S | T | M | V | S | T |
| | | *Petrosia ficiformis* meta LXNJ01003202 | A | L | T | N | L | T | L | G | V | A | E | R | L | H | L | A | P | K | G | A | R | A | V | S | S |
| | | *Petrosia ficiformis* meta LXNJ01014139 | A | L | T | N | L | T | L | R | V | G | E | R | L | R | L | F | P | E | G | A | R | A | V | S | T |
| | No activity | *Aplysina aerophoba* meta JGIcombinedJ30088_10000835 2 of | R | C | C | I | A | G | S | K | L | A | E | V | L | R | L | F | P | K | G | S | A | E | V | V | R |
| | | *Petrosia ficiformis* meta LXNJ01004423 | K | A | M | I | G | G | L | R | V | A | E | A | F | R | F | L | P | K | G | S | A | A | V | V | R |
| | | *Theonella swinhoei* associated Chromatiales sp. MAG 2 of 2 | R | M | I | I | G | F | F | R | L | T | E | A | M | R | I | F | P | K | G | S | I | E | I | V | Q |
| | | Hotspring, Beatty, Nevada meta_Ga0114945_100210811 | V | V | T | N | L | A | V | R | G | L | E | L | V | R | I | A | P | R | G | A | A | A | V | S | T |
| | | *Nitrospira* sp., coral reef MAG 2 of 2 2682229270 | K | V | F | M | A | G | S | K | L | A | E | V | L | R | L | F | P | K | G | S | A | E | V | V | R |
| | | **Aplysina aerophoba* meta JGIcombinedJ30088_100153964 | R | V | T | K | G | S | L | W | L | L | E | R | L | R | V | V | P | R | G | T | L | Q | V | A | K |
| | | **Aplysina aerophoba meta* JGIcombinedJ30088_100077196 | T | L | T | H | V | T | V | Q | V | L | E | K | L | K | I | A | P | Q | G | S | T | Q | V | S | T |
| | | **Aplysina aerophoba meta* JGIcombinedJ30088_100209763 | I | V | T | H | G | T | L | Q | L | M | E | A | L | G | I | V | P | R | G | T | V | R | V | S | G |
| | | **Aplysina aerophoba meta* JGIcombinedJ30088_100057164 | A | V | T | H | G | S | V | W | T | M | E | R | L | G | V | L | P | S | G | T | L | R | V | S | R |
| | | **Sarcotragus foetidus* meta LXNI01006544 | W | V | T | H | R | T | V | R | L | L | E | L | F | R | I | V | P | R | G | T | T | E | V | A | G |
| | | **Sarcotragus foetidus* meta LXNI01016685 | I | V | T | H | G | T | L | Q | L | L | E | A | L | G | I | V | P | K | G | T | V | R | V | H | G |
| | | Consensus | A | L | T | H | G | T | L | R | V | L | E | A | L | R | I | A | P | K | G | S | X | E | V | S | R |
| | | **Consensus #** | **401** | **402** | **403** | **404** | **405** | **406** | **407** | **408** | **409** | **410** | **411** | **412** | **413** | **414** | **415** | **416** | **417** | **418** | **419** | **420** | **421** | **422** | **423** | **424** | **425** |

**Sterol MT C-term domain**

| | | *S.cerevisiae* # | 340 | 341 | 342 | 343 | 344 | 345 | 346 | 347 | 348 | 349 | 350 | 351 | 352 | 353 | 354 | 355 | 356 | 357 | 358 | 359 | 360 | 361 | 362 | 363 | 364 |
|---|---|---|---|---|---|---|---|---|---|---|---|---|---|---|---|---|---|---|---|---|---|---|---|---|---|---|---|
| Eukaryotic | (1) | *Saccharomyces cerevisiae* | A | L | E | N | A | A | V | G | L | V | A | G | G | K | S | K | L | F | T | P | M | M | L | F | V |
| | | *Trypanosoma brucei* | I | L | E | E | G | A | V | N | L | V | K | G | G | E | L | G | I | F | T | P | S | F | F | V | K |
| | | *Arabidopsis thaliana* SMT1 | F | L | E | Q | A | A | E | G | L | V | D | G | G | R | R | E | I | F | T | P | M | Y | F | F | L |
| | (2) | *Arabidopsis thaliana* SMT2 | M | L | F | K | T | A | D | Y | L | T | R | G | G | E | T | G | I | F | S | P | M | H | M | I | L |
| | | *Glycine max* SMT1 | F | L | E | K | A | A | E | G | L | V | E | G | G | K | R | E | I | F | T | P | M | Y | F | F | L |
| | | *Chlamydomonas reinhardtii* | M | L | V | E | V | A | K | S | L | I | Q | G | G | E | S | G | I | F | T | P | M | H | L | L | L |
| | | *Pneumocystis jirovecii* | V | L | L | L | A | A | D | A | L | V | D | A | G | K | Q | Q | I | F | S | P | M | Q | M | W | V |
| Bacterial | (3) | Chlamydiae sp., drinking water MAG 1 of 2 | F | L | N | L | G | A | D | G | F | V | E | A | G | K | R | K | L | F | T | P | N | V | F | F | L |
| | | *Aplysina aerophoba* meta JGIcombinedJ30088_10000835 1 o | L | L | N | A | G | A | D | A | L | V | E | G | G | K | S | G | I | F | T | P | M | F | F | F | L |
| | | *Aplysina aerophoba* meta Ga0209021_10001462 meta | T | L | E | L | C | A | A | A | L | A | E | A | G | R | L | G | I | F | T | P | M | Y | F | I | H |
| | | **Aplysina aerophoba* meta JGIcombinedJ30088_1000330310 | M | L | N | L | A | G | T | A | F | A | E | A | G | R | L | G | I | F | S | P | M | Y | F | I | L |
| | (2) | **Aplysina aerophoba* meta Ga0209754_100026449 | F | L | N | V | A | G | E | A | F | A | E | A | G | R | L | G | I | F | S | P | M | Y | F | L | L |
| | | *Petrosia ficiformis* meta LXNJ01000389 | T | L | N | L | C | A | A | A | M | A | E | A | G | R | L | G | I | F | T | P | M | Y | F | I | H |
| | | **Aplysina aerophoba* meta Ga0209544_100039960 | K | L | N | L | C | A | A | A | M | A | E | A | G | R | L | G | I | F | T | P | M | Y | F | L | H |
| | | *Nitrospira* sp., coral reef MAG 1 of 2 2682229274 | L | L | N | R | G | A | D | A | L | V | S | G | G | E | S | G | I | F | T | P | M | F | F | F | L |
| | | *Sandaracinus* sp., marine MAG 1 of 2 2775843484 | F | L | H | A | G | A | D | A | L | V | E | G | G | E | S | G | I | F | T | P | M | F | W | F | V |
| | | Freshwater, Lake Fryxell, Antarctica meta | F | L | N | R | G | A | D | A | L | V | R | G | G | E | L | G | I | F | T | P | M | A | F | F | L |
| | | Western Arctic Ocean meta Ga0133547_104202862 | L | L | N | R | A | A | E | G | L | V | T | G | G | E | L | G | I | F | T | P | M | Y | L | V | V |
| | (1) | **Aplysina aerophoba* meta JGIcombinedJ30088_100127566 | I | L | N | T | A | A | A | A | L | A | E | S | G | R | L | R | I | F | T | P | M | Y | F | V | L |
| | | Chlamydiae sp., drinking water MAG 2 of 2 2619789261 | F | L | N | M | G | A | D | A | F | V | E | A | G | K | R | K | L | F | T | P | S | L | F | F | L |
| | | *Sandaracinus* sp., marine MAG 2 of 2 2775843485 | F | L | N | Q | G | A | D | A | L | V | A | G | G | K | T | G | T | F | T | P | M | Y | F | F | L |
| | | Marine sediment, Gulf of Thailand meta | L | L | I | Q | T | A | D | A | L | V | D | A | G | R | L | G | I | F | T | P | M | Y | Y | F | R |
| | | Spirochaetes sp., soil MAG 2709604172 | L | L | I | K | T | A | D | Y | L | V | E | A | G | K | K | G | I | F | T | P | M | F | Y | F | R |
| | | Hotspring sediment, Dewer Creek, BC meta | L | L | R | L | A | G | D | P | I | A | R | A | G | R | A | G | I | F | T | P | M | L | F | F | H |
| | | Lake sediment, Walker Lake, Nevada meta | L | L | N | A | G | A | D | A | L | I | A | A | G | E | T | G | I | F | T | P | M | F | Y | V | R |
| | | Marine sediment, Helgoland, North Sea meta | M | L | G | R | A | A | E | G | L | I | A | G | G | E | T | G | I | F | T | P | S | F | F | F | L |
| | | *Sarcotragus foetidus* meta LXNI01004480 | L | L | N | T | A | A | D | A | L | V | E | G | G | R | T | G | I | F | T | P | M | F | F | F | L |
| | | *Theonella swinhoei* associated Chromatiales sp. MAG 1 of 2 | L | L | N | R | G | A | D | A | L | V | A | G | G | E | S | G | L | F | T | P | A | F | F | F | L |
| | | *Petrosia ficiformis* meta LXNJ01003202 | F | L | N | W | G | A | D | T | L | V | R | G | G | Q | T | G | I | F | T | P | M | F | F | T | L |
| | | *Petrosia ficiformis* meta LXNJ01014139 | I | L | N | A | G | A | D | A | L | V | D | G | G | R | T | G | V | F | T | P | M | F | Y | F | L |
| No activity | | *Aplysina aerophoba* meta JGIcombinedJ30088_10000835 2 o | L | L | D | R | T | A | D | A | Y | V | A | G | G | K | S | G | I | F | T | P | L | Y | C | F | L |
| | | *Petrosia ficiformis* meta LXNJ01004423 | L | M | D | R | T | A | D | A | Y | V | A | G | G | K | T | G | V | F | T | P | L | Y | C | F | L |
| | | *Theonella swinhoei* associated Chromatiales sp. MAG 2 of 2 | L | M | D | R | T | A | E | A | Y | I | A | G | G | K | T | G | I | F | T | P | L | Y | C | F | L |
| | | Hotspring, Beatty, Nevada meta_Ga0114945_100210811 | F | L | N | A | A | A | A | A | L | A | R | G | G | E | T | G | V | F | T | P | S | F | Y | F | L |
| | | *Nitrospira* sp., coral reef MAG 2 of 2 2682229270 | L | M | D | R | T | A | E | A | Y | V | A | G | G | K | S | G | L | F | T | P | L | Y | C | F | L |
| | | **Aplysina aerophoba* meta JGIcombinedJ30088_100153964 | M | L | S | L | A | A | A | A | F | G | E | A | G | R | L | G | I | F | T | P | M | Y | F | I | H |
| | | **Aplysina aerophoba meta* JGIcombinedJ30088_100077196 | F | L | L | E | S | A | K | V | W | A | E | S | G | R | L | G | I | L | S | P | M | Y | F | V | L |
| | | **Aplysina aerophoba meta* JGIcombinedJ30088_100209763 | L | L | N | L | C | A | T | A | M | V | A | S | G | R | L | G | I | F | T | P | M | Y | F | L | H |
| | | **Aplysina aerophoba meta* JGIcombinedJ30088_100057164 | L | L | N | M | A | A | A | A | F | A | E | S | G | Q | L | G | I | F | T | P | M | Y | F | V | L |
| | | **Sarcotragus foetidus* meta LXNI01006544 | L | L | N | K | S | A | D | A | L | V | E | A | G | R | L | G | I | F | T | P | C | Y | F | V | H |
| | | **Sarcotragus foetidus* meta LXNI01016685 | L | L | N | L | C | A | T | A | M | V | A | A | G | R | L | G | I | F | T | P | M | Y | L | V | H |
| | | Consensus | L | L | N | L | A | A | D | A | L | V | E | G | G | R | L | G | I | F | T | P | M | Y | F | F | L |
| | | **Consensus #** | 426 | 427 | 428 | 429 | 430 | 431 | 432 | 433 | 434 | 435 | 436 | 437 | 438 | 439 | 440 | 441 | 442 | 443 | 444 | 445 | 446 | 447 | 448 | 449 | 450 |

| | | *S. cerevisiae* # | 365 | 366 | 367 | 368 | 369 | 370 | 371 | 372 | 373 | 374 | 375 | 376 | 377 | 378 | 379 | 380 | 381 | 382 | 383 |
|---|---|---|---|---|---|---|---|---|---|---|---|---|---|---|---|---|---|---|---|---|---|
| Eukaryotic | (1) | *Saccharomyces cerevisiae* | A | R | K | P | E | N | A | E | T | P | S | Q | T | S | Q | E | A | T | Q |
| | | *Trypanosoma brucei* | A | R | K | P | R | L | G | E | E | L | S | C | - | - | - | - | - | - | - |
| | | *Arabidopsis thaliana* SMT1 | A | R | K | P | E | - | - | - | - | - | - | - | - | - | - | - | - | - | - |
| | (2) | *Arabidopsis thaliana* SMT2 | C | R | K | P | E | S | P | E | E | S | S | - | - | - | - | - | - | - | - |
| | | *Glycine max* SMT1 | A | R | K | P | D | S | D | R | N | - | - | - | - | - | - | - | - | - | - |
| | | *Chlamydomonas reinhardtii* | F | R | K | P | G | A | D | K | K | K | - | - | - | - | - | - | - | - | - |
| | | *Pneumocystis jirovecii* | C | R | K | P | L | V | - | - | - | - | - | - | - | - | - | - | - | - | - |
| Bacterial | (3) | Chlamydiae sp., drinking water MAG 1 of2 | V | M | R | P | - | - | - | - | - | - | - | - | - | - | - | - | - | - | - |
| | | *Aplysina aerophoba* meta JGIcombinedJ30088_10000835 1 of | A | R | K | P | R | R | S | E | D | - | - | - | - | - | - | - | - | - | - |
| | | *Aplysina aerophoba* meta Ga0209021_10001462 meta | A | R | K | P | E | - | - | - | - | - | - | - | - | - | - | - | - | - | - |
| | | **Aplysina aerophoba* meta JGIcombinedJ30088_1000330310 | A | R | K | P | D | - | - | - | - | - | - | - | - | - | - | - | - | - | - |
| | (2) | **Aplysina aerophoba* meta Ga0209754_100026449 | G | R | K | P | E | - | - | - | - | - | - | - | - | - | - | - | - | - | - |
| | | *Petrosia ficiformis* meta LXNJ01000389 | A | R | K | P | D | K | H | E | H | K | - | - | - | - | - | - | - | - | - |
| | | **Aplysina aerophoba* meta Ga0209544_100039960 | G | R | K | P | E | - | - | - | - | - | - | - | - | - | - | - | - | - | - |
| | | *Nitrospira* sp., coral reef MAG 1 of2 2682229274 | A | R | K | P | G | A | R | E | T | P | - | - | - | - | - | - | - | - | - |
| | | *Sandaracinus* sp., marine MAG 1 of2 2775843484 | A | E | R | P | A | - | - | - | - | - | - | - | - | - | - | - | - | - | - |
| | | Freshwater, Lake Fryxell, Antarctica meta | A | Q | K | P | L | - | - | - | - | - | - | - | - | - | - | - | - | - | - |
| | | Western Arctic Ocean meta Ga0133547_104202862 | A | R | K | P | K | G | - | - | - | - | - | - | - | - | - | - | - | - | - |
| | (1) | **Aplysina aerophoba* meta JGIcombinedJ30088_100127566 | V | R | K | P | A | - | - | - | - | - | - | - | - | - | - | - | - | - | - |
| | | Chlamydiae sp., drinking water MAG 2 of2 2619789261 | V | M | R | P | - | - | - | - | - | - | - | - | - | - | - | - | - | - | - |
| | | *Sandaracinus* sp., marine MAG 2 of2 2775843485 | V | R | K | P | E | A | - | - | - | - | - | - | - | - | - | - | - | - | - |
| | | Marine sediment, Gulf of Thailand meta | A | R | K | V | A | P | R | A | - | - | - | - | - | - | - | - | - | - | - |
| | | Spirochaetes sp., soil MAG 2709604172 | A | K | K | - | - | - | - | - | - | - | - | - | - | - | - | - | - | - | L |
| | | Hotspring sediment, Dewer Creek, BC meta | A | R | K | G | A | R | - | - | - | - | - | - | - | - | - | - | - | - | - |
| | | Lake sediment, Walker Lake, Nevada meta | L | R | K | P | K | - | - | - | - | - | - | - | - | - | - | - | - | - | - |
| | | Marine sediment, Helgoland, North Sea meta | A | R | R | K | - | - | - | - | - | - | - | - | - | - | - | - | - | - | - |
| | | *Sarcotragus foetidus* meta LXNI01004480 | A | R | K | P | Q | H | P | G | D | - | - | - | - | - | - | - | - | - | - |
| | | *Theonella swinhoei* associated Chromatiales sp. MAG 1 of2 | A | R | K | P | E | N | R | G | T | E | - | - | - | - | - | - | - | - | - |
| | | *Petrosia ficiformis* meta LXNJ01003202 | A | R | K | P | L | R | S | T | D | - | - | - | - | - | - | - | - | - | - |
| | | *Petrosia ficiformis* meta LXNJ01014139 | A | R | K | P | A | R | D | - | - | - | - | - | - | - | - | - | - | - | - |
| | No activity | *Aplysina aerophoba* meta JGIcombinedJ30088_10000835 2 of | A | R | K | P | L | - | - | - | - | - | - | - | - | - | - | - | - | - | - |
| | | *Petrosia ficiformis* meta LXNJ01004423 | A | R | K | P | E | - | - | - | - | - | - | - | - | - | - | - | - | - | - |
| | | *Theonella swinhoei* associated Chromatiales sp. MAG 2 of2 | A | Q | K | P | L | - | - | - | - | - | - | - | - | - | - | - | - | - | - |
| | | Hotspring, Beatty, Nevada meta_Ga0114945_100210811 | A | R | K | P | A | - | - | - | - | - | - | - | - | - | - | - | - | - | - |
| | | *Nitrospira* sp., coral reef MAG 2 of2 2682229270 | A | R | K | P | F | - | - | - | - | - | - | - | - | - | - | - | - | - | - |
| | | **Aplysina aerophoba* meta JGIcombinedJ30088_100153964 | A | R | K | P | D | - | - | - | - | - | - | - | - | - | - | - | - | - | - |
| | | **Aplysina aerophoba meta* JGIcombinedJ30088_100077196 | A | R | K | P | L | S | A | R | E | - | - | - | - | - | - | - | - | - | - |
| | | **Aplysina aerophoba meta* JGIcombinedJ30088_100209763 | A | R | K | P | R | - | - | - | - | - | - | - | - | - | - | - | - | - | - |
| | | **Aplysina aerophoba meta* JGIcombinedJ30088_100057164 | A | R | K | P | A | N | C | - | - | - | - | - | - | - | - | - | - | - | - |
| | | **Sarcotragus foetidus* meta LXNI01006544 | V | R | R | P | R | - | - | - | - | - | - | - | - | - | - | - | - | - | - |
| | | **Sarcotragus foetidus* meta LXNI01016685 | A | R | K | P | K | - | - | - | - | - | - | - | - | - | - | - | - | - | - |
| | | Consensus | A | R | K | P | E | - | - | - | - | - | - | - | - | - | - | - | - | - | - |
| | | **Consensus #** | **451** | **452** | **453** | **454** | **455** | **456** | **457** | **458** | **459** | **460** | **461** | **462** | **463** | **464** | **465** | **466** | **467** | **468** | **469** |

**Supplementary Figure 1. Full MAFFT-linsi alignment of SMT protein sequences. Positions are numbered to the *S. cerevisiae* SMT sequence (top) and the consensus sequence (bottom).** Numbers in parenthesis denote the number of methylations performed by the SMTs. Bolded residues indicate those conserved among functional SMTs. Boxed residues indicate sites subjected to site-directed mutagenesis in this study. Residues shaded in yellow indicate those within 5 Å of 24-methylenecholesterol docked into the Chl_ SMT ColabFold predicted structure or other SMT predicted structures aligned to that SMT. Residues shaded in magenta indicate those within 5 Å of SAM. Residues shaded in orange indicate those within 5 Å of both 24-methylencecholsterol and SAM. Mutants of the positions shaded in cyan were previously found to permit the *S. cerevisiae* SMT to perform a second methylation. Red text indicates residues in nonfunctional SMTs that deviate from those conserved among functional SMTs. *SMT homologs new to this study.
